# Discovery and mechanism of action of small molecule inhibitors of ceramidases

**DOI:** 10.1101/2021.06.15.448479

**Authors:** Robert D. Healey, Essa M. Saied, Xiaojing Cong, Gergely Karsai, Ludovic Gabellier, Julie Saint-Paul, Elise Del Nero, Sylvain Jeannot, Marion Drapeau, Simon Fontanel, Damien Maurel, Shibom Basu, Cedric Leyrat, Guillaume Bossis, Cherine Bechara, Thorsten Hornemann, Christoph Arenz, Sebastien Granier

**Author notes:** These authors contributed equally (Robert D. Healey, Essa M. Saied).

## Abstract

Sphingolipid metabolism is tightly controlled by enzymes to regulate essential processes such as energy utilisation and cell proliferation. The central metabolite is ceramide, a pro-apoptotic lipid catabolized by ceramidase enzymes to ultimately produce pro-proliferative sphingosine-1-phosphate. Human ceramidases can be soluble proteins (acid and neutral ceramidase) or integral membrane proteins (alkaline ceramidases). Increasing ceramide levels to increase apoptosis has shown efficacy as a cancer treatment using small molecules inhibiting a soluble ceramidase. Due to the transmembrane nature of alkaline ceramidases, no specific small molecule inhibitors have been reported. Here, we report novel fluorescent substrates (FRETceramides) of ceramidases that can be used to monitor enzyme activity in real-time. We use FRETceramides to discover the first drug-like inhibitors of alkaline ceramidase 3 (ACER3) which are active in cell-based assays. Biophysical characterization of enzyme:inhibitor interactions reveal a new paradigm for inhibition of lipid metabolising enzymes with non-lipidic small molecules.

**Table of contents summary:** Use of synthetic fluorescent ceramide molecules allows the discovery of the first selective drug-like small molecule inhibitors for alkaline ceramidase 3, an intra-membrane enzyme involved in sphingolipid metabolism in health and disease.

## Introduction

Dysregulation of lipid metabolism is a hallmark of many cancers and metabolic diseases. However the inherent complexities of the lipidome make studying lipid metabolism challenging. The sphingolipid family of bioactive lipids are key regulators of metabolic physiology, with at least 33 enzymes directly involved in sphingolipid metabolism^1–3^. Ceramides are biomolecules at the core of sphingolipid biology regulating critical cell membrane properties and signalling^4^. The molecular control of ceramide levels as a target area for pharmacological intervention has matured significantly in recent years^5–7^. Many enzymes involved in sphingolipid metabolism are integral membrane proteins, however small molecule inhibitors targeting specifically these classes of enzymes remain to be developed.

“Could ceramides become the new cholesterol?”^8^ This recent essay clearly reflects the novel paradigm that ceramides are critical lipids regulating human physiology, contributing in particular to insulin resistance and metabolic diseases. It also highlights how urgent it is to investigate this exciting class of bioactive sphingolipids. Beyond their role as metabolic messengers, ceramides are strongly connected with cancer signalling as they are regulators of cell fate (i.e. proliferation vs apoptosis)^7, 9, 10^. In the nervous system, ceramides are of fundamental importance in neuronal function as the precursors of most sphingolipids critically participate in myelin sheath formation and maintenance^11^. Indeed, defects in ceramide metabolism can lead to neurological dysfunction and severe clinical conditions^12, 13^.

Ceramidases are hydrolase enzymes which cleave ceramides to produce fatty acids and bioactive lipid sphingosine. The five ceramidase genes are commonly further categorized by pH required for optimal activity acid ceramidase (aCDase, ASAH1) a soluble enzyme, neutral ceramidase (nCDase, ASAH2) a membrane-anchored enzyme and alkaline ceramidases (ACER1, ACER2, ACER3) integral membrane proteins. Recent reports have also suggested other ceramidases exist, such as the adiponectin receptors or related members^14, 15^. These enzymes are crucial for sphingolipid homeostasis^16^ however since they share a common substrate, ceramides, developing selective drug-like inhibitors has been challenging, with the exception of acid ceramidase which possesses a unique cysteine-based hydrolase activity^17^.

Alkaline ceramidase 3 (ACER3) is essential during early human development, as a single point mutation (E33G) causes enzyme inactivation, leading to progressive leukodystrophy in patients^18^. Recent structural data provided the molecular basis of ACER3 dysfunction^19^: it possesses a 7-transmembrane alpha-helical structure, architecturally similar to an inverted GPCR^20^, the active site comprises a coordinated Zinc ion which is allosterically linked to a calcium binding site, catalytically linked to activity and disrupted by the E33G genetic variant. Ceramide homeostasis has been directly linked to cancer progression^7^ and ACER3 overexpression has recently been highlighted in various cancers^21, 22^. Furthermore, ACER3 has been shown to play a role in progression of metabolic associated fatty liver disease^23^. In order to further validate ACER3 as a therapeutic target in any capacity, selective, potent and cell active inhibitors are required. A major hurdle in developing ACER3 inhibitors is a lack of a simple, robust assay to monitor activity.

Here we describe the development of a FRET based-assay utilising chemically engineered ceramides named FRETceramides amenable to high-throughput activity screening of ceramidases (ACER3, nCDase and aCDase). We use this assay to discover small molecule inhibitors targeting an intramembrane ceramidase, ACER3. We perform computationally guided ligand optimization and characterize the optimized inhibitor biophysically. We propose a novel paradigm for developing ceramidase inhibitors by exploiting enzyme conformational dynamics which can be applied to other classes of enzymes with shared substrates. Finally, cell-based bioassays validate this inhibitor as the first selective, non-toxic inhibitor of ACER3 active in cells. Together with our recent structural studies^14, 19^ the data presented here provide structural dynamic insights into the mechanism by which intramembrane ceramidases hydrolyse ceramide to produce bioactive lipid sphingosine. In addition, these are the first tool compounds reported for this emerging class of enzyme and can be used to gain new insights into the sphingolipid metabolic signalling network.

## Results

### FRETceramides can directly monitor ceramidase activity in real-time

In extension of our previously developed concept using fluorophore-modified synthetic substrates to monitor enzymatic function in real-time^24–26^, we designed and synthesised a suite of doubly fluorophore-modified ceramides (Fig. 1) as substrates for ceramidases. Coumarin and nitrobenzoxadiazole (NBD) were utilised as donor and acceptor fluorophores due to compatible excitation/emission maxima for FRET and relatively small sizes, therefore minimally impacting on substrate recognition by the enzymes. We tested substrate recognition and hydrolysis using five ceramidases: neutral ceramidase (nCDase), acid ceramidase (aCDase), and the three alkaline ceramidases (ACER1, ACER2, ACER3, Fig. 1). Intramembrane ACER3 was able to hydrolyse the synthetic substrate FRETcer1. Membrane-anchored enzyme nCDase displayed a promiscuous activity, hydrolysing two synthetic substrates, FRETcer2 and FRETcer3. No synthetic substrates were hydrolysed by soluble aCDase nor intramembrane ACER1 and ACER2 (Fig. 1 and Supplementary Fig. 1). The high signal-noise ratio displayed by the synthetic FRETceramide substrates upon hydrolysis by ACER3 or nCDase displays the amenability of these substrates as excellent tools to perform activity assays and ligand screening. The present study focuses on ligand discovery for ACER3.

**Fig. 1.**
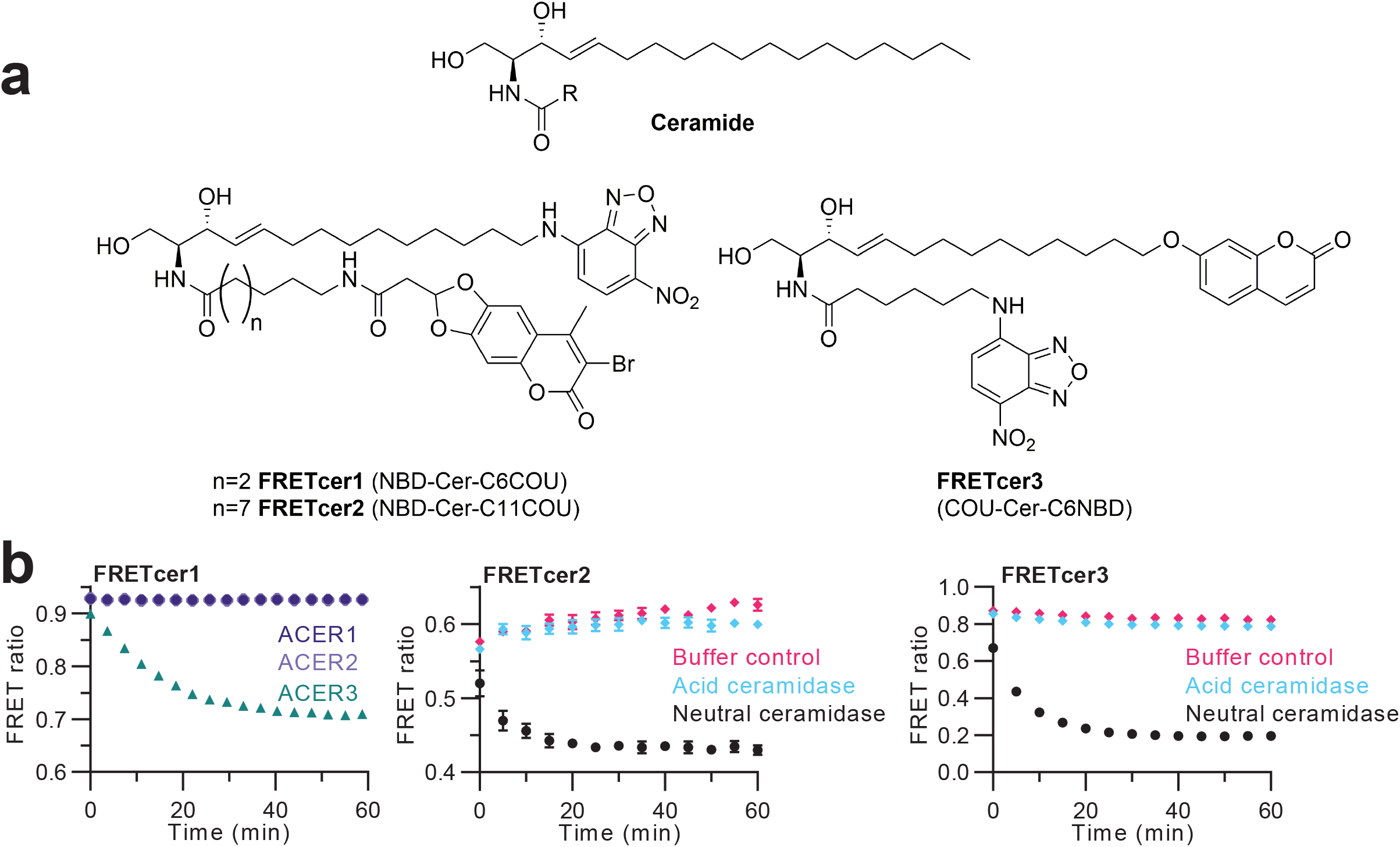
FRETceramides are tools to measure activity of ceramidases in real-time. **a**, Chemical structure of ceramide (R represents a saturated or unsaturated acyl chain) and FRETceramides (**FRETcer1**, **FRETcer2**, **FRETcer3**). **b**, Real-time analysis of ceramidase activity measured by hydrolysis of FRETceramides. ACER3 activity can be measured using **FRETcer1** and neutral ceramidase activity can be monitored using **FRETcer2** and **FRETcer3**. No hydrolysis by ACER1, ACER2 or acid ceramidase was observed for these synthetic substrates. Experiments were performed on three independent protein preparations, in quadruplicates, data is representative of a single experiment, error bars represent standard deviation.

### Characterization of FRETcer1

In order to validate FRETcer1 as a synthetic substrate for ACER3, we monitored the change in fluorescence when the detergent-purified enzyme was incubated with the substrate. Although the coumarin (donor) emission drastically increased as the reaction proceeded, minimal change in NBD (acceptor) emission was observed (Supplementary Fig. 2). This is due to the local environment sensitivity exhibited by the fluorophore NBD^27^ which, when conjugated to sphingosine, will remain in the hydrophobic ACER3:detergent micelle complex following substrate cleavage. The observed Michaelis-Menten kinetics of ACER3 with FRETcer1 showed a Km of 10.76 ± 2.03 µM (Supplementary Fig. 2). We confirmed that the change in fluorescence was indeed due to cleavage of the probe using LC-MS (Supplementary Fig. 2).

Ceramide hydrolysis by ceramidases produces two products, sphingosine and free fatty acid. ACER3 has an intramembrane Zinc-based active site cavity enveloped by seven transmembrane alpha helices^19^. Ceramide substrates partially enter the cavity from the lipid bilayer to be hydrolysed by ACER3 with the fatty acid moiety of ceramide completely inside the cavity whilst the sphingosine moiety rests closer to the lipid bilayer. This mechanism is consistent with the observed substrate preference exhibited by ACER3 for unsaturated (fatty acid moiety) ceramide species over their saturated counterparts in microsome-based assays^28^; however direct hydrolysis of saturated ceramides by ACER3 has also been observed *in vitro*^19^. To validate the synthetic FRETcer1 substrate for inhibitor discovery, we explored the inhibitory effect of endogenous ACER3 lipidic products (fatty acids and sphingosine) on enzymatic activity. All the free fatty acids tested showed inhibitory activity, and unsaturated fatty acids were more potent inhibitors than their saturated counterparts (Supplementary Fig. 2). Furthermore, sphingosine was the least effective lipidic inhibitor of ACER3 activity, only displaying partial inhibition of the enzyme activity (Supplementary Fig. 2). This indicates that ACER3 enzymatic activity may be controlled by a product auto-inhibition mechanism regulated at the enzyme level by fatty acid concentrations, rather than sphingosine levels. Since typical ceramide hydrolysis is measured using an LC-MS based assay to quantify sphingosine^29^, the auto-inhibitory effects of these enzymatic products have until now been unknown.

### HTS with FRETceramide reveals non-lipidic ACER3 inhibitors

Targeting the active site of ACER3 with a non-lipidic, drug-like small molecule could be challenging given the lipidic nature of the endogenous substrate, ceramide. We initiated an inhibitor-finding program targeting ACER3, by miniaturising a FRET-based screening assay utilising FRETcer1 as substrate. We performed two screens, the first using the commercially available LOPAC library containing 1280 small molecules and the second using a subset of the Enamine Discovery Diversity library containing 10,658 unique small molecules (Fig. 2). In total we have screened 11,938 small molecules across 50 assay plates, the Z’ was calculated for each plate and ranged between 0.5 - 0.9. The screening assay was performed using purified ACER3 solubilised in L-MNG detergent and compounds were tested at a final concentration of 10 µM (Fig. 2). Screening identified 98 hits; compounds which elicited an inhibition response between 40-120%. These initial hits were ordered as fresh powders and subjected to dose-response experiments, resulting in 11 primary hits active on purified protein (Fig. 2).

**Fig. 2.**
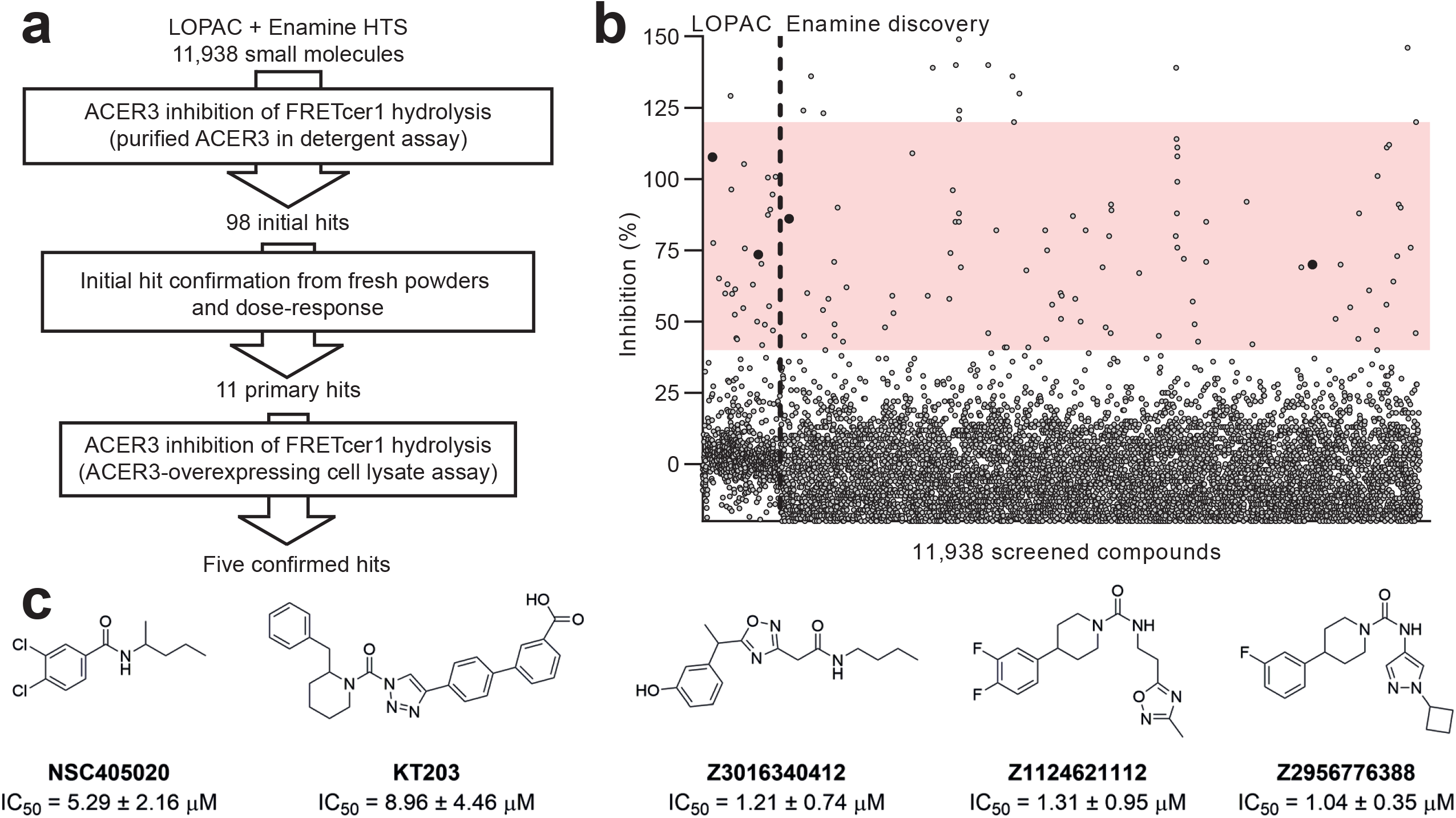
HTS ACER3 inhibitor discovery and hit confirmation. **a**, The workflow undertaken in this study. **b**, Screening results from LOPAC library (10 µM compound concentration) and a subset of the Enamine discovery library (20 µM compound concentration). Assays performed using purified ACER3 in L-MNG detergent micelles and **FRETcer1** as substrate. Confirmed hit compounds are depicted as black filled circles. **c**, Molecular structures of the five confirmed hits from two scaffolds, amides: **NSC405020** and **Z3016340412** and piperidine ureas: **KT203**, **Z1124621112** and **Z2956776388**. IC_50_ values were calculated from at least three independent dose-response experiments using **FRETcer1** as a substrate for ACER3 activity in cell lysates.

A secondary assay was developed to validate hits in a setting of higher biological complexity. Crude membranes containing many proteins, lipids and other cellular components were prepared from *Spodoptera frugiperda* Sf9 cells overexpressing ACER3 and activity was monitored by FRET using FRETcer1 as substrate. This secondary assay revealed that in the presence of lipids and nonspecific proteins, some compounds no longer displayed ACER3 inhibition probably due to binding other membrane proteins or interactions with lipids. Following dose-response experiments and calculations of potency in the cell lysate assay (Supplementary Fig. 3), five compounds were identified as confirmed inhibitors (Fig. 2). We classified the ACER3 inhibitors into two distinct scaffolds: amide-based inhibitors (NSC405020 and Z3016340412) and piperidine urea-based inhibitors (KT203, Z1124621112 and Z2956776388). We tested all confirmed hits for inhibitory activity against other ceramidases. Piperidine urea-based inhibitor KT203 displayed moderate inhibitory action against neutral ceramidase, with an IC_50_ of 19.54 ± 3.18 µM (Supplementary Fig. 3). The observed nCDase activity of KT203 indicates this compound may serve as a scaffold for the development of a new generation of nCDase inhibitors. No acid ceramidase inhibition was detected by confirmed ACER3 inhibitors (Supplementary Fig. 3).

### Optimization of ACER3 inhibitors

We selected the amide-scaffold compound NSC405020 and performed a ligand optimization study. NSC405020 has no reported cytotoxicity and has been used *in vivo* as an apparent PEX inhibitor in the millimolar range without significant cytotoxic effects^30–32^. We first confirmed NSC405020 inhibited hydrolysis of the endogenous substrate, CerC18:1 using an LC-MS assay. This assay revealed that NSC405020 does inhibit hydrolysis of CerC18:1, but also that ACER3 is able to hydrolyse the inhibitor, producing dichlorobenzoic acid (Supplementary Fig. 4). NSC405020 is thus a competitive substrate for ACER3. Based on this finding, we designed and synthesized a set of NSC405020 analogues, optimizing the scaffold to create a non-hydrolysable NSC405020 analogue with potent ACER3 inhibitory activity. The activity of the amide-modified analogues (ES_ACR01-07, Supplementary Fig. 5) revealed that some structural features around the amide-scaffold are essential for inhibitory activity; ester, sulfonamide and urea analogues displayed no inhibitory activity. Interestingly, reversing the amide position from benzamide to anilide produced an active inhibitor, Compound 01 (ES_ACR03) which was not hydrolysed by ACER3 (Supplementary Fig. 5).

To further guide ligand optimization efforts, a computational approach combining docking and replica-exchange molecular dynamics (MD) simulations was employed. This approach revealed a binding site for ACER3 inhibitors inside the transmembrane region of ACER3, and explained how ACER3 is able to hydrolyse NSC405020 (Fig. 3). The carbonyl oxygen of NSC405020 binds ACER3 in the active site, forming a polar contact with ACER3-coordinated Zinc ion and allowing ACER3 to process the inhibitor as a substrate (Fig. 3, Supplementary Fig. 4). Compound 01 is not processed by ACER3 and interacts with the enzyme higher in the cavity and away from the active site, forming a carbonyl hydrogen bond with ACER3 at Tyr149 on transmembrane helix 5 (TM5, Fig. 3). Based on the computational results, Compound 01 was used as a lead-scaffold, and a set of aryl-substituted analogues were designed, synthesised and characterized (Supplementary Fig. 5). By introducing a *m*-trifluomethyl substitution in place of *m*-chloro, Compound 02 (ES_ACR13) displayed a significant increase in potency of one-log unit to IC_50_ 0.70 µM (Fig. 3). Docking and simulation of Compound 02 revealed hydrophobic interactions with ACER3 at Leu70 of TM2 and Trp220 on TM7. The *m*-trifluoromethyl substitution on Compound 02 causes a torsional rotation of the aryl allowing a π-π interaction with Tyr149 on TM5 that was not observed with Compound 01 in addition to the carbonyl amide hydrogen bond. Furthermore, a potential halogen interaction between fluorine and Gln146 is expected although poorly defined in the forcefield used in our computations (Fig. 3).

**Fig. 3.**
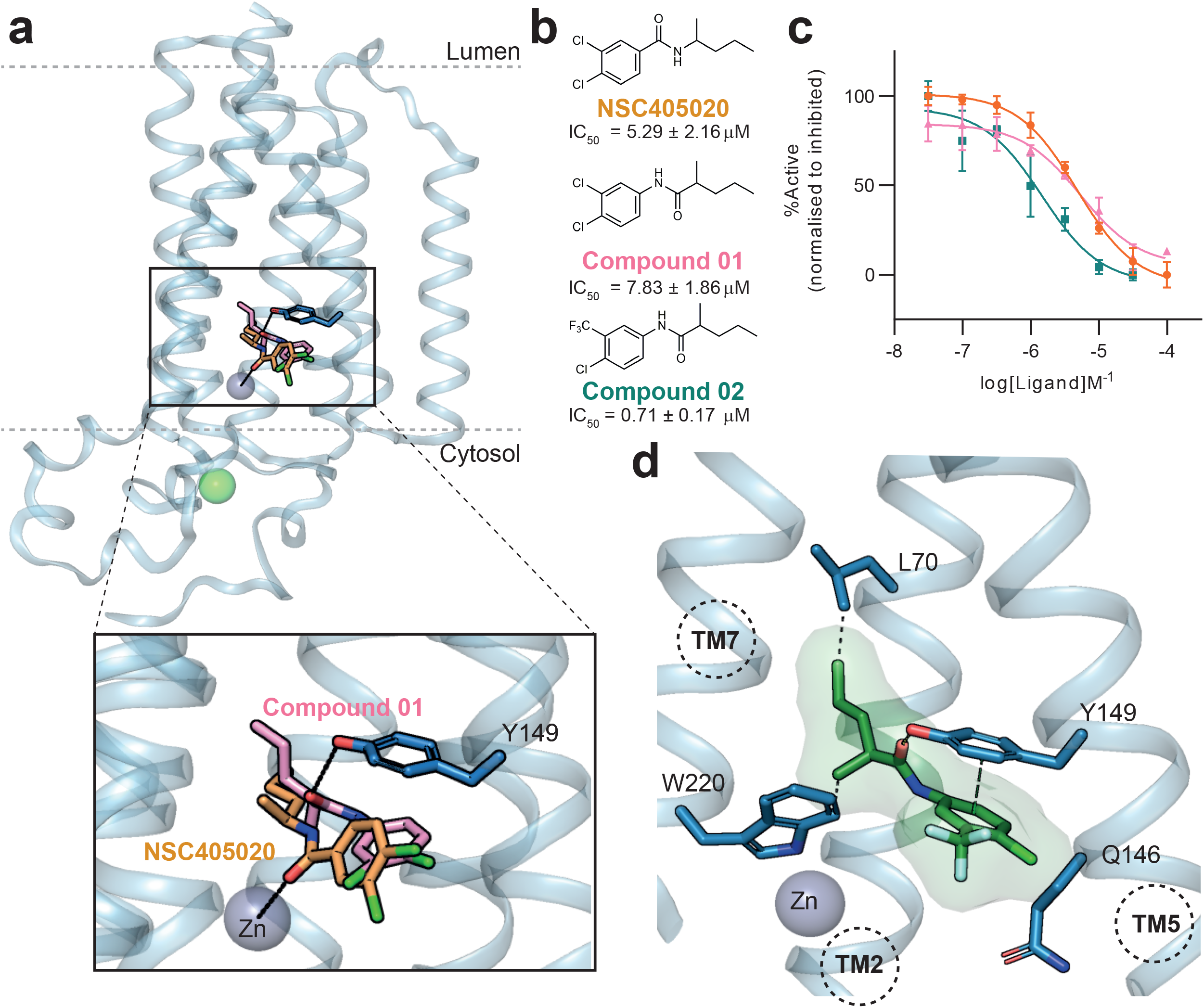
Optimisation of ACER3 inhibitors. **a**, Overall structure of intramembrane inhibitor binding site with **NSC405020** and **Compound 01** overlaid in orange and pink, respectively. Inset: carbonyl oxygen of **NSC405020** interacts with Zinc whereas carbonyl oxygen of **Compound 01** hydrogen bonds with Tyr149. **b**, Dose-response curves and **c**, structures of **NSC405020** analogues. IC_50_ values were calculated from at least three independent dose-response experiments using **FRETcer1** as a substrate for ACER3 activity in cell lysates. Experiments were performed on three independent preparations in quadruplicates, data is representative of a single experiment, error bars represent standard deviation. IC_50_ values were calculated as mean from all experiments. **d**, Close-up view of docked and relaxed pose for **Compound 02** binding site. **Compound 02** makes hydrophobic interactions with transmembrane helices 2, 5 and 7 of ACER3 and a hydrogen bond with Tyr149 on TM5.

We next explored modifications at the acyl moiety of the Compound 01 scaffold (Supplementary Fig. 5). These efforts led to Compound 03 (ES_ACR15) which exhibited an increase in potency of one-log unit to IC_50_ 0.71 µM (Supplementary Fig. 5). Surprisingly, when we combined the optimized aryl-substitution moiety and acyl moiety, Compound 04 (ES_ACR27), the compound no longer displayed ACER3 inhibitory activity (Supplementary Fig. 5). The loss of ACER3 activity for Compound 04 was likely due to both substitutions increasing the inhibitor size resulting in a conformation too large to fit inside the transmembrane cavity of ACER3. Nevertheless, Compound 04 was tested for ceramidase specificity and displayed some inhibitory effects on neutral ceramidase activity, Compound 04 could thus serve as a starting point for medicinal chemistry optimisation as an inhibitor of neutral ceramidase.

### ACER3 dynamics control hydrolase activity

To explore ACER3 conformational changes induced by Compound 02 and for binding site validation, we used hydrogen-deuterium exchange coupled to mass spectrometry analysis (HDX-MS, ^33, 34^. We incubated ACER3 purified in L-MNG detergent micelles with vehicle (DMSO) or Compound 02 and followed the differential deuteration (ΔHDX) of ACER3 in order to monitor regions that shows variation of solvent accessibility (protected or unprotected) in the presence of the inhibitor. HDX-MS experiments were performed in triplicates for three different biological samples following recommendations^35^, and no correction of back exchange was applied since we are interested in differential and not absolute HDX. Common coverage maps from ACER3 peptides identified with enzyme alone and in the presence of Compound 02 showed a sequence coverage >78 % for all three biological replicates (Supplementary Fig. 6A and Supplemental Data 1). However, the total number of identified ACER3 peptides was always significantly higher in the presence of Compound 02 compared to the enzyme alone. This indicates that binding of Compound 02 is somehow facilitating the enzymatic digestion of ACER3 by pepsin.

Differential fractional uptake heat maps showed mainly statistically significant protected regions in ACER3 in the presence of Compound 02 (Fig. 4 and Supplementary Fig. 6). Interestingly, peptides from the predicted binding region 135-148 encompassing TM5 were protected from exchange in the presence of Compound 02 in all biological replicates (Fig. 4), further validating the calculated binding pose. HDX-MS also revealed an effect on ACER3 regions TM1 and ECL1 that were more protected in the presence of Compound 02 (Fig. 4). As these regions were not predicted to directly interact with the inhibitor, we hypothesize that binding of Compound 02 induces ACER3 allosteric conformational changes away from its binding site. No significant effect of Compound 02 on deuterium uptake was observed in other regions such as ECL1 (Fig. 4). Taken together the HDX-MS data revealed the putative Compound 02 binding site and its effect on ACER3 conformations.

**Fig. 4.**
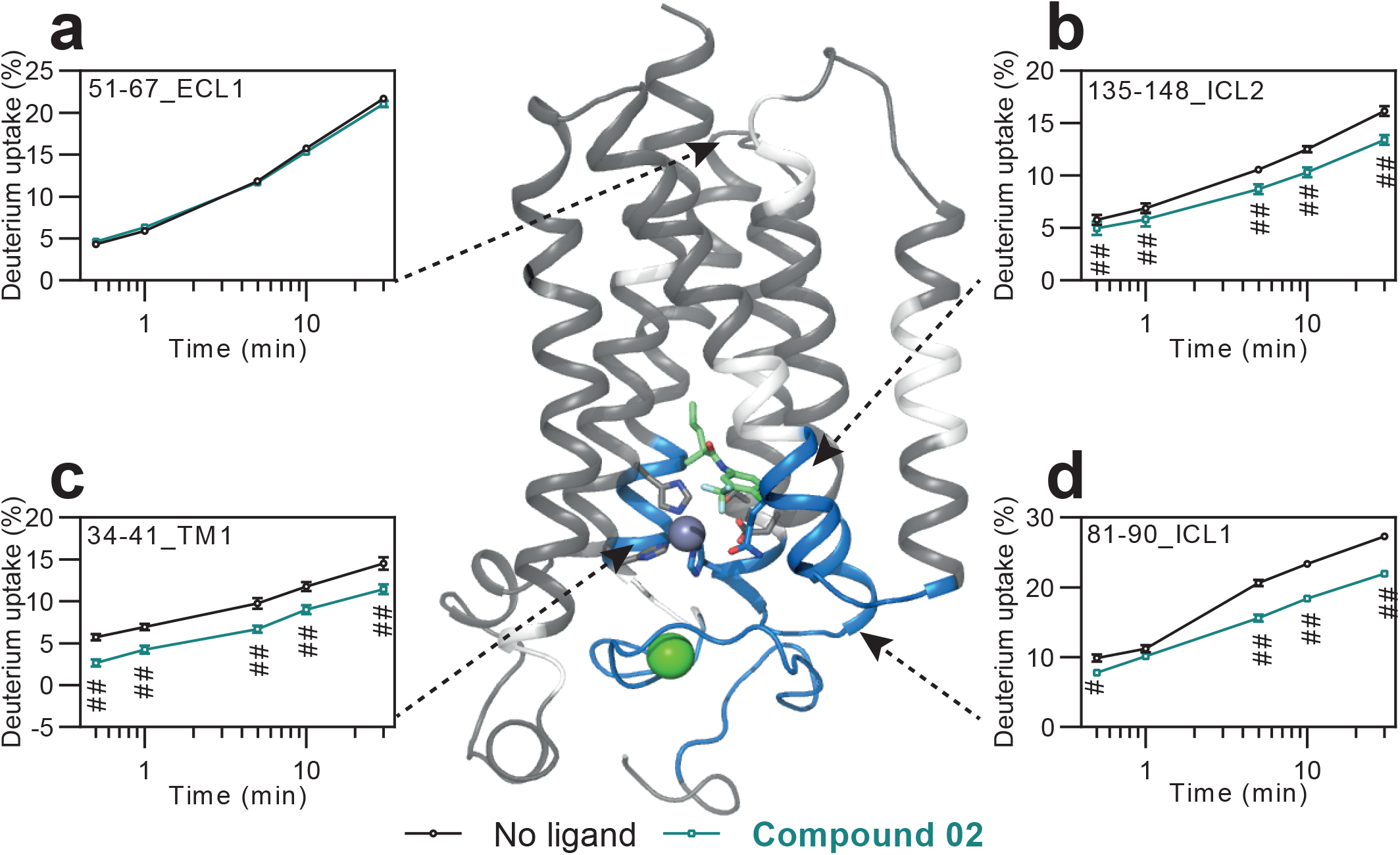
HDX-MS results of ACER3 with small molecule inhibitor, Compound 02. An ACER3 model depicted as a cartoon with ΔHDX data reflecting all tested biological replicates mapped by colour. ΔHDX-protected regions are shown in blue, regions with no significant ΔHDX in gray and regions with no sequence coverage in white. **Compound 02** is shown as green sticks, interacting residues covered by HDX and active-site residues are shown as sticks. **a**, The ECL1 region on ACER3 wild-type does not show any statistically significant ΔHDX in the presence of **Compound 02**. **b**, The ICL2 region (135-148) contains binding residues for **Compound 02** and displays a protected ΔHDX in the presence of **Compound 02**. **c**, ICL1 and **d**, TM1, are ΔHDX protected regions in the presence of **Compound 02**, although these regions do not contain residues in direct contact with **Compound 02**. Experiments were performed on three independent biological samples in triplicates and presented regions were consistent between all. Peptide traces shown are representative of a single experiment. Statistically significant using hybrid test in Deuteros 2.0, ^#^ > 99%, ^##^ > 99.9%.

To explore conformational changes of ACER3 induced by Compound 02, we performed replica-exchange molecular dynamics simulations^36^. The system was prepared using the ACER3:Compound 02 complex, generated for ligand optimization, inserted into a phosphatidylcholine (POPC) lipid bilayer. In simulations without ligand, we observed oscillatory movements of TM4 and TM6 demonstrating the dynamic nature of these regions (Fig. 5a). In the presence of an endogenous substrate, CerC18:1, TM4 and TM6 were stabilised and TM5 around ICL2 displayed an oscillatory motion between opening and closing (Fig. 5b). For ceramide hydrolysis to occur, the ceramide molecule must enter into the active site of ACER3, most likely via the lipid bilayer. In the crystal structure, the active site is visible to the lipid bilayer via an opening between TM5 and TM6^19^. Exploring the binding site of ceramide reveals a hook-shaped cavity accommodating the lipid acyl chain (Fig. 5e). To characterize these dynamic regions during simulation, we defined distance measurements at Zinc-Lys138 and Zinc-Ile143, measuring TM4 and TM5 movements, respectively and a TM5-TM6 angle to illustrate the opening or closure of TM5 (Fig. 5d).

**Fig. 5.**
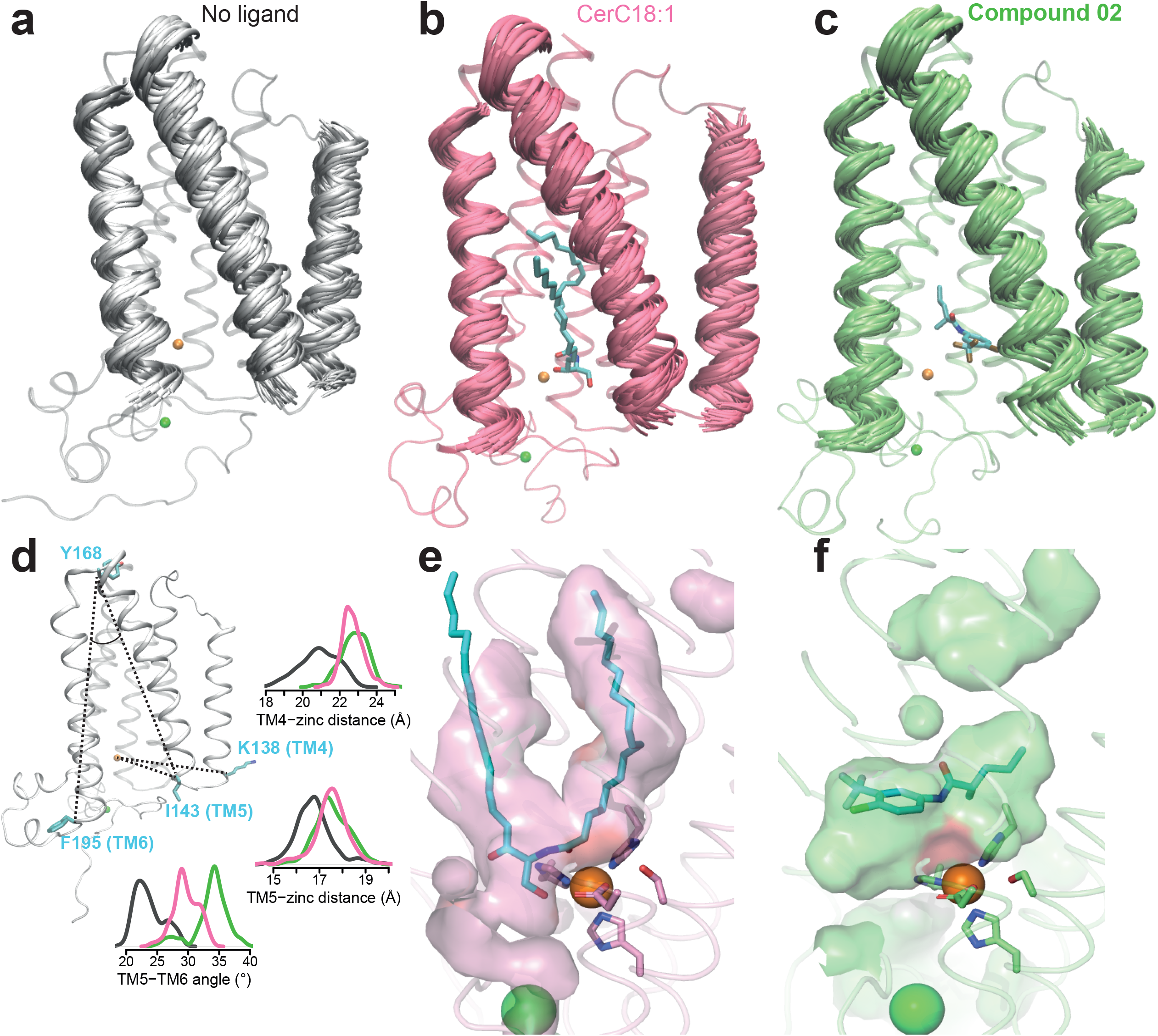
MD simulations reveal distinct conformational states of ACER3 with substrate or inhibitor. **a**, Molecular dynamics ensembles of ACER3 with no ligand reveal that ACER3 is an intrinsically dynamic enzyme. **b**, MD ensembles of ACER3-CerC18:1 complex show a stabilisation on the intracellular sides of TM4 and TM6, whilst TM5 continues to oscillate. **c**, MD ensembles of ACER3-Compound 02 complex display an opening of TM4, TM5 and TM6. Highly flexible loop regions are shown as a single conformation for clarity. **d**, Measurements of angles and distances from MD trajectories. **e**, The binding site of CerC18:1 is a continuous hook-shaped cavity within the transmembrane core of ACER3. **f**, Compound 02 binding to ACER3 results in allosteric rearrangement inside the transmembrane bundle, resulting in the formation of two distinct cavities.

Conversely, in the presence of Compound 02 TM4 and TM6 translate into stabilised, outward conformations (Fig. 5c, d). TM5 also translated outward, via the cytosolic loop (ICL2) linking TM4 and TM5, resulting in an opening and stabilisation of the TM5-TM6 cavity entrance. Compound 02 sits inside the entrance of the cavity, the TM5-TM6 interface, blocking the path from the lipid bilayer to the active site (Fig. 5). The conformational changes of ACER3 induced by binding of Compound 02 leads to an overall rearrangement of the transmembrane, breaking the ceramide binding cavity (Fig. 5f). Taken together, HDX-MS and MD simulations suggest that Compound 02 inhibits ACER3 hydrolysis activity by a dual mechanism involving 1) blocking substrate entry into the active site and 2) stabilising a single open-conformation of ACER3 with an allosteric rearrangement of the ceramide binding site.

### Compound 02 inhibits ACER3 in cells

Prior to testing **Compound 02** in cells, we confirmed that the inhibitor was selective for ACER3 against other relevant ceramidases. **Compound 02** showed at least >1000-fold selectivity for inhibition of ACER3 against both acid and neutral ceramidases (Supplementary Fig. 7). To test inhibition of ACER3 by **Compound 02** in living cells, we generated two cell lines in FreeStyle 293-F cells, inducibly overexpressing either ACER3-wild type (ACER3-wt) or ACER3-E33G mutant, a disease-causing mutation which renders ACER3 inactive^18, 19^. The E33G mutant cell line was used as a negative control for non-specific activity of **Compound 02** in the absence of active ACER3. Similar expression levels of wild type and mutant ACER3 were observed in each cell line (Fig. 6). These cell lines were treated with **Compound 02** or vehicle for 24 hours and quantitative sphingolipidomic analysis was performed. In ACER3-wt cells, **Compound 02** significantly increased the abundant long-chain ceramides, with up to a 50% increase in the most affected ceramide species (CerC16:0 and CerC18:0) whereas very long chain ceramides were not changed (Fig. 6). In the ACER3-E33G cell line, CerC16:0 was detected at 63.1 ± 7.9 pmoles/million cells whereas in the ACER3-wt cells was 29.9 ± 1.0 pmoles/million cells (Fig. 6). When ACER3-wt cells were treated with **Compound 02**, CerC16:0 was elevated to 53.2 ± 3.2 pmoles/million cells, whereas when ACER3-E33G cells were treated with **Compound 02**, CerC16:0 was unchanged compared to vehicle. Taken together, those experiments reproducibly demonstrate that **Compound 02** engages directly with ACER3 resulting in modulation of ceramide levels in live cells (Fig. 6).

**Fig. 6.**
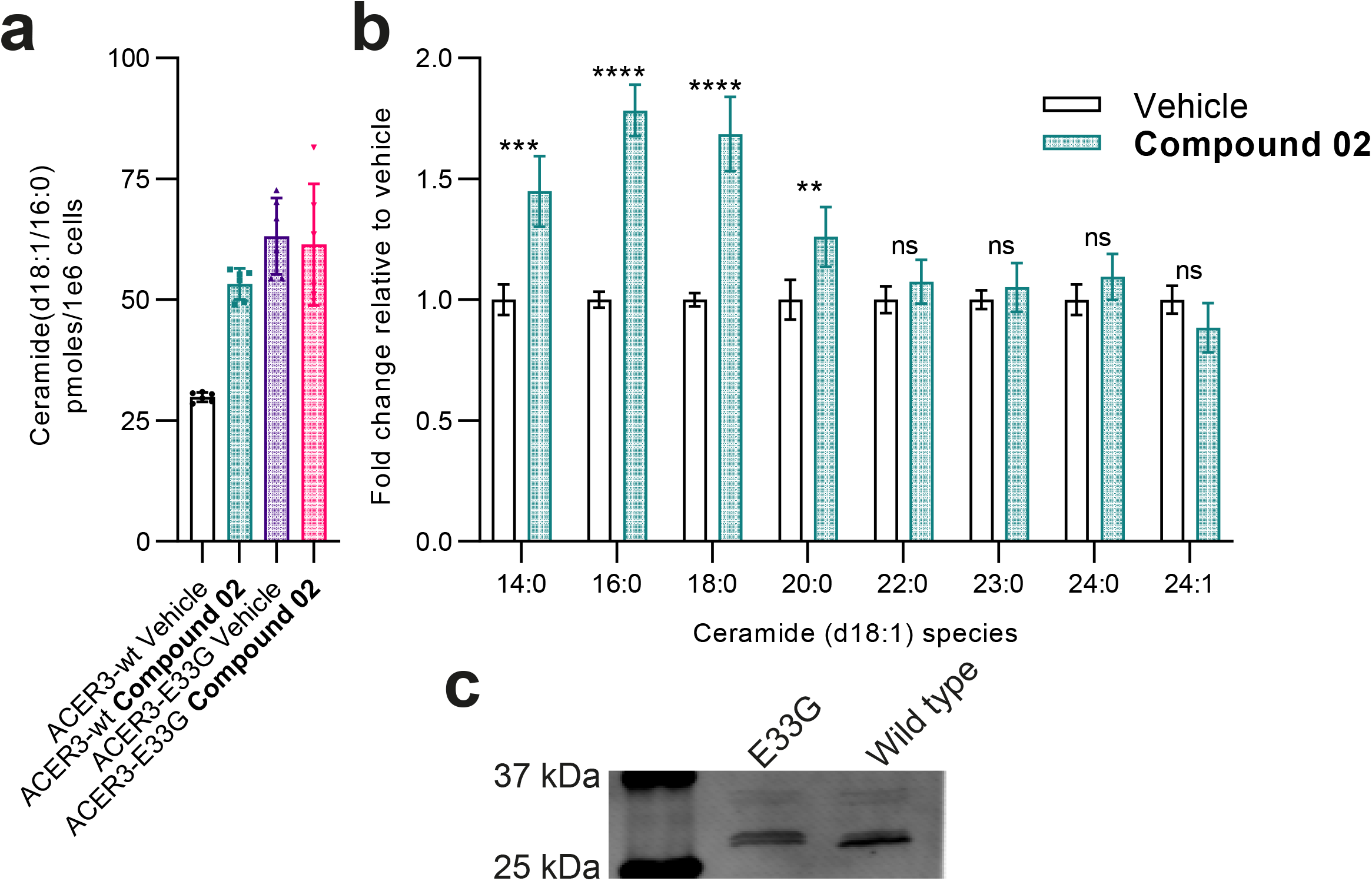
Compound 02 inhibits ACER3 in live cells. FreeStyle293-F cells expressing ACER3-wt or ACER3-E33G were treated with 12.9 µM **Compound 02** or vehicle (0.0129% DMSO) in culture medium for 24 hours. Sphingolipids were extracted and quantified by LC-MS. Two independent experiments were performed with six technical replicates, the same trend of sphingolipid species modulation was observed in both experiments. Data is representative of a single experiment and plotted as mean ± SD. Asterisks indicate statistical significance over vehicle, ** p < 0.005, *** p < 0.0005, **** p < 0.0001. **a**, Ceramide(d18:1/16:0) quantification in cells expressing ACER3-wt or ACER3-E33G treated with **Compound 02**. No effect of **Compound 02** is observed in ACER3-E33G expressing cells, indicating that **Compound 02** is a specific ACER3 inhibitor and engages the target in live cells. **b**, **Compound 02** treatment for 24 hours in cells overexpressing ACER3 alters long-chain ceramide levels. **c**, Western blot reveals similar expression overexpression levels between ACER3-wt and ACER3-E33G. Detection: ACER3 (anti-Flag) in 24 hour doxycycline induced FreeStyle293-F cells.

To validate **Compound 02** as an inhibitor of ACER3 in endogenously expressing cells, we also treated and analysed sphingolipids in U937 cells^21^. In these cells, **Compound 02** significantly increased ceramide and hexosyl-ceramide levels whilst simultaneously decreasing deoxy-ceramide levels (Supplementary Fig. 8). A comprehensive sphingolipid analysis of U937 cells revealed most ceramide(d18:1) species were elevated, whilst dihydro-ceramide(d18:0) species were less affected (Supplementary Fig. 8). This analysis highlighted the role of ACER3 in maintaining homeostasis in sphingolipid metabolism as a large number of metabolites were modulated to some degree by treatment with **Compound 02** (Supplementary Fig. 8).

## Discussion

The therapeutic potential of manipulating ceramide levels through inhibition of sphingolipid metabolising enzymes remains to be fully exploited due to lack of selective and potent inhibitors. Some promising examples of modulating the pathway have been shown such as reducing ceramide synthesis with myriocin, an irreversible inhibitor of serine palmitoyl-transferase (SPT), which prevented or reversed atherosclerosis in mice^37^. Decreasing ceramides through SPT inhibition has also been shown to prevent or reverse insulin resistance, hepatic steatosis, type 2 diabetes, hypertension and/or cardiomyopathy^38–42^. More recently, it was shown in mice that removal of sphingolipid delta(4)-desaturase (DES1) improves metabolic homeostasis, promoting DES1 as a novel target for insulin sensitization^43^. In this work, our characterization of **Compound 02** adds a new and selective inhibitor of ACER3 to the field for further experiments in more complex biological settings and disease models.

The metabolic control of sphingolipid homeostasis has been implicated in a number of physiological processes and diseases^2^. Understanding the pathways and developing therapies requires selective inhibitor molecules. In the case of inhibiting ACER3, enzymatic inhibition has until now relied on genetic inhibition^21–23^ or lipid analogues as inhibitors^44^. This report describes the first selective inhibitor of ACER3 with drug-like properties which is active in living cells. The FRETceramides we have introduced in this work allow for direct visualisation of ceramidase activity in real-time and present a new assay methodology for studying ceramidases. Using these synthetic substrates, we have gained new insights into auto-inhibition of ACER3 regulated by fatty acid product concentrations.

We report the discovery of specific inhibitors for ACER3 from two distinct scaffolds. Using biophysical methods and computational approaches, we have optimized and characterized the mechanism of action for a small-molecule inhibitor of ACER3, **Compound 02**. Utilising MD simulations and HDX-MS, we see that **Compound 02** interaction with ACER3 changes the conformation of the enzyme, stabilising ACER3 in an open conformation. Inhibiting proteins by stabilising inactive conformations has recently been proposed as a novel drug design strategy for inhibition of TNF signalling^45^. Our work demonstrates conclusively that activity of sphingolipid metabolising enzymes, whose substrates are inherently lipidic, can be controlled using non-lipidic small molecules by controlling protein dynamics. We have demonstrated that **Compound 02** is able to engage with ACER3 in living cells, modulating levels of long-chain sphingolipids.

As ACER3 dysfunction during development is associated with severe neuropathy, we envision that **Compound 02** could modulate sphingolipid levels at different stages of development and in various tissues. Indeed, increasing ceramide levels in cancer through inhibition of acid ceramidase is an effective treatment for increasing apoptosis and resensitizing cells to chemotherapy in some cancers^46–48^. Targeting ACER3 for a similar effect has not yet been shown, though overexpression of ACER3 has been observed in some cancers ^21, 22, 49^. Furthermore ACER3 expression has been intrinsically linked to progression of nonalcoholic steatohepatitis^23^. With **Compound 02**, therapeutic opportunities for ACER3 inhibition can now be explored, for example in cancer and liver disease. In summary, this first report of a specific ACER3 inhibitor will enable deeper understanding of the effects of bioactive sphingolipid modulation in physiology and disease from a pharmacological perspective.

## Methods

### General synthetic methods

All commercially available reagents were purchased from TCI, Sigma-Aldrich, Fluka or Acros and used without further purification, unless otherwise specified.

The structure of all synthesized compounds was confirmed with ^1^H NMR, ^13^C NMR, and MS analyses. (^1^H, ^13^C) NMR spectra of the synthesized compounds were recorded on Bruker Avance-500 nuclear magnetic resonance spectrometers (^1^H at 400 or 500 MHz and ^13^C at 100 or 126 MHz) as a solution in deuterated methanol (MeOD), deuterated chloroform (CDCl_3_) or deuterated water (D_2_O) at 27°C, unless otherwise indicated. NMR spectra are reported as chemical shifts (δ) in parts per million (ppm) as referenced to the residual protons in the deuterated solvent (^1^H NMR: *d*-chloroform: δ 7.27 ppm; *d*-methanol: δ 3.31ppm; ^13^C NMR: *d*-chloroform: δ 77.16 ppm; *d*-methanol δ 49.05 ppm, unless otherwise indicated unless otherwise noted).). UPLC−MS analyses were performed on an AGILENT 6120 UPLC-MS system (Santa Clara, USA) consisting of an SQD (single quadrupole detector) mass spectrometer equipped with an electrospray ionization interface (ESI) and a photodiode array detector (DAD). The detection was performed in full scan mode and the major observable molecular ion and selected fragments and clusters have been reported.

Detailed synthetic procedures are available in **Supplementary Data 3**.

### Hydrolysis of FRETceramides by transmembrane ceramidases in cellular membranes

Stable inducible cell lines expressing each transmembrane ceramidase were prepared in homemade FreeStyle 293-F(TetR) cells using the lentiviral system^50^. Membranes from each construct were prepared from suspension cultures after two days of doxycycline-induced protein expression. Cells were lysed by dounce homogenization in lysis buffer (10 mM Tris-HCl, pH 7.5, 0.1 mM EDTA, 0.25 U/µL benzonase, protease inhibitors) and membranes isolated by centrifugation (20,000*g*). Membranes were washed with high salt buffer (50 mM HEPES, pH 7.5, 1 M NaCl, 0.25 U/µL benzonase, protease inhibitors), centrifuged and resuspended in membrane reaction buffer (50 mM HEPES, pH 7.5, 150 mM NaCl). ACER3 overexpressing insect cell membranes were also prepared for dose-response experiments. The same membrane preparation protocol was utilised for mammalian and insect expressed proteins. FRETceramidase assays were performed in 384-well black plates. Membranes (25-50 µg total protein) were mixed with different concentrations of inhibitors for 15 minutes then FRETceramides were added to 10 µM and fluorescence was monitored for 3 hours at 30 °C using 347 nm excitation with 410 nm/535 nm emissions for coumarin/NBD. FRET ratio was plotted as Em_acc_/(Em_acc_+Em_don_).

### Purification of ACER3

ACER3 fused with C-terminal BRIL (ACER3 residues 2-244 with C-terminal BRIL) and N-terminal FLAG-tag, referred to as ACER3-BRIL, was expressed by baculovirus and in *Sf9* cells and purified as described previously (ref. Nat Comm, 2018, Vasilauskaite-Brooks). Following purification, monomeric ACER3-BRIL was concentrated to 100 µM and snap-frozen in 5 µL aliquots for high-throughput screening. For all other enzymatic assays and HDX-MS experiments, wild-type ACER3 was used, no significant activity difference was observed between ACER3-BRIL and wild-type ACER3.

### ACER3 FRETceramide high-throughput screening assay

Inhibitor screening was performed using the LOPAC-1280 compound library from Sigma and a subset of the Enamine discovery diversity set from Enamine. Purified ACER3-BRIL (50-100 nM) in HTS buffer (0.002% L-MNG, 20 mM HEPES, pH 7.5, 100 mM NaCl) was incubated with compounds (10 µM) for 10 minutes at 25 °C. FRETcer1 was added to 10 µM and the reaction proceeded for 1 hour at 25 °C, fluorescence was read using 347 nm excitation and 410 nm/535 nm emissions. Oleic acid (10 µM) was used as inhibition positive control. Exclusion of compounds with intense autofluorescence was performed using raw emissions at 410 nm and 535 nm. More details are included in Supplementary Table 1.

### HDX-MS experiments

HDX-MS experiments were performed using a Synapt G2-Si HDMS coupled to nanoAQUITY UPLC with HDX Automation technology (Waters Corporation). ACER3 in L-MNG detergent was concentrated up to 10-20 µM and optimization of the sequence coverage was performed on undeuterated controls. Analysis of ACER3 with vehicle or mixed with 10 and 100 equivalent Compound 02 were performed as follows: 3 µL of sample are diluted in 57 µL of undeuterated for the reference or deuterated last wash SEC buffer. The final percentage of deuterium in the deuterated buffer was 95%. Deuteration was performed at 20°C for different times (**Supplementary Data 1** & 2). Next, 50 µL of reaction sample are quenched in 50 µL of quench buffer (50 mM KH_2_PO_4_, 50 mM K_2_HPO_4_ and 2M guanidine-HCl, pH 2.3) at 0°C. 80 µL of quenched sample are loaded onto a 50 µL loop and injected on an Enzymate pepsin column (300 Å, 5 µm, 2.1 mm X 30 mm, Waters) maintained at 15°C, with 0.2% formic acid at a flowrate of 100 µL/min and an additional backing pressure of 6000 psi controlled by the HDX regulator kit (Waters). The peptides are then trapped at 0°C on a Vanguard column (ACQUITY UPLC BEH C18 VanGuard Pre-column, 130Å, 1.7 µm, 2.1 mm X 5 mm, Waters) for 3 min, before being loaded at 40 µL/min onto an Acquity UPLC column (ACQUITY UPLC BEH C18 Column, 1.7 µm, 1 mm X 100 mm, Waters) kept at 0°C. Peptides are subsequently eluted with a linear gradient (0.2% formic acid in acetonitrile solvent at 5% up to 35% during the first 6 min, then up to 40% and 95% over 1 min each) and ionized directly by electrospray on a Synapt G2-Si mass spectrometer (Waters). HDMS^E^ data were obtained by 20-30 V trap collision energy ramp. Lock mass accuracy correction was made using a mixture of leucine enkephalin. The pepsin column was then washed three times with Guanidine-HCl 1.5 M, acetonitrile 4% and formic acid 0.8% and a blank is performed between each sample in order to minimize the carry-over. All timepoints were performed in triplicates.

Peptide identification was performed from undeuterated data using ProteinLynx global Server (PLGS, version 3.0.3, Waters). Peptides are filtered by DynamX (version 3.0, Waters) using the following parameters: minimum intensity of 1000, minimum product per amino acid of 0.2, maximum error for threshold of 5 ppm. All peptides were manually checked and data was curated using DynamX. Back exchange was not corrected since we are measuring differential HDX and not absolute one. Statistical analysis of all ΔHDX data was performed using Deuteros 2.0^51^ and only peptides with a 99% confidence interval were considered.

### Computational docking and molecular dynamics simulations

The crystal structure of ACER3 (PDB: 6G7O) was used as the initial coordinates for docking and MD. The ligands were docked to the initial ACER3 structure using Autodock Vina [10.1002/jcc.21334]. Residues in the ligand-binding pocket were set flexible during docking. The receptor-ligand complexes were embedded in a bilayer of POPC. The protonation states of titratable residues were predicted at pH 7 using the H++ server^52^. Each system was solvated in a periodic 80 × 80 × 95 Å^3^ box of explicit water and neutralized with 0.15 M of Na+ and Cl-ions. Effective point charges of the ligands were obtained by RESP fitting^53^ of the electrostatic potentials calculated with the HF/6-31G* basis set using Gaussian 09^54^. Parameters for the active-site zinc coordination were calculated with the HF/6-31G* basis set using Gaussian 09 and fitted with the MCPY program^55^. The Amber 99SB-ildn, lipid 14 and GAFF force fields were used for the proteins, the lipids and the ligands, respectively. The TIP3P and the Joung-Cheatham models were used for the water and the ions, respectively.

After energy minimization, all-atom MD simulations were carried out using Gromacs 5.1^56^ patched with the PLUMED 2.3 plugin^57^. Each system was gradually heated to 300 K and pre-equilibrated during 10 ns of brute-force MD in the NPT-ensemble. The replica exchange with solute scaling (REST2)^36^ technique was employed to enhance the sampling with 48 replicas in the NVT ensemble. The protein and the ligands were considered as “solute” in the REST2 scheme–force constants of their van der Waals, electrostatic and dihedral terms were subject to scaling. The effective temperatures used for generating the REST2 scaling factors ranged from 300 K to 700 K, following a distribution calculated with the Patriksson-van der Spoel approach (temperature predictor for parallel tempering simulations). Exchange between replicas was attempted every 1000 simulation steps. This setup resulted in an average exchange probability of ∼40%. We performed 50 ns × 48 replicas MD in the NVT ensemble for each system. The first 20 ns were discarded for equilibration. The original unscaled replica (at 300 K effective temperature) was collected and analyzed. Cluster analysis of the ligand binding pose was carried out using the Gromacs Cluster tool. The middle structure of the most populated cluster was selected as the final binding pose. Protein:inhibitor interactions were visualised using PLIP^58^ and PyMOL Molecular Graphics System (Schrödinger, LLC).

### Expression and purification of nCDase

Neutral ceramidase (nCDase) (extracellular domain residues 99-780^59^) fused with a C-terminal TwinStrep-tag and N-terminal melittin secretion sequence was expressed using the baculovirus system in *Sf9* insect cells. Secreted nCDase was purified from culture medium using Strep-Tactin resin (IBA Life Sciences) and further purified using a Superdex 200 size-exclusion chromatography column equilibrated in nCDase reaction buffer (10 mM HEPES, pH 7.5, 150 mM NaCl).

### nCDase FRETceramidase assay

nCDase (50 nM) was incubated with ACER3 inhibitors for 10 minutes at 25 °C. FRETcer3 was added to 10 µM and the reaction proceeded for 2 hours at 30 °C, fluorescence was read using 347nm excitation and 410nm/535nm emissions. Heat-inactivated nCDase was used as 0% activity control.

### Expression and purification of aCDase

aCDase (residues 22-395^60^) flanked with an N-terminal melittin secretion sequence and TwinStrep-tag was expressed using the baculovirus system in *Sf9* insect cells. Secreted aCDase was purified from culture medium using Strep-Tactin resin and further purified using a Superdex 200 size-exclusion chromatography column equilibrated in nCDase reaction buffer. Follow size-exclusion, the pH of the buffer was adjusted to 4.5 with sodium acetate (100 mM) and proteolytic autocleavage was performed at 37 °C for 2-3 days.

### aCDase ceramidase LCMS assay

aCDase (200 nM) was incubated with ACER3 inhibitors for 15 minutes 25 °C in aCDase reaction buffer (10 mM HEPES, 100 mM sodium acetate, pH 4.5, 150 mM NaCl). Ceramide-12:0 was added to 20 µM and the reaction proceeded for 1 hour at 37 °C. Reactions were quenched by adding methanol to 50% (v/v) and snap-freewing. For analysis, samples were centrifuged and sphingosine was quantified in the supernatant by UPLC-MS as described previously^14^. Heat-inactivated aCDase was used as 0% activity control.

### Generation of FreeStyle 293-F ACER3-wt and ACER3-E33G cell lines

ACER3 wild-type and E33G constructs were designed with an N-terminal Flag tag for immuno-detection followed by the protein sequence. The construct was cloned into an inducible IRES-emGFP containing vector which enabled expression induction to be quickly confirmed. The vector, pHR-CMV-TetO2_3C-Twin-Strep_IRES-EmGFP was a gift from A. Radu Aricescu (Addgene plasmid # 113884)^50^. Stably transfected cell lines expressing ACER3 after induction by doxycycline were generated using standard lentiviral transfection and transduction procedures. The transduced cell line was a homemade FreeStyle 293-F(TetR) inducible cell line, made from commercially available FreeStyle 293-F (Thermo) cells which are useful for both adherent and suspension cultures. Cells were maintained in standard cell culture medium comprising DMEM (Sigma), 10% FCS (Fetcal Calf Serum), streptomycin/penicillin antibiotics and supplemented with L-glutamine at 37°C and 5% CO2. All cells were regularly tested for mycoplasma and no contaminations were detected.

### FS293-TetR cell treatment

One day before plating and treatment, ACER3-wt or ACER3-E33G expression was induced in FreeStyle 293-F(TetR) cells with 1 µg/mL doxycycline (Sigma). Induced cells were counted and resuspended at 0.5×10^6^ cells/mL in medium containing vehicle (0.0125% DMSO) or Compound 02 (12.5 µM) and plated in 6-well plates, 2mL/well. After 24 hours, cells were harvested by trypsinization, washed with PBS twice and counted. Cell pellets were snap-frozen and stored at −20 °C prior to lipid extraction and analysis.

### U937 cell treatment

U937 cells (DSMZ) were cultured at 37°C in RPMI supplemented with 10% FBS and streptomycin/penicillin in the presence of 5% CO_2_. Cells were authenticated by the American Type Culture Collection using short tandem repeat analysis. All cells were regularly tested for mycoplasma and no contaminations were detected. For sphingolipids analysis, U937 cells were seeded at a density of 3 × 10^5^/ml and treated with vehicle (0.025% DMSO) or Compound 02 (25 μM) for 48 hours prior to harvesting. Cells were collected by centrifugation after treatment, washed with PBS and counted. Cell pellets were snap-frozen and stored at −20 °C prior to lipid extraction and analysis.

### Quantitative sphingolipidomics

Lipid extraction was performed as previously described^13^ with some modifications. Lipids were extracted by adding 1 mL of a mixture of methanol:MTBE:chloroform (MMC) 4:3:3 (v/v/v) to 0.5-2.0 million snap-frozen cells. The MMC mix was fortified with 100 pmoles/mL of the internal standards (Avanti Polar Lipids): d7-sphinganine (d18:0), d7-sphingosine (d18:1), dihydroceramide (d18:0:12:0), 1-deoxydihydroceramide (m18:0/12:0), ceramide (d18:1/12:0), 1-deoxyceramide (m18:1/12:0), glucosylceramide (d18:1/8:0), sphingomyelin (18:1/12:0) and 50 pmoles/mL d7-sphingosine-1-phosphate. After brief vortexing, the samples were continuously mixed in a Thermomixer (Eppendorf) at 37 °C, 1400 rpm for 60 min. Protein precipitation was performed by centrifugation for 5 min, 16,000*g*, 25 °C. The single-phase supernatant was collected, dried under nitrogen gas and stored at –20 °C until analysis. Before analysis, the dried lipids were dissolved in 100 µL MeOH. Liquid chromatography was performed as previously described^13^ with some modifications. The lipids were separated using a C30 Accucore LC column (ThermoFisher Scientific, 150 mm * 2.1 mm * 2.6 µm) using the following mobile phases; A) Acetonitrile:Water (6:4) with 10 mM ammonium acetate and 0.1 % formic acid, B) Isopropanol: Acetonitrile (9:1) with 10 mM ammonium acetate and 0.1 % formic acid at a flow rate of 0.26 ml/min. The following gradient was applied:

1. 0.0-2.0 min (ramp 90-80 %A, 10-20 %B),
2. 2.0-3.0 min (ramp 80-70 %A, 20-30 %B),
3. 3.0-7.0 min (ramp 70-40 %A, 30-60 %B)
4. 7.0-18.0 minutes (ramp 40-10 %A, 60-90 %B)
5. 18.0-24.0 minutes (ramp 10-0 %A, 90-100 %B)
6. 24.0-28.0 minutes (isocratic 0 %A, 100 %B)
7. 28.0-28.1 minutes (ramp 0-90 %A, 100-10 %B)
8. 28.1-30.0 minutes (isocratic 90 %A, 10 %B).

The liquid chromatography was coupled to a hybrid quadrupole-orbitrap mass spectrometer Q-Exactive (ThermoFisher Scientific); samples were analyzed in positive and negative mode using a heated electrospray ionization (HESI) interface. The following parameters were used: spray voltage 3.5 kV, Aux gas heater temperature 300 °C, sheath gas flow rate 40 AU, Aux gas 10 AU and capillary temperature of 300 °C. The detector was set to an MS2 method using a data dependent acquisition with top10 approach with stepped collision energy between 25 and 30. A 70 000 resolution was used for the full spectrum and a 17500 for MS2. A dynamic exclusion filter was applied which will excludes fragmentation of the same ions for 20 sec. Identification and Quantification was achieved as previously published^13^, with the following identification criteria:

1. Resolution with an accuracy of 5 ppm from the predicted mass at a resolving power of 140000 at 200 m/z.

2. Isotopic pattern fitting to expected isotopic distribution.

3. Matching retention time on synthetic standards if available.

4. Specific fragmentation patterns:

a) free sphingoid base: [M+H]+ à [M+H - H2O]+ and [M+H - H2O - H2O]+

b) 1-deoxysphingoid base: [M+H]+ à [M+H - H2O]+

c) S1P: [M+H]+ à [M+H - H2O]+ and [M+H - H2O - HPO4]+

d) (1-deoxy)Ceramide: [M+H]+ à [M+H - H2O]+ and [M+H - H2O - (fatty acid)]+

e) Hexosylceramide: [M+H]+ à [M+H - H2O]+ and [M+H - H2O - (fatty acid) - glycoside]+

f) Sphingomyelin: [M+H]+ à [M+H - H2O]+ and [M+H - H2O - (fatty acid) - PO4Choline]+

## Acknowledgments

This project received funding from the European Research Council (ERC) under the European Union’s Horizon 2020 research and innovation programme (grant agreement 647687). We thank the ARPEGE platform at the Institut de Génomique Fonctionnelle for providing facilities and technical support. C.A is grateful for the funding by the Deutsche Forschungsgemeinschaft DFG Grant No. AR 376/18-1. Mass spectrometry experiments were carried out using the facilities of the Montpellier Proteomics Platform (PPM, BioCampus Montpellier). This work was supported by the regional funds FEDER/Région Occitanie. This work was publicly funded through ANR (the French National Research Agency) under the “Investissements d’avenir” programme with the reference ANR-16-IDEX-0006» (University of Montpellier Isite-MUSE).

## Author contributions

R.D.H. expressed and purified proteins with help from J.S-P., S.J., M.D. & S.F. R.D.H. performed enzymatic assays with help from J.S-P. & D.M. E.M.S. & C.A. designed and synthesized the FRET ceramide probes presented in this study, performed in silico docking to structure-based design the different generations of NSC405020 inhibitor, synthesized the structural modifications of NSC405020 inhibitor, performed HPLC-MS analysis for all ceramidases experiments. C.A. supervised E.M.S and provided technical guidance. Protein-based mass spec experiments were performed by R.D.H. & E.D.N., supervised by C.B. Simulations were performed by X.C., C.L. & S.B. Cell-based assays were performed by R.D.H., M.D., L.G. & G.B. Lipidomics analysis was performed by K.G. supervised by T.H. R.D.H. & S.G. designed experiments, supervised the project and wrote the manuscript with input from all authors.

## Additional information

Raw data and summary tables for HDX-MS experiments are included as **Supplementary Data 1** and **Supplementary Data 2**, respectively. Synthetic procedures and characterization data is included as **Supplementary Data 3**.

---

**Supplementary Fig. 1.**
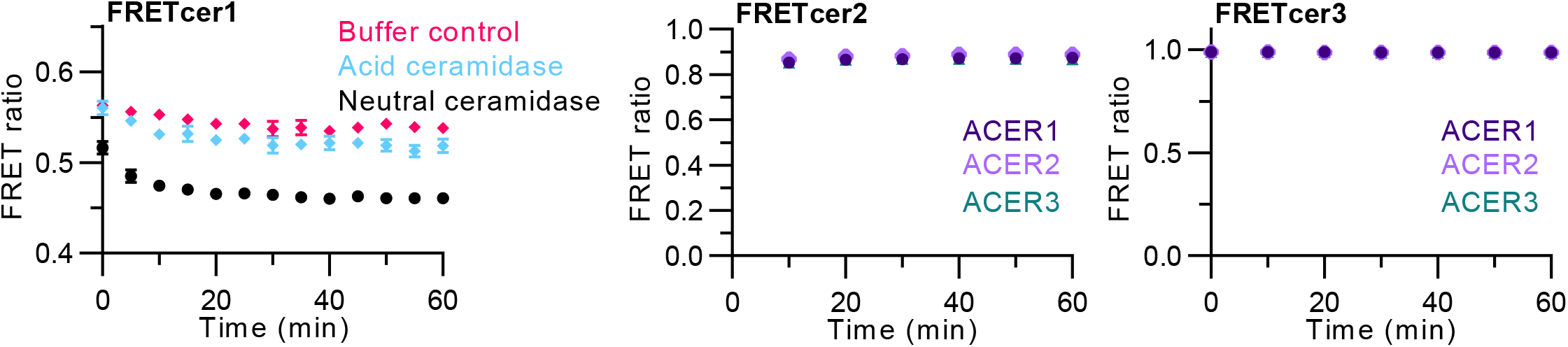
Additional activity of ceramidases on FRETceramides. **a**, Soluble ceramidases; acid ceramidase and neutral ceramidase, display minimal hydrolysis activity toward synthetic substrate **FRETcer1**. Intramembrane ceramidases ACER1, ACER2 and ACER3 display no hydrolysis activity for synthetic substrates **b** and **c**, **FRETcer2** and **FRETcer3**, respectively. Experiments were performed independently at least three times, data is representative of a single experiment, error bars represent standard deviation. Acid ceramidase activity was confirmed by an LC-MS assay monitoring hydrolysis of ceramide.

**Supplementary Fig. 2.**
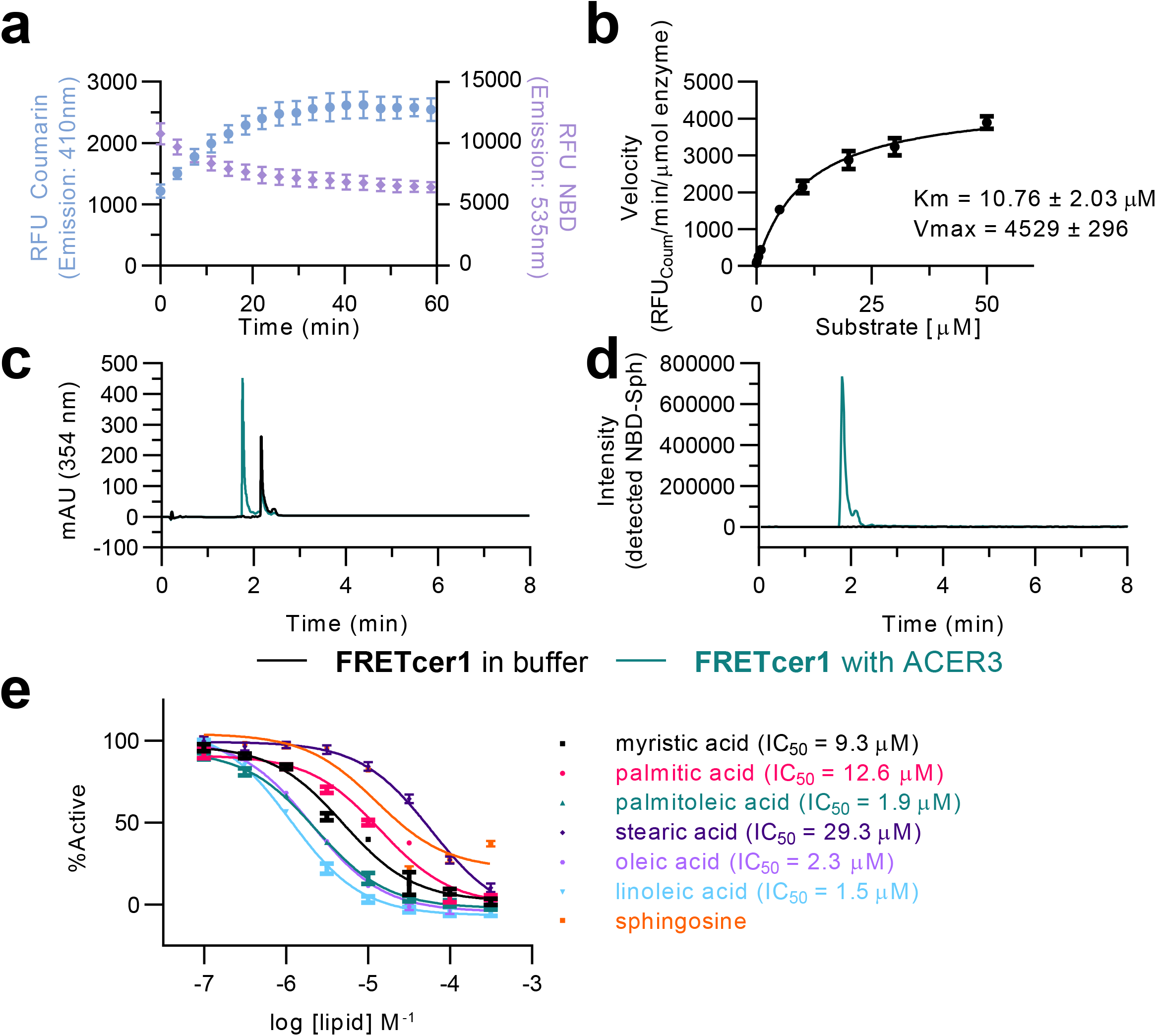
Characterization of synthetic substrate FRETcer1 hydrolysis by ACER3. **a**, Donor and acceptor emissions of **FRETcer1** when hydrolysed by ACER3 (10 µM **FRETcer1**, 0.1 µM ACER3). **b**, Michaelis-Menten analysis of ACER3 with substrate **FRETcer1**. Velocity was calculated as RFU_Coumarin_/time(minutes)/enzyme concentration(μM). FRETcer1 was confirmed as a synthetic substrate for ACER3 using LCMS observation of substrate and product, **c**, NBD absorbance is detected as a single peak for FRETcer1 alone (black), and two peaks when incubated with ACER3 (green). **d**, LCMS ion trace of NBD-sphingosine from (**c**) showing the hydrolysis product is only detected after cleavage by ACER3. **e**, ACER3 activity is inhibited by endogenously produced lipids including fatty acids and sphingosine. Unsaturated fatty acids are more potent inhibitors of ACER3 than saturated counterparts. Experiments were performed three times independently, in quadruplicate, data is representative of a single experiment, error bars represent standard deviation. IC_50_ values were calculated as mean from all experiments.

**Supplementary Fig. 3.**
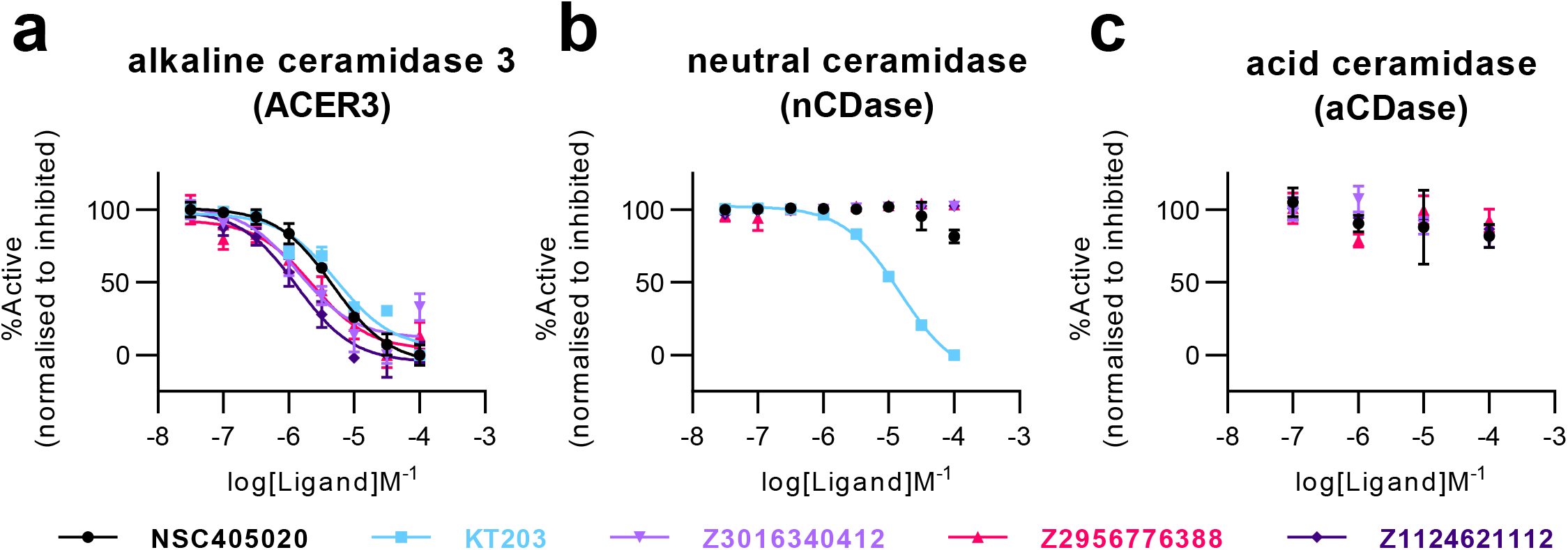
Confirmed ACER3 inhibitor screening hits inhibition of ACER3, neutral ceramidase and acid ceramide. **a**, Dose-response inhibition of ACER3. Activity of ACER3 in cell lysate assay (50 µg protein/well) was monitored by fluorescence using **FRETcer1** as substrate (10 µM). **b**, Dose-response inhibition of nCDase. **KT203** displays a non-selective inhibition of neutral ceramidase (IC_50_ 19.54 ± 3.18 µM), no inhibitory effect was seen for the other hit compounds. Activity of neutral ceramidase (0.5 µM) was monitored by fluorescence using **FRETcer3** as substrate (10 µM), heat denatured neutral ceramidase was used as inhibited control. **c**, Dose-response inhibition of acid ceramide, no inhibition of acid ceramidase was observed for any compounds. Activity of acid ceramidase (1 µM) was monitored by LC-MS using ceramideC12 (10 µM) as substrate, heat denatured acid ceramidase was used as inhibited control.

**Supplementary Fig. 4.**
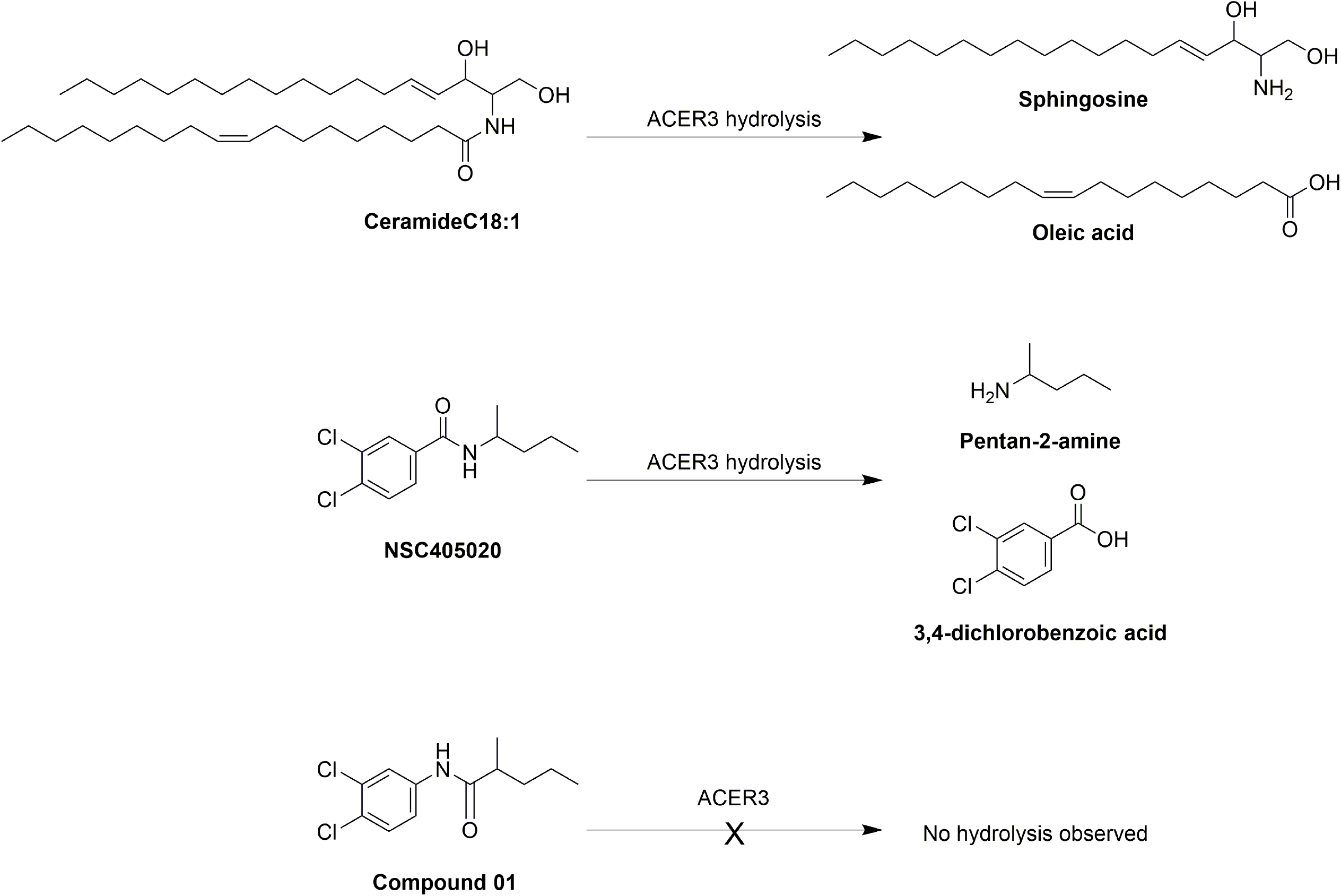
NSC405020 inhibits ACER3 hydrolysis of ceramide through competition as a substrate. Schematic of ACER3 hydrolase activity of endogenous substrate CeramideC18:1 and synthetic substrate **NSC405020**. No hydrolysis was observed for anilide containing analogues such as **Compound 01**.

**Supplementary Fig. 5.**
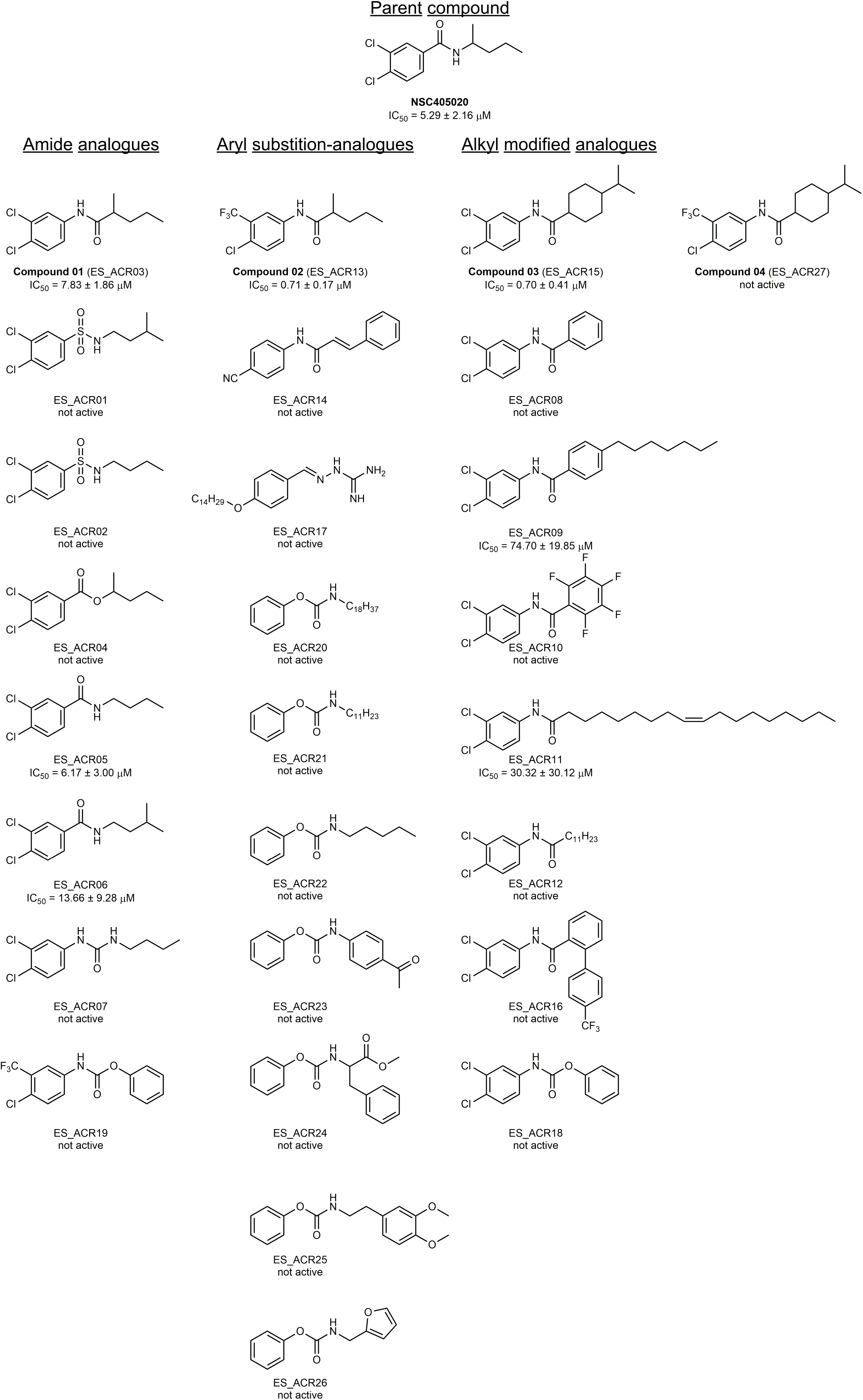
Structures of all NSC405020 SAR analogues. Reported IC_50_ values were calculated from at least three independent dose-response experiments using **FRETcer1** as a substrate for ACER3 activity in cell lysates.

**Supplementary Fig. 6.**
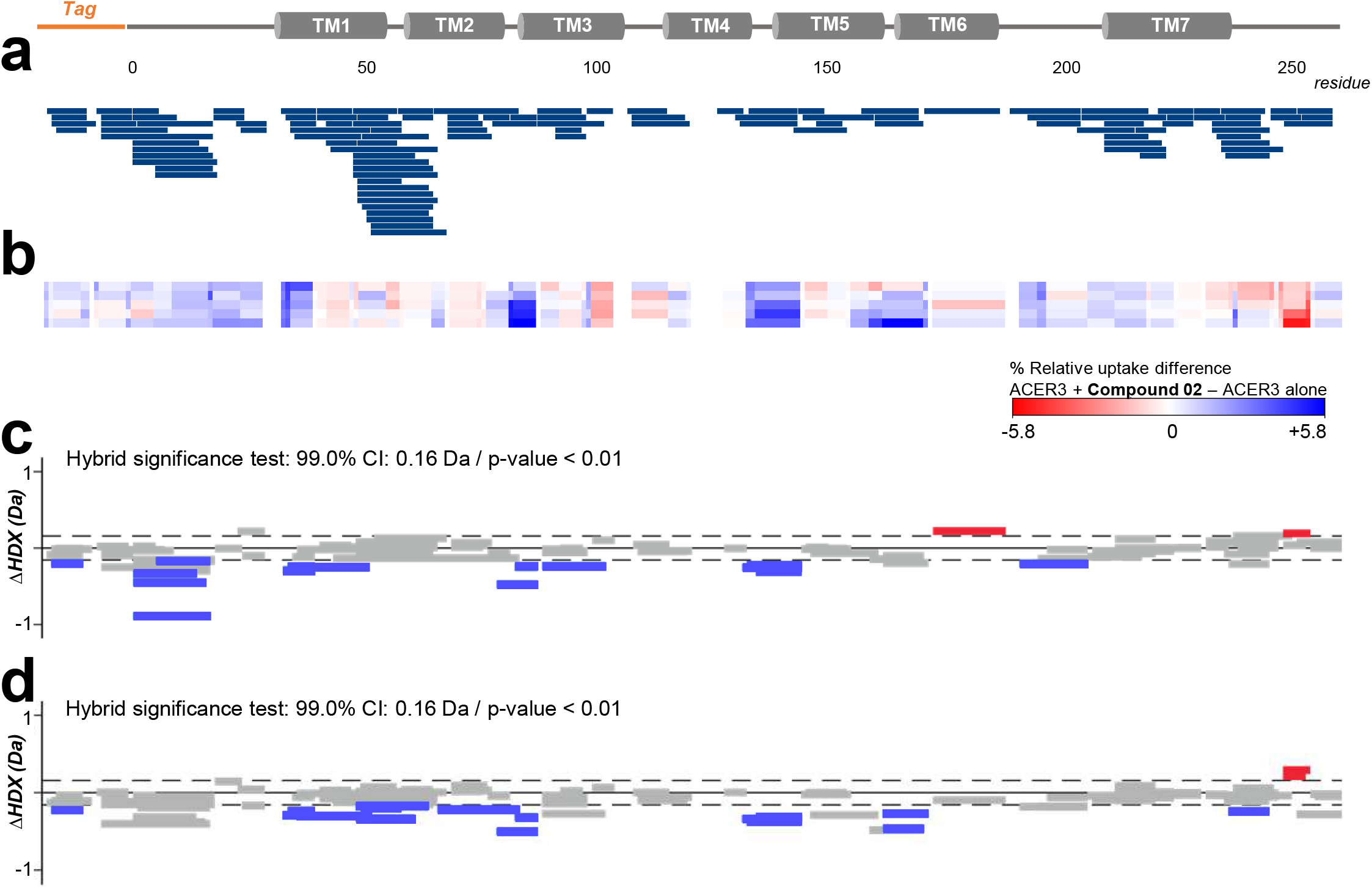
Supplementary data for HDX experiments. **a**, Common coverage map for differential HDX analysis of one biological replicate of ACER3 alone and in the presence of **Compound 02**. Number of peptides 111, sequence coverage 94.77%, redundancy 4.54. **b**, Heat map showing the relative uptake difference (ACER3 alone – ACER3 + **Compound 02**), protected and unprotected regions represented in blue red respectively. **c** and **d**, Wood’s plots comparing the relative deuterium uptake (ACER3 + **Compound 02** – ACER3 alone) after incubation in deuterated solvent for 5 min and 30 min, respectively. The dotted lines represent the 99% confidence interval, which indicates the level of difference of deuterium uptake between the two compared states is statistically significant based on a hybrid test. Red and blue coloured bars indicate statistically significant deprotected or protected peptides, respectively.

**Supplementary Fig. 7.**
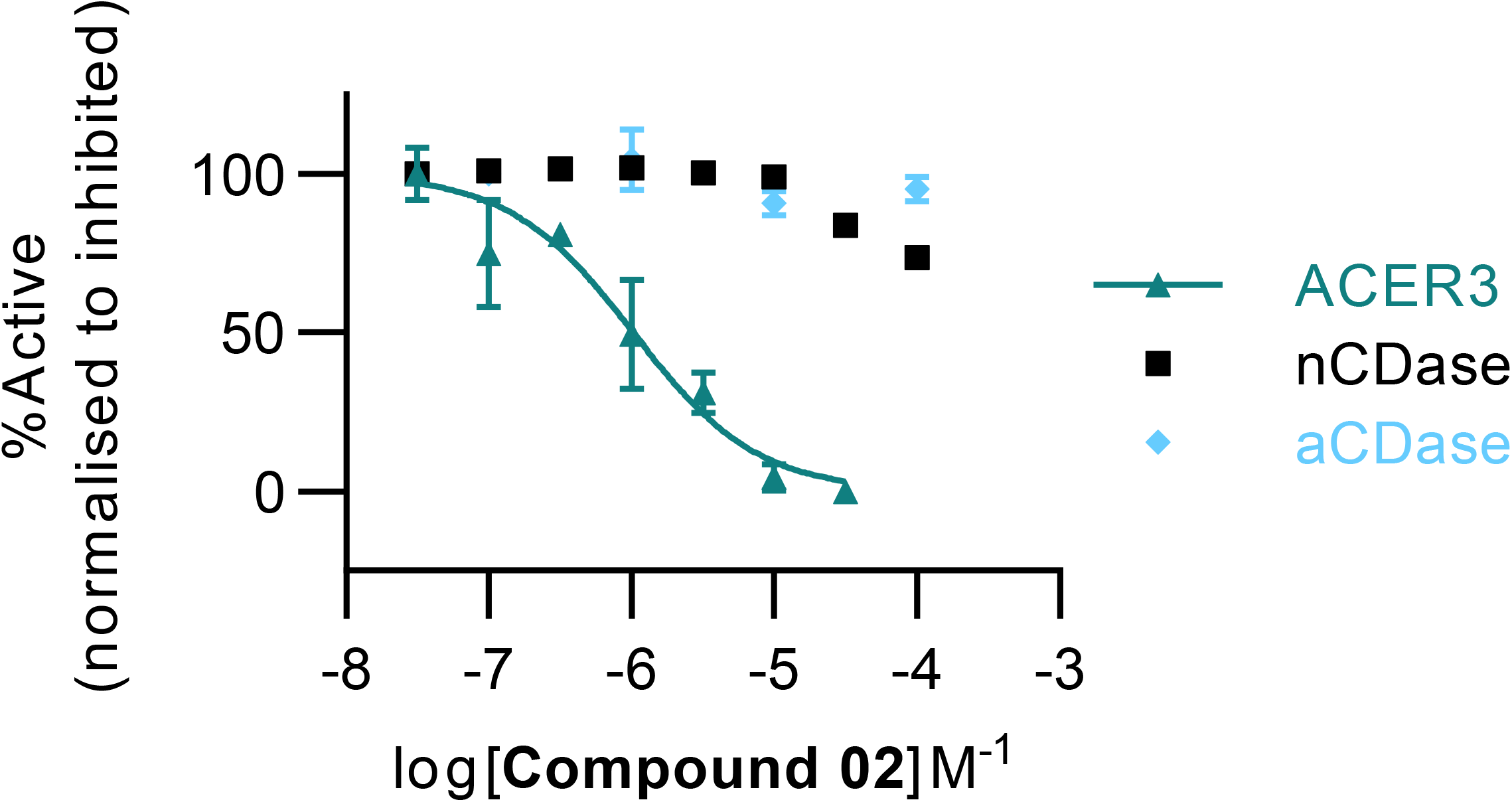
Compound 02 is a specific inhibitor for ACER3 and not other ceramidases. Activity of ACER3, neutral ceramidase and acid ceramidase in the presence of **Compound 02**. Activity of ACER3 in cell lysate assay was monitored by fluorescence using **FRETcer1** as substrate. Activity of neutral ceramidase was monitored by fluorescence using **FRETcer3** as substrate, heat denatured neutral ceramidase was used as inhibited control. Activity of acid ceramidase was monitored by LC-MS using ceramideC12 as substrate, heat denatured acid ceramidase was used as inhibited control.

**Supplementary Fig. 8.**
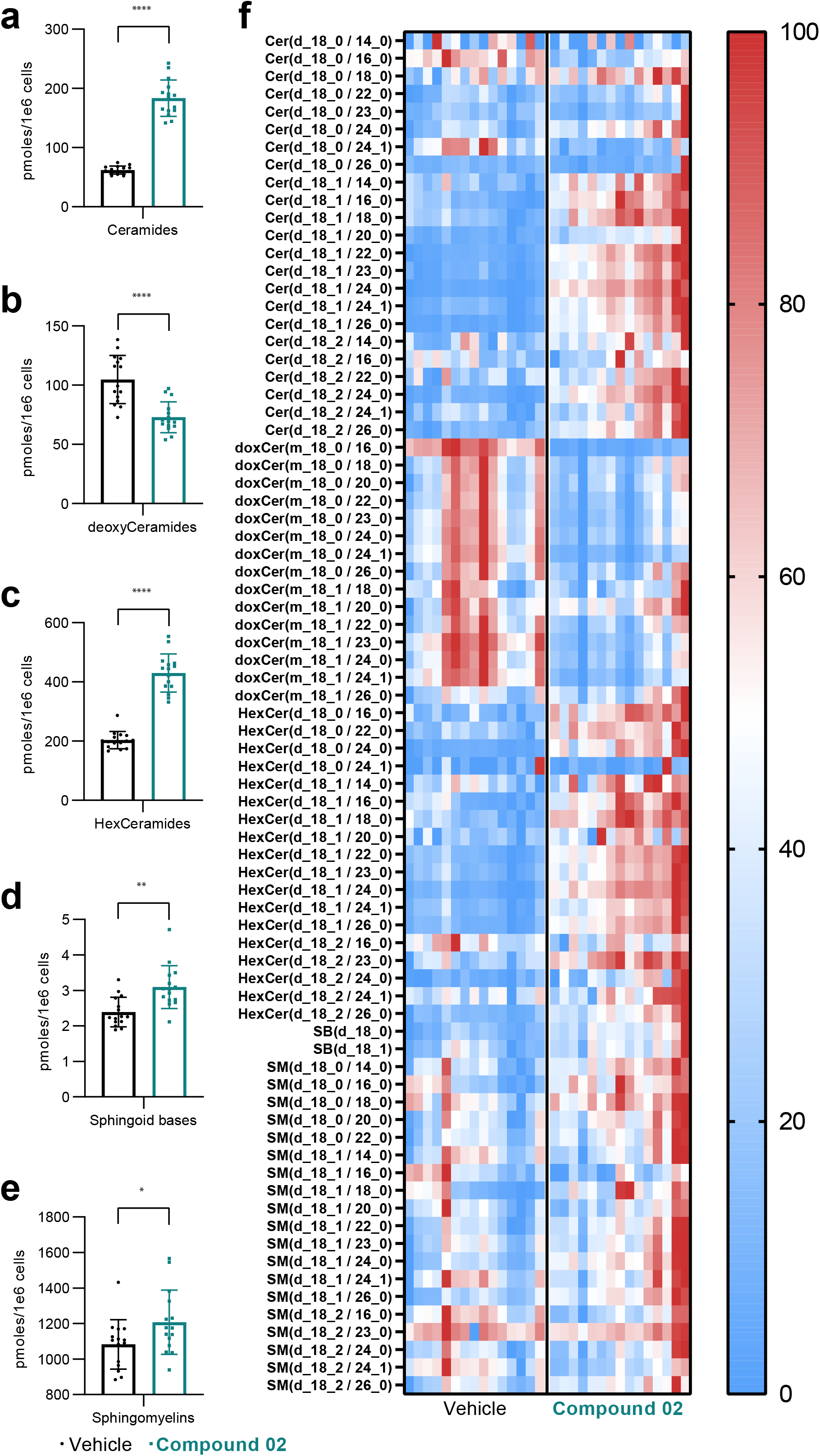
Sphingolipidomics performed in U937 cells. Total detected sphingolipids in cells after treatment with vehicle (0.025% DMSO) or **Compound 02** (25 μM) for 48 hours. **a**, Total ceramides, **b**, total deoxy-Ceramides, **c**, total hexosyl-ceramides, **d**, total sphingoid bases, and **e**, total sphingomyelins. **f**, Heat map of all detected species normalised by maximum and minimum per species. Composition of ceramides, deoxy-ceramides, hexosyl-ceramides and sphingomyelins. Data is from three independent experiments and plotted as mean ± SD. Asterisks indicate statistical significance over vehicle, * < 0.05, ** p < 0.005, *** p < 0.0005, **** p < 0.0001.

**Supplementary Table 1.**
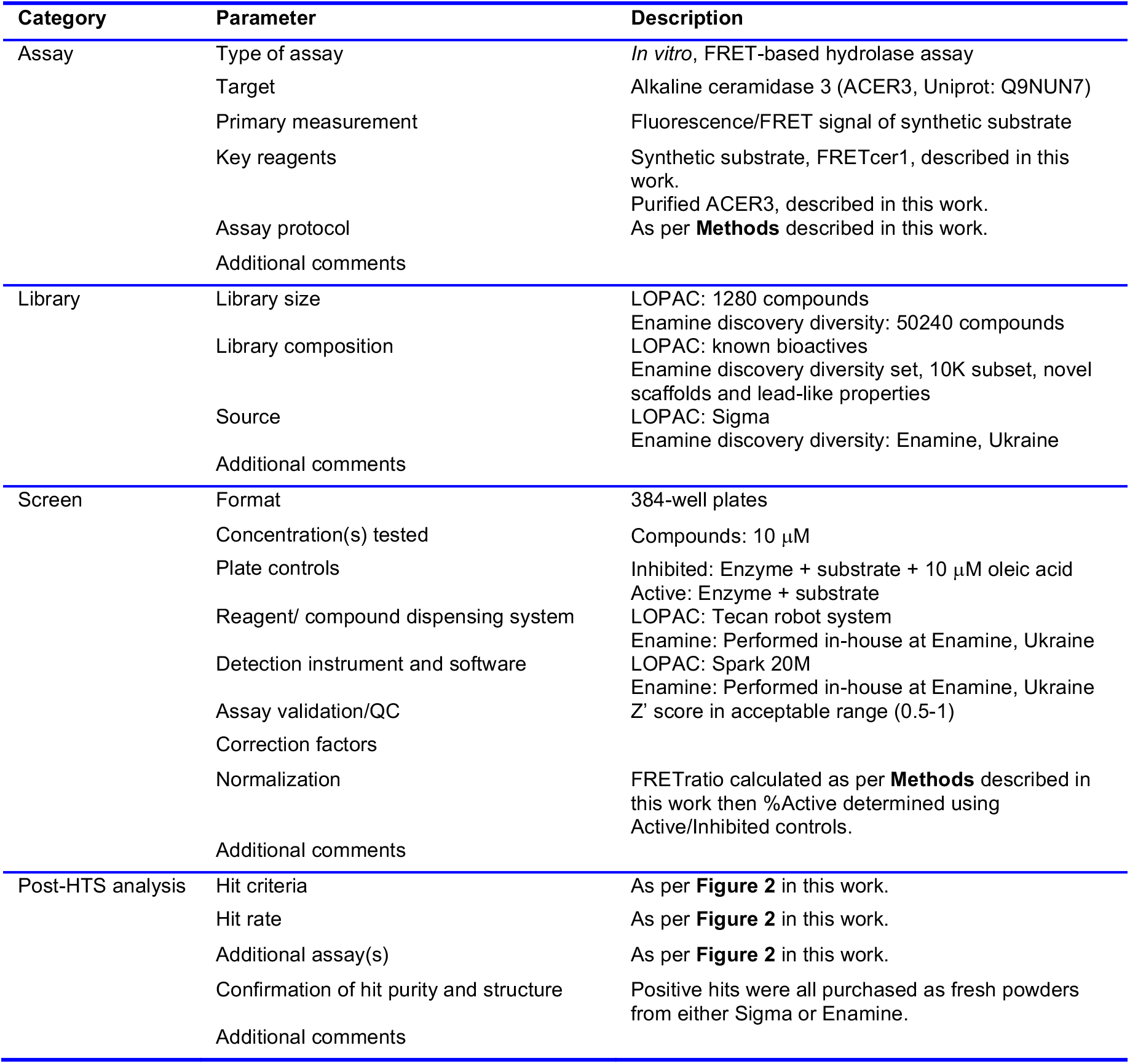
Small molecule screening data

## Supplementary Data

### I. General description of methods

Unless otherwise specified, all reactions were carried out in oven-dried (>120°C) glassware equipped with a magnetic stir bar and a rubber septum under a positive pressure of argon. Air- or moisture-sensitive reagents were transferred to the reaction vessel under positive pressure of argon via syringe. Air and/or moisture sensitive reactions were carried out in well dried glassware under an argon atmosphere with dry, freshly distilled solvents using standard syringe-cannula/septa techniques. Reactions were run at room temperature (20-25°C) unless otherwise noted in the experimental procedure and reported reaction temperatures refer to the external temperatures measured for the bath in which the reaction vessel was immersed. Heating was obtained using a silicone oil bath. For reactions run below room temperature, the term “-78°C” refers to a bath of acetone and dry ice, “-20°C” refers to a slurry of sodium chloride and ice-water bath, and “0°C” refers to an ice-water bath. Removal of residual solvents was accomplished by evacuation of the container for a period of 12-20 hours using a high vacuum line.

#### Chromatography

The thin layer chromatography studies were performed on pre-coated silica gel 60-F_254_ on aluminum sheets (Merck KGaA) and spots were detected by UV illumination (254 nm), and/or spraying with 1.3% ninhydrin solution, ceric ammonium molybdate (Seebach reagent) solution (25 g MoO_3_·H_3_PO_4_·H_2_O, 10 g Ce(SO_4_)_2_·4H_2_O, 60 ml H_2_SO_4_ and 905 ml of H_2_O) or KMnO_4_ solution (1.5 g KMnO_4_, 10 g K_2_CO_3_ and 1.25 ml 10% NaOH in 200 ml water) followed by heating. Preparative flash column chromatography was performed manually using glass columns of different size packed with Silica Gel 60M (0.04-0.063 mm) as stationary phase with indicated eluent systems in parenthesis following the description of purification. Solvent ratios for chromatography and R_f_ values are reported in v/v% ratios.

#### Spectroscopic Data

The structure of all synthesized compounds was confirmed with ^1^H NMR, ^13^C NMR, DEPT, ^31^P NMR and MS analysis. ^1^H, ^13^C and ^31^P NMR spectra were recorded on Bruker AVANCE II 300, AVANCED PX 300, AVANCE 400 and Bruker ADVANCE III 500 spectrometers (^1^H at 300, 400 or 500 MHz, ^13^C at 75.4, 101.2 or 125.7 MHz and ^31^P at 161.9 or 202.4 Hz) as solutions in CDCl_3_, CD_3_OD or mixtures of those at 25 °C. Chemical shifts (δ) are reported in parts per million (ppm): multiplicities are indicated as s (singlet), d (doublet), t (triplet), q (quartet), m (multiplet) and br (broad). Chemical shifts are given in ppm with respect to TMS as an external standard, (^1^H, APT, ^13^C, δ = 0.00) with calibration against the residual solvent signal or 85% H_3_PO_4_ (^31^P, δ= 0.00) as external standard. The coupling constants *J* are given in Hz.

#### Mass Spectroscopy

Mass spectroscopy (MS) experiments were recorded on an AGILENT 6120 UPLC–MS system consisting of an SQD (single quadrupole detector) mass spectrometer equipped with an electrospray ionization interface (ESI) in the positive and negative ion detection modes. The samples were separated on a Zorbax Eclipse Plus C18 column (particle size 1.8 µ m, 2.1 × 50 mm) using a UPLC pump at a flow rate of 0.8 ml per min with a ternary solvent system of MeOH-H_2_O-HCOOH, methanol (99.9% MeOH: 0.1% HCOOH, v/v). The column was first equilibrated using a mixture of 95% mobile phase A and 5% mobile phase B, and then 10 µ l of the sample was injected. This was followed by a ramp gradient over 2 min to 95% phase B and 5% phase A, which remained until 7 min, followed by a ramp gradient back down to 95% solvent A and 5% solvent B for 1 min, and column equilibration with the same mixture for 1 min. The detection was performed in full scan mode and the major observable molecular ion and selected fragments and clusters have been reported.

### II. Synthesis procedures and analytical data of compounds

#### II. 1 Synthesis of FRET-ceramide probes

**Scheme 1:**
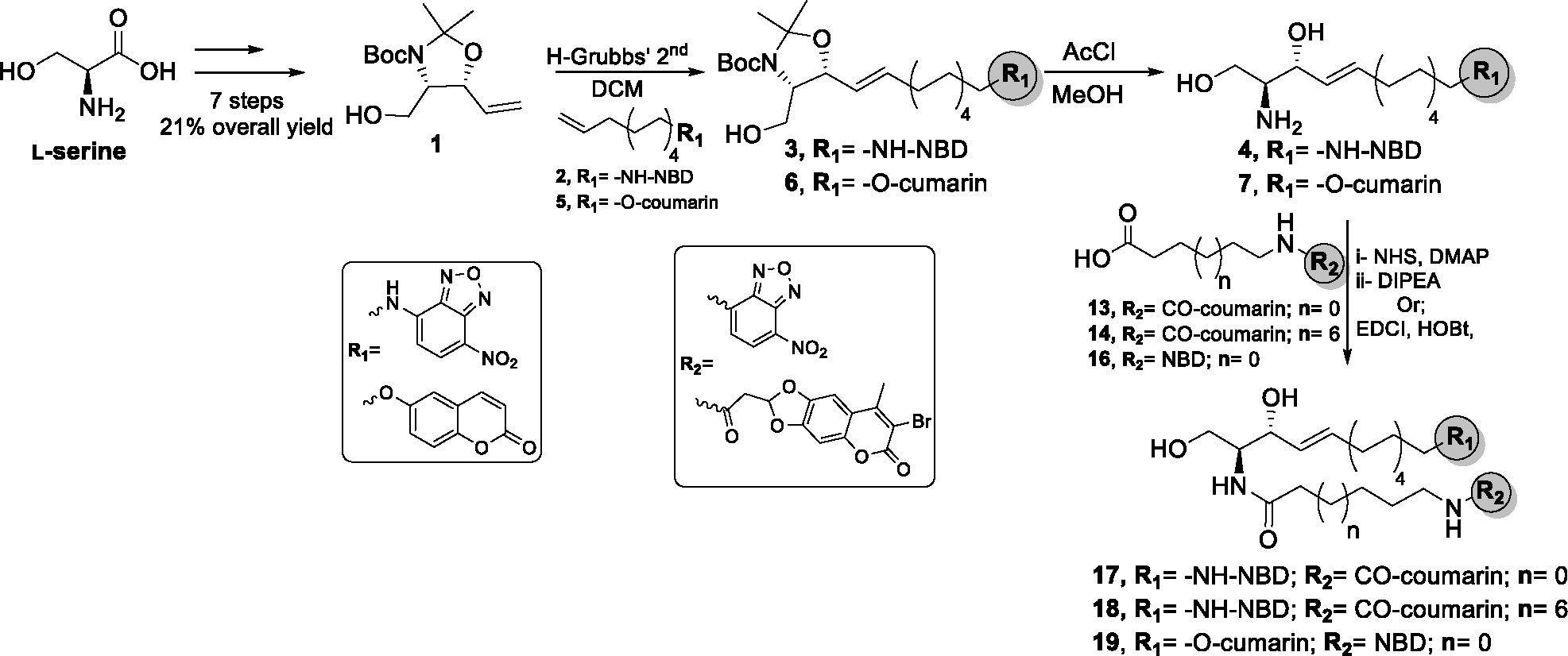
Synthesis of novel FRET-ceramide probes.

##### a) Synthesis of fluorescently labelled sphingosine analogues

**(4*S*,5*R*)-*tert*-Butyl-4-(hydroxymethyl)-2,2-dimethyl-5-vinyloxazolidine-3-carboxylate** (**1**).

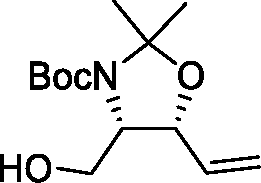

Compound **1** (2.1 g, 8.2 mmol) was synthesized starting from L-serine (4g, 38 mmol) in 7 steps with 21% overall yield. ^1^

R*_f_*: 0.48 (PE/EA 3:2, visualized with 1.3% ninhydrine). ^1^H NMR (CDCl_3,_ 500 MHz, ppm) δ 5.74-5.96 (m, 1H), 5.29-5.38 (m, 1H), 5.17-5.21 (m, 1H), 4.45-4.51 (m, 1H), 3.77-3.95 (m, 1H), 3.59 (dd, *J=* 5.8, 11.1 Hz, 1H), 3.34 (br.m, 1H), 1.41-1.46 (m, 6H), 1.25 (br.s, 9H). ^13^C NMR (CDCl_3,_ 126 MHz, ppm): δ 154.37, 131.98, 119.35, 93.30, 81.38, 76.52, 63.54, 61.89, 28.54, 24.86. ESI-MS *m/z* calcd for C_13_H_23_NO_4_Na [M+Na]^+^ 280.15; observed 280.2.

***N-(undecenyl)-7-nitrobenzo[c][1,2,5]oxadiazol-4-amine (2)***

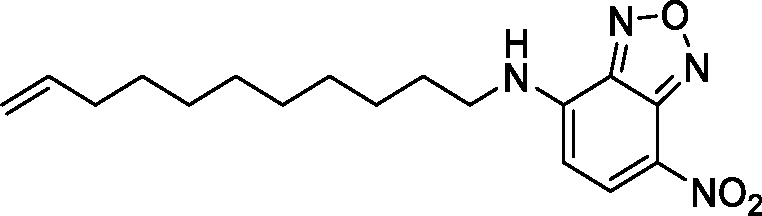

A stirred solution of 4-chloro-7-nitrobenzo[*c*][1,2,5]oxadiazole (NBD-Cl) (0.32 g, 1.6 mmol) in methanol (8 mL) at 0°C was treated with DIPEA (1.1 mL, 6.44 mmol), followed by 11-amino-1-undecene (0.3 g, 1.77 mmol). After the resulting reaction mixture stirred for 16h at ambient temperature (as judged by TLC analysis), the solvent was removed under reduced pressure. The resultant crude product was purified by flash column chromatography using cyclohexane and ethylacetate as eluents (20-25% ethylacetate in cyclohexane) to provide compound **2** as a red solid.^1–3^

Yield: 440 mg (83 %). R*_f_*: 0.48 (cyclohexane/EtOAc 4:1, visualized with UV illumination at 366 nm).^1^H NMR (CDCl_3,_ 500 MHz, ppm): δ 8.50 (d, *J*= 8.6 Hz, 1H), 6.30 (br. s, 1H), 6.18 (d, *J*= 8.7 Hz, 1H), 5.81 (ddt, *J*= 6.7, 10.2, 16.9 Hz, 1H), 4.91-5.03 (m, 2H), 3.50 (q, *J*= 7.0, 13.1 Hz, 2H), 2.04 (app. q, *J*= 6.8 Hz, 2H), 1.78-1.84 (m, 2H), 1.30-1.51 (m, 12H). ^13^C NMR (CDCl_3,_ 125 MHz, ppm): δ 144.38, 144.01, 139.24, 136.67, 114.3, 98.64, 44.14, 33.88, 29.51, 29.47, 29.31, 29.17, 28.99, 28.66, 27.06. ESI-MS: *m/z* calcd for C_17_H_25_N_4_O_3_ [M+H]^+^ 333.19; observed 333.2.

**(4*S*,5*R*)-*tert*-butyl-4-(hydroxymethyl)-2,2-dimethyl-5-((*E*)-11-((7-nitrobenzo[*c*][1,2,5]oxadi-azol-4-yl)amino)undec-1-en-1-yl)oxazolidine-3-carboxylate (3)**

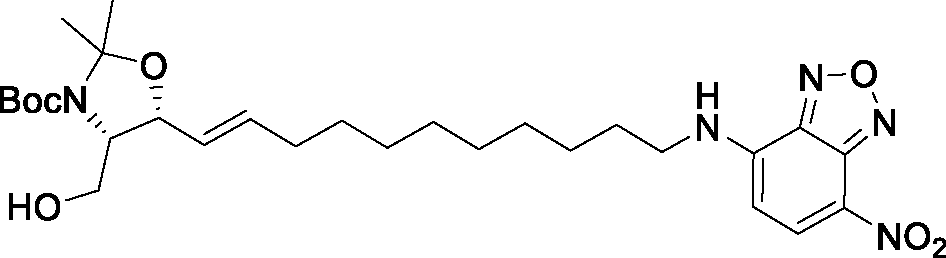

To a stirred solution of compound **1** (180 mg, 0.7 mmol.) in dry dichloromethane (10 mL) under argon atmosphere was added NBD-alkene cross partner **2** (920 mg, 2.8 mmol), followed by a catalytic amount of Hoveyda-Grubbs Catalyst 2^nd^ Generation (7 mol%). After the resulting reaction mixture was stirred for 16h at ambient temperature (until the TLC analysis showed no further change in the composition of the reaction mixture), the solvent was removed *in vacuo*. Purification of the resultant crude product by flash column chromatography using pet ether and ethylacetate as eluents (from 20-30% ethylacetate in pet ether) afforded protected NBD-labeled sphingosine **3** as a red waxy oil. ^1–3^

Yield: 230 mg (59%). R*_f_*: 0.4 (PE/EA 3:2, visualized with UV illumination at 366 nm). ^1^H NMR (CDCl_3,_ 500 MHz, ppm): δ 8.49 (d, *J*= 8.6 Hz, 1H), 6.29 (br. s, 1H), 6.17 (d, *J*= 8.6 Hz, 1H), 5.74-5.83 (m, 1H), 5.45-5.59 (m, 1H), 4.50-4.55 (m, 1H), 3.89-4.05 (m, 1H), 3.49-3.58 (m, 3H), 2.05-2.09 (m, 2H), 1.76-1.86 (m, 2H), 1.29-1.56 (m, 28H).^13^C NMR (CDCl_3,_ 125 MHz, ppm): δ 151.71, 144.24, 143.96, 136.52, 133.36, 129.32, 124.08, 98.35, 92.36, 69.81, 44.18, 32.17, 29.68, 29.20, 28.52, 28.35, 26.96. ESI-MS *m/z* calcd for C_28_H_44_N_5_O_7_ [M+H]^+^ 562.32; observed 562.3.

**(2*S*,3*R*,*E*)-2-amino-14-((7-nitrobenzo[*c*][1,2,5]oxadiazol-4-yl)amino)tetradec-4-ene-1,3-diol (4)**

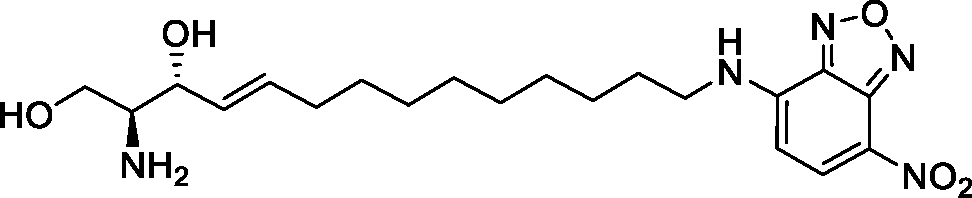

An ice-cold solution of triflouroacetic acid in dichloromethane (TFA-DCM) (2:1; 2 mL) was added dropwise over 10 min to a stirred solution of protected NBD-sphingosine **3** (0.1 g, 0.18 mmol) in dichloromethane (2 mL) at 0°C. The resulting reaction mixture was allowed to stir for 30 min at 0°C, gradually warmed to ambient temperature and stirred for additional 1h (as judged by TLC analysis for almost a complete deprotection; visualized with UV illumination at 366 nm). The reaction mixture was diluted with CH_2_Cl_2_ (20 mL) and subsequently quenched with a saturated NaHCO_3_ solution (15 mL). The layers were separated, and the aqueous layer was extracted with CH_2_Cl_2_ (4x 10 mL). The combined organic layer was sequentially washed with water (40 mL), brine (40 mL), dried over anhydrous Na_2_SO_4_, filtered and concentrated under reduced pressure. Purification of the resultant residue by flash column chromatography on silica gel using ethyacetate and isopropanol as eluents (3% isopropanol in ethylacetate) afforded the desired NBD-labeled sphingosine **4** as red-waxy solid. ^2, 3^

Yield: 53 mg (72%). R*_f_*: 0.56 (ethylacetate/isopropanol 4:1, visualized with UV illumination at 366 nm). ^1^H NMR (CD_3_OD, 500 MHz, ppm): δ 8.52 (d, *J* = 8.9 Hz, 1H). 6.33 (d, *J* = 8.9 Hz, 1H), 5.84 (m, 1H), 5.46 (tdd, *J =* 6.7 Hz, 15.2 Hz, 1H), 4.26-4.31 (m, 1H), 3.79 (dd, *J* = 4.1 Hz, 11.6 Hz, 1H), 3.57 (dd, *J* = 4.7 Hz, 11.6 Hz, 1H), 3.48-3.53 (br.s, 1H), 3.20-3.23 (m, 2H), 2.10 (m, 2H), 1,76-1,82 (m, 2H), 1.28-1.33 (m, 14H). ^13^C NMR (CD_3_OD, 125 MHz, ppm): δ 137.25, 135.14, 127.16, 97.93,. 69.6, 58.02, 57.15, 52.64, 31.91, 29.20, 29.14, 28.93, 28.82, 28.71, 26.63. ESI-MS *m/z* calcd for C_20_H_31_N_5_O_5_Na [M+Na]^+^ 444.22; observed 444.2.

**6-(undec-10-en-1-yloxy)-2H-chromen-2-one (5)**

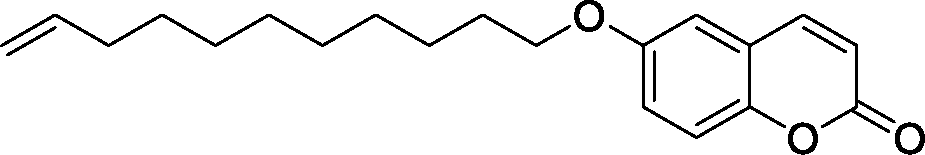

To a stirred solution of 11-bromoundecene (1g, 4.2 mmol) in dry DMF (10 mL) under argon atmosphere was added 6-hydroxycoumarin (740 mg, 4.6 mmol), followed by potassium carbonate (1.6 g, 12.6 mmol). After the resulting reaction mixture was allowed to stir at 70°C for 12h (as indicated by TLC analysis for complete substitution reaction), the reaction was quenched with water (100 mL). The resulting mixture was extracted with ethylacetate (3×100 mL) and the combined organic layers was washed with brine, dried over anhydrous Na_2_SO_4_, filtered and concentrated under reduced pressure. The obtained crude mixture was purified by flash column chromatography over silica gel using pet. ether and ethylacetate as eluents (0-10% ethylacetate in pet. ether) to afford the desired coumarin labelled alkene **6** as white solid.

Yield: 1.39 g (90 %). R*_f_*: 0.52 (PE/EA 8:2, visualized with UV illumination at 366 nm). ^1^H NMR (500 MHz, CDCl_3_, ppm) δ 7.64 (d, *J* = 9.5 Hz, 1H), 7.37 (d, *J* = 8.5 Hz, 1H), 6.84 (dt, *J* = 5.3, 2.4 Hz, 2H), 6.25 (d, *J* = 9.5 Hz, 1H), 5.83 (ddt, *J* = 16.9, 10.2, 6.7 Hz, 1H), 5.06 – 4.90 (m, 2H), 4.02 (t, *J* = 6.5 Hz, 2H), 2.06 (dd, *J* = 14.1, 6.8 Hz, 2H), 1.89 – 1.77 (m, 2H), 1.55 – 1.27 (m, 12H). ^13^C NMR (125 MHz, CDCl_3_, ppm) δ 162.58, 161.43, 156.07, 143.58, 139.33, 128.82, 114.28, 113.14, 113.05, 112.49, 101.46, 68.80, 33.92, 29.60, 29.53, 29.44, 29.23, 29.10, 29.04, 26.08. ESI-MS: *m/z* calcd for C_20_H_27_O_3_ [M+H]^+^ 315.19; observed 315.2.

**(4*S*,5*R*)-*tert*-butyl-4-(hydroxymethyl)-2,2-dimethyl-5-((*E*)-11-((2-oxo-2H-chromen-7-yl)-oxy)undec-1-en-1-yl)oxazolidine-3-carboxylate (6)**

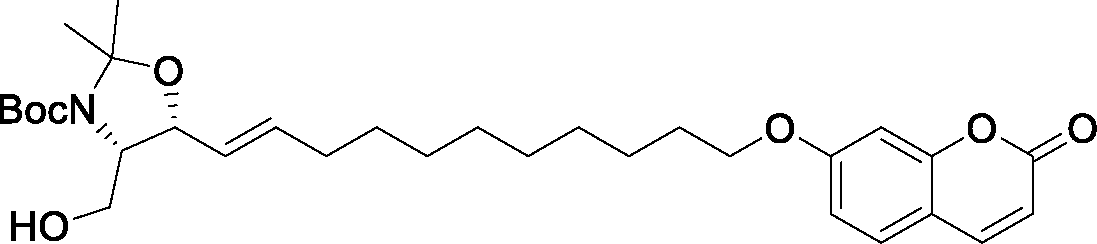

Under exclusion of light, to a stirred solution of compound **1** (100 mg, 0.39 mmol.) in dry dichloromethane (5 mL), under argon atmosphere, was added the coumarin alkene partner **5** (0.37 g, 1.2 mmol), followed by a catalytic amount of Hoveyda-Grubbs Catalyst 2^nd^ Generation (7 mol%). After the resulting reaction mixture was stirred at ambient temperature for 12h (until the TLC analysis showed no further change in the composition of the reaction mixture; visualized with UV illumination at 366 nm), the solvent was removed under reduced pressure. Purification of the resultant crude mixture by flash column chromatography using pet. ether and ethylacetate as eluents (from 25-30% ethylacetate in pet. ether) furnished the desired protected labelled-sphingosine **6** as white waxy-solid.

Yield: 147 mg (69%). R*_f_*: 0.47 (PE/EA 3:2, visualized with UV illumination at 366 nm). ^1^H NMR (500 MHz, CDCl_3_, ppm)) δ 7.56 (d, *J* = 9.4 Hz, 1H), 7.29 (d, *J* = 8.6 Hz, 1H), 6.76 (d, *J* = 8.5 Hz, 1H), 6.73 (d, *J* = 2.0 Hz, 1H), 6.17 (d, *J* = 9.4 Hz, 1H), 5.75 – 5.68 (m, 1H), 5.47 (dd, *J* = 15.4, 6.4 Hz, 1H), 4.26 (m, 1H), 3.94 (t, *J* = 5.6 Hz, 2H), 3.87 (dd, *J* = 11.4, 3.5 Hz, 1H), 3.64 (dd, *J* = 11.0, 3.2 Hz, 1H), 3.54 (m, 1H), 1.99 (dd, *J* = 14.0, 6.9 Hz, 2H), 1.77 – 1.71 (m, 2H), 1.44 – 1.35 (m, 15H), 1.31 – 1.19 (m, 12H).^13^C NMR (126 MHz, CDCl_3_, ppm) δ 162.59, 161.47, 156.08, 143.61, 143.59, 134.13, 130.49, 129.20, 128.83, 113.15, 113.05, 112.51, 101.50, 75.03, 68.80, 65.93, 62.81, 49.79, 32.39, 31.05, 29.55, 29.47, 29.39, 29.24, 29.19, 29.10, 29.06, 28.52, 26.04. ESI-MS *m/z* calcd for C_31_H_46_NO_7_ [M+H]^+^ 544.33; observed 544.3.

**7-(((12*R*,13*S*,*E*)-13-amino-12,14-dihydroxytetradec-10-en-1-yl)oxy)-2*H*-chromen-2-one (7)**

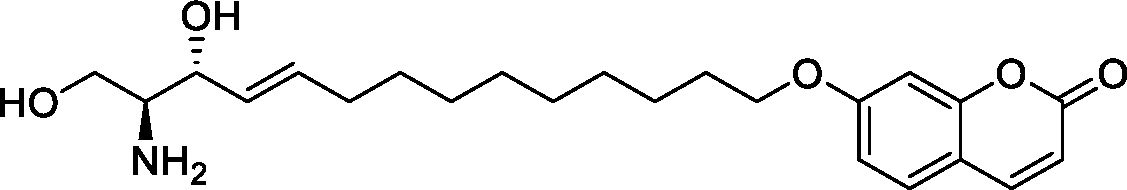

A solution of compound **6** (120 mg, 0.22 mmol) in methanol (2 mL) at 0°C was treated with acetyl chloride (150 µL, 2.2 mmol). After the resulting reaction mixture was stirred at 0°C for 1h and for additional 2h at ambient, the solvent was removed under reduced pressure. The obtained residue was treated under stirring with petroleum ether (5mL) to provide CUM-labelled sphingosine **7** as a white waxy-solid. The resultant crude product was obtained in pure form (as indicated by TLC; NMR and LC-MS analysis) and was used directly in the next step.

Yield: 75 mg (84%). R*_f_*: 0.52 (ethylacetate/isopropanol 4:1, visualized with UV illumination at 366 nm). ^1^H NMR (500 MHz, MeOD, ppm) δ 7.85 (d, *J* = 9.6 Hz, 1H), 7.50 (d, *J* = 8.6 Hz, 1H), 6.90 (dd, *J* = 8.6, 2.4 Hz, 1H), 6.85 (d, *J* = 2.3 Hz, 1H), 6.23 (d, *J* = 9.4 Hz, 1H), 5.85 (dt, *J* = 14.0, 6.4 Hz, 1H), 5.46 (dd, *J* = 15.4, 6.8 Hz, 1H), 4.32 – 4.27 (m, 1H), 4.05 (t, *J* = 6.4 Hz, 3H), 3.79 (dd, *J* = 11.6, 4.0 Hz, 1H), 3.67 (dd, *J* = 11.6, 8.3 Hz, 1H), 3.20 (dt, *J* = 8.4, 4.3 Hz, 1H), 2.09 (dd, *J* = 14.0, 7.0 Hz, 2H), 1.84 – 1.77 (m, 1H), 1.51 – 1.31 (m, 12H). ^13^C NMR (126 MHz, MeOD, ppm) δ 163.85, 163.37, 156.87, 145.66, 136.41, 130.19, 128.05, 114.06, 113.68, 113.02, 102.08, 70.75, 69.59, 59.20, 58.25, 33.35, 33.15, 30.38, 30.30, 30.19, 30.11, 29.93, 29.89, 26.84. ESI-MS *m/z* calcd for C_23_H_34_NO_5_ [M+H]^+^ 404.24; observed 404.2.

##### b) Synthesis of fluorescently labeled fatty acids

***(i)- Synthesis of coumarin-labelled fatty acids:***

**Scheme 2:**
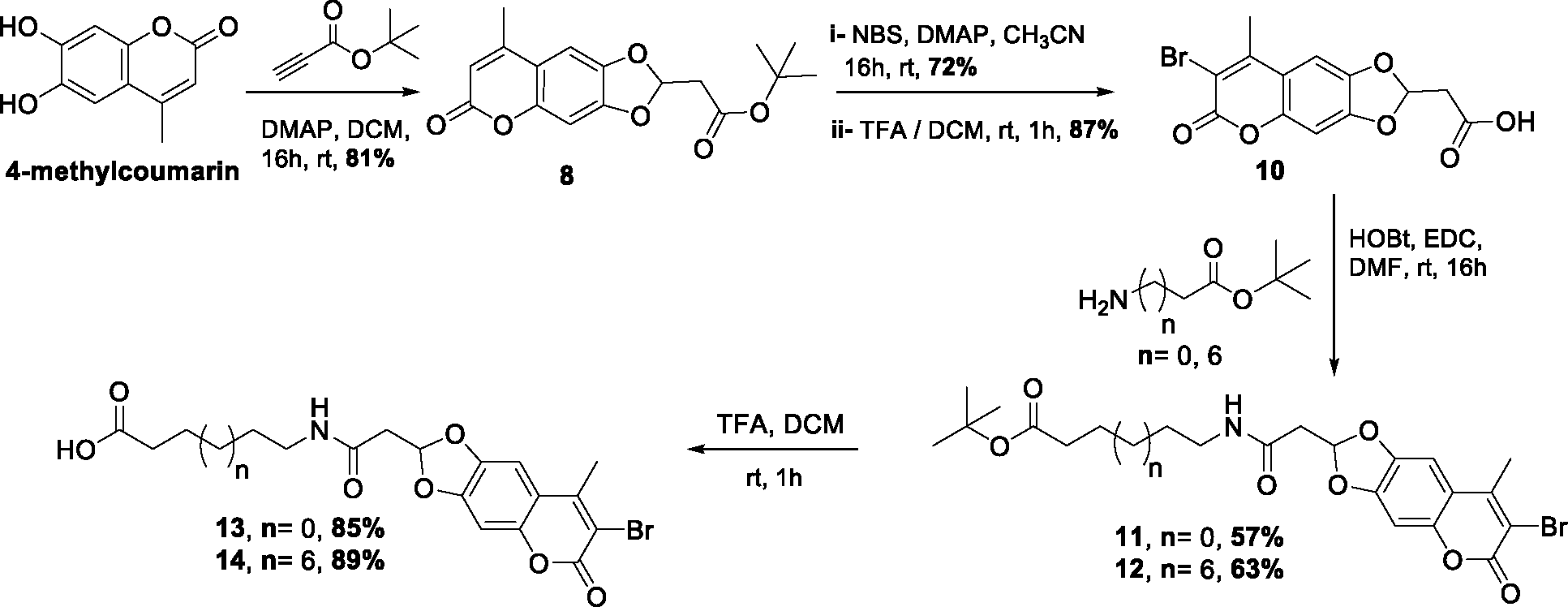
Synthesis of coumarin-labelled fatty acids.

***tert*-butyl 2-(8-methyl-6-oxo-6*H*-[1,3]dioxolo[4,5-*g*]chromen-2-yl)acetate (8)**

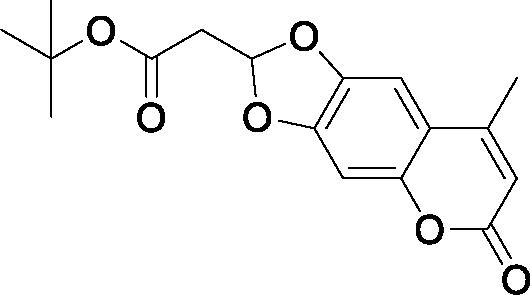

A stirred solution of *tert*-butyl propiolate (1.1 g, 8.7 mmol) in dry CH_2_Cl_2_ (50 mL) at ambient temperature under argon atmosphere was treated with 6,7-dihydroxy-4-methylcoumarin (1.2 g, 6.2 mmol), followed by 4-dimethylaminopyridine (1.3 g, 10.5 mmol, DMAP). After being stirred at the same conditions for 16h, the reaction mixture was diluted with CH_2_Cl_2_ (100 mL), and subsequently washed with saturated NaHCO_3_ solution (90 mL). The aqueous layer was extracted with CH_2_Cl_2_ (3x 80 mL), and the combined organic layers were washed with brine solution (70 mL), dried over anhydrous Na_2_SO_4_, filtered and concentrated *in vacuo*. Purification of the resultant crude mixture by flash column chromatography using cyclohexane and ethylacetate as eluents (from 20-25% ethylacetate in cyclohexane) afforded compound **8** as white solid. ^4^

Yield: 1.6 g (81 %). R*_f_*: 0.4 (cyclohexane/ethylacetate 4:1, visualized with UV illumination at 366 nm). ^1^H NMR (CDCl_3,_ 500 MHz, ppm): δ 6.97 (s, 1H). 6.79 (s, 1H), 6.61 (t, *J* = 5.2 Hz, 1H), 6.19 (s 1H), 2.98 (d, *J* = 5.1 Hz, 2H), 2.45 (d, *J* = 1.3 Hz, 3H), 1.46 (s, 9H). ESI-MS *m/z* calcd for C_17_H_19_O_6_ [M+H]^+^ 319.12; observed 319.1.

***tert*-butyl 2-(7-bromo-8-methyl-6-oxo-6*H*-[1,3]dioxolo[4,5-*g*]chromen-2-yl)acetate (9)**

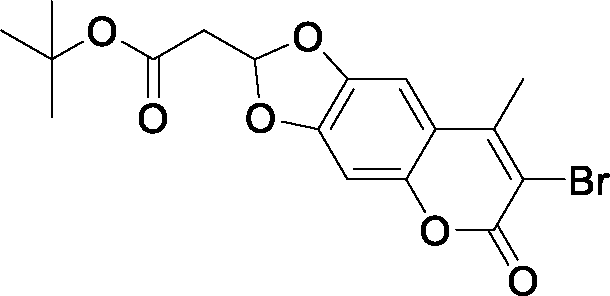

To a stirred solution of compound **8** (1.4 g, 4.4 mmol) in dry acetonitrile (28 mL) under argon atmosphere at ambient temperature was added *N*-bromosuccinimide (1.6 g, 9.2 mmol, NBS), followed by a catalytic amound of sodium acetate (54 mg, 0.44 mmol, 10 mol%). Afer reaction mixture was stirred at the same conditions for 16h, the solvent was removed under reduced pressure. The resultant residue was partioned between CH_2_Cl_2_ and brine solution, and the layers were separated. The aqueous layer was then extracted several time with CH_2_Cl_2_ (3x 70 mL), and the combined organic layers were dried over anhydrous Na_2_SO_4_, filtered and concentrated. The obtained crude residue was purified by flash column chromatography on silica gel using cyclohexane and ethylacetate as eluents (from 15-20% ethylacetate in cyclohexane) to provide compound **9** as yellowish white solid.^4^

Yield: 1.25 g (72 %). R*_f_*: 0.5 (cyclohexane/ethylacetate 4:1, visualized with UV illumination at 366 nm). ^1^H NMR (CDCl_3,_ 500 MHz, ppm): δ 6.99 (s, 1H). 6.81 (s, 1H), 6.59 (t, *J* = 5.2 Hz, 1H), 2.94 (d, *J* = 5.2 Hz, 2H), 2.55 (s, 3H), 1.47 (s, 9H). ^13^C NMR (CDCl_3,_ 125 MHz, ppm): δ 166.95, 157.51, 151.08, 150.97, 149.04, 145.38, 113.84, 110.45, 102.69, 98.35, 82.47, 41.36, 28.17, 20.17. ESI-MS *m/z* calcd for C_17_H_18_O_6_Br [M+H]^+^ 397.03; observed 397.0.

**2-(7-bromo-8-methyl-6-oxo-6*H*-[1,3]dioxolo[4,5-*g*]chromen-2-yl)acetic acid (10)**

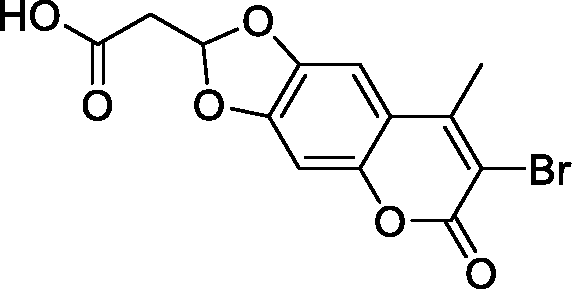

A solution of compound **9** (1g, 2.5 mmol) in CH_2_Cl_2_ (25 mL) at 0°C was treated dropwise with a solution of triflouroacetic acid in dichloromethane (TFA-DCM) (3:1; 8 mL). After being stirred at the same conditions for 30min, and for additional 1h at ambient temperature (as monitored by TLC analysis for complete deprotection of *tert*-butyl ester), the reaction mixture was diluted with CH_2_Cl_2_ (80 mL) and subsequently quenched with a saturated NaHCO_3_ solution (70 mL). The layers were separated, and the aqueous layer was extracted with CH_2_Cl_2_ (3x 50 mL). The combined organic layer was sequentially washed with water (100 mL), brine (100 mL), dried over anhydrous Na_2_SO_4_, filtered and concentrated under reduced pressure using a high vacuum line to afford compound **10** as yellow solid. The compound was obtained in a pure form (as determined by NMR, ESI-MS analysis), and was used directly in the next step without any further purification. ^4^

Yield: 740 mg (87 %). R*_f_*: 0.3 (DCM/MeOH 9:1, visualized with UV illumination at 366 nm). ^1^H NMR (DMSO, 500 MHz, ppm): δ 7.21 (s, 1H), 6.86 (s, 1H), 6.69 (t, *J* = 5.1 Hz, 1H), 3.14 (d, *J* = 5.1 Hz, 2H), 2.57 (s, 3H). ^13^C NMR (DMSO, 125 MHz, ppm): δ 169.12, 157.31, 151.89, 151.10, 149.42, 145.38, 114.23, 110.47, 103.12, 98.42, 40.49, 20.26. ESI-MS *m/z* calcd for C_13_H_10_O_6_Br [M+H]^+^ 340.97; observed 341.0.

***tert*-Butyl-6-(2-(7-bromo-8-methyl-6-oxo-6*H*-[1,3]dioxolo[4,5-*g*]chromen-2-yl)acetamido)-hexanoate (11)**

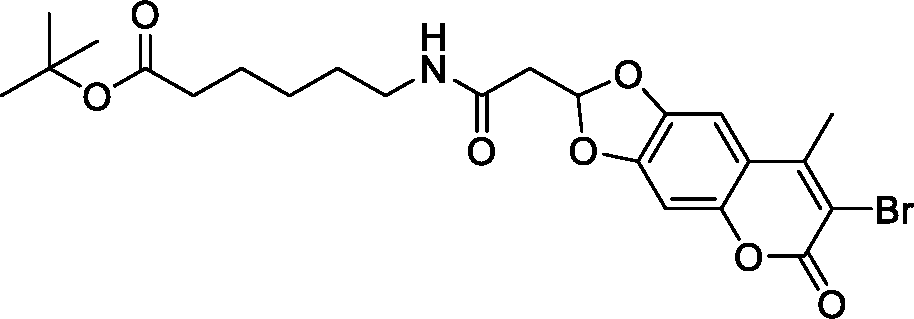

To a stirred solution of compound **10** (445 mg, 1.3 mmol) in a mixture of DCM:DMF (3:1; 13 mL) at 0°C under argon atmosphere was sequentially added *N*,*N*-diisopropylethylamine (DIPEA) (0.68 mL, 3.9 mmol), 1-hydroxybenzotriazol hydrate (HOBt) (0.22 g, 1.6 mmol) and 1-ethyl-3-(3-dimethylaminopropyl)carbodiimide hydrochloride (EDCI.HCl) (0.31 g, 1.6 mmol). After the resulting mixture was stirred at the same conditions for 20 min (during while a clear solution was formed), a solution of *tert*-butyl 6-aminohexanoate (0.22 g, 1.2 mmol) in dry DMF (2.4 mL) was dropwisely added over a period of 10 min. The resulting reaction mixture was allowed to stir at 0°C for 30 min, and for additional 12h at ambient temperature (as judged by TLC analysis for a complete consumption of the amine). The solvent was subsequently removed under reduced pressure, and the obtained residue was portioned between ethylacetate (50 mL) and saturated NaHCO_3_ solution (50 mL). The layers were separated and the aqueous layer was extracted several times with ethylacetate (3×50 mL). The organic extracts were combined and successively washed with water (2×100 mL) and brine solution (100 mL), dried over anhydrous NaSO_4_, filtered and concentered in *vacuo*. The resultant residue was immediately purified by flash column chromatography on silica gel using cyclohexane and ethylacetate as eluents (from 30-45% ethylacetate in cyclohexane) to afford compound **11** as pale-yellow solid.

Yield: 380 mg (57 %). R*_f_*: 0.55 (DCM/MeOH 9:1, visualized with UV illumination at 366 nm). ^1^H NMR (CDCl_3,_ 500 MHz, ppm): δ 6.97 (s, 1H), 6.76 (s, 1H), 6.64 (t, *J* = 5.2 Hz, 1H), 6.05 (t, *J* = 5.4 Hz, 1H), 2.90 (d, *J* = 5.2 Hz, 2H), 2.53 (s, 3H), 2.22 (t, *J* = 7.3 Hz, 2H), 1.57 (ddt, *J* = 7.3, 14.8, 24.2, Hz, 2H), 1.43 (s, 9H), 1.32 – 1.39 (m, 2H). ^13^C NMR (CDCl_3,_ 125 MHz, ppm): δ 173.25, 166.44, 20.17, 157.45, 151.12, 150.89, 148.94, 145.32, 111.10, 110.40, 113.82, 102.75, 98.32, 80.39, 42.23, 39.56, 35.38, 29.07, 28.25, 26.30, 24.53. ESI-MS *m/z* calcd for C_22_H_29_NO_7_Br [M+H]^+^ 510.11; observed 510.1.

***tert*-butyl 11-(2-(7-bromo-8-methyl-6-oxo-6*H*-[1,3]dioxolo[4,5-g]chromen-2-yl)acetamido)-undecanoate (12)**

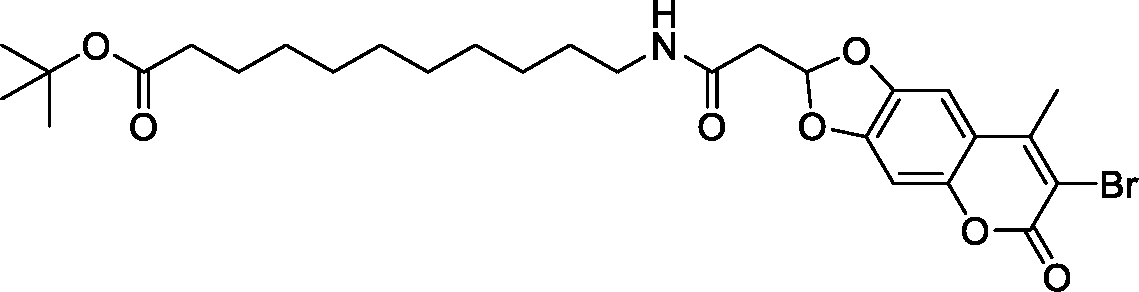

A stirred solution of coumarin derivative **10** (340 mg, 1 mmol) in dry DCM (10 mL) under argon atmosphere at 0°C was treated with DIPEA (550 µL, 3 mmol) followed by EDCI.HCl (250 mg, 1.3 mmol) and HOBt (175 mg, 1.3 mmol). After the resulting mixture was stirred at the same conditions for 15 min, *tert*-butyl 11-aminoundecanoate (285 mg, 1.1 mmol) was added. The resulting reaction mixture was allowed to stir at ambient temperature for 12h (as indicated by TLC analysis), and then the solvent was evaporated under vacuum. The obtained residue was then portioned between DCM (60 mL) and saturated NaHCO_3_ solution (60 mL), and the layers were separated. The aqueous layer was extracted with DCM, and the combined organic layers were washed with brine, dried over anhydrous NaSO_4_, filtered and concentered under reduce pressure. Flash column chromatography of the obtained crude mixture over silica gel using pet. ether and ethylacetate as eluents (25-30% ethylacetate in pet. ether) afforded the desired product **12** as pale-yellow solid.

Yield: 365 mg (63 %). R*_f_*: 0.55 (PE/EA 1:3, visualized with UV illumination at 366 nm). ^1^H NMR (500 MHz, CDCl_3_, ppm) δ 6.95 (s, 1H), 6.74 (s, 1H), 6.64 (t, *J* = 5.1 Hz, 1H), 5.98 (br.s, 1H), 3.28 (dd, *J* = 13.1, 6.9 Hz, 2H), 2.90 (d, *J* = 5.1 Hz, 2H), 2.52 (s, 3H), 2.18 (t, *J* = 7.5 Hz, 2H), 1.55 (t, *J* = 7.3 Hz, 2H), 1.50 (dd, *J* = 14.2, 7.1 Hz, 2H), 1.43 (s, 9H), 1.26 (m, 12H). ^13^C NMR (126 MHz, CDCl_3_, ppm) δ 173.48, 166.36, 157.40, 151.07, 150.91, 148.93, 145.35, 113.79, 111.16, 110.38, 102.71, 98.28, 80.06, 42.23, 39.93, 35.73, 29.57, 29.47, 29.37, 29.34, 29.18, 28.26, 26.96, 25.20, 20.14. ESI-MS *m/z* calcd for C_28_H_39_NO_7_Br [M+H]^+^ 580.19; observed 580.2.

**6-(2-(7-bromo-8-methyl-6-oxo-6*H*-[1,3]dioxolo[4,5-*g*]chromen-2-yl)acetamido)hexanoic acid (13)**

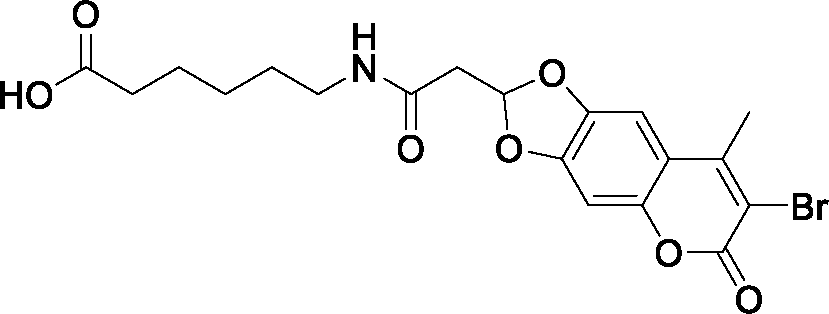

To a solution of compound **11** (0.3 g, 0.59 mmol) in CH_2_Cl_2_ (6 mL) at 0°C was added dropwise a solution of triflouroacetic acid in DCM (3:1; 4mL). The resulting reaction mixture was stirred at the same conditions for 30min and for additional 1h at ambient temperature (as judged by TLC analysis for complete deprotection), before it was carefully quenched with saturated NaHCO_3_ solution (50 mL). The layers were separated, and the aqueous layer was extracted again with DCM (3x 50 mL). The combined organic phases were washed with brine solution (100 ml), dried over anhydrous NaSO_4_, filtered and concentered using a high vacuum line to afford compound **13** as a pale-yellow solid. The compound was obtained in a pure form (as judged by NMR and ESI-MS analysis) and was used directly in the next step without any further purification.

Yield: 230 mg (85 %). R*_f_*: 0.35 (DCM/MeOH 4:1, visualized with UV illumination at 366 nm). ^1^H NMR (DMSO, 500 MHz, ppm): δ 8.08 (t, *J* = 5.5 Hz, 1H), 7.38 (s, 1H), 7.13 (s, 1H), 6.67 (t, *J* = 5.4 Hz, 1H), 3.06 (dd, *J* = 6.8, 12.7 Hz, 2H), 2.85 (d, *J* = 5.4 Hz, 2H), 2.54 (s, 3H), 2.19 (t, *J* = 7.4 Hz, 2H), 1. 46 – 1. 53 (m, 2H), 1. 36 – 1. 43 (m, 2H), 1.30 – 1.23 (m, 2H). ^13^C NMR (DMSO, 125 MHz, ppm): δ 174.44, 166.00, 156.49, 152.04, 150.67, 148.41, 144.85, 113.20, 111.47, 108.94, 103.45, 97.71, 40.77, 38.36, 33.61, 28.70, 25.92, 24.19, 19.99. ESI-MS *m/z* calcd for C_19_H_21_NO_7_Br [M+H]^+^ 454.05; observed 454.1.

**11-(2-(7-bromo-8-methyl-6-oxo-6H-[1,3]dioxolo[4,5-g]chromen-2-yl)acetamido)undecanoic acid (14)**

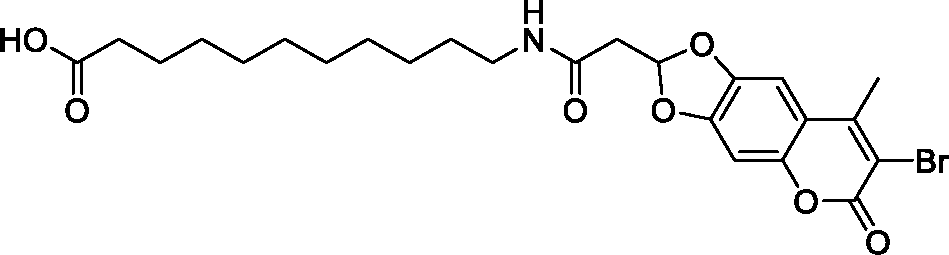

A stirred solution of compound **12** (300 mg, 0.52 mmol) in DCM (20 mL) at ambient temperature was dropwisely treated with trifluoroacetic acid (5mL). The resulting reaction mixture was stirred at the same conditions for 1h, and subsequently the solvent was removed under reduced pressure. The resultant residue was treated with pet. ether under stirring, during while a faint brown solid formed. The obtained solid was filterate and washed with DCM to afford the desired coumarine fatty acid **14** as a pale-yellow solid. The obtained product was pure (as indicated by UV-MS analysis and NMR analysis) and was used directly in the next step.

Yield: 240 mg (89 %). R*_f_*: 0.41 (DCM/MeOH 6:1, visualized with UV illumination at 366 nm). ^1^H NMR (500 MHz, DMSO, ppm) δ 8.07 (t, *J* = 5.4 Hz, 1H), 7.37 (s, 1H), 7.12 (s, 1H), 6.68 (t, *J* = 5.3 Hz, 1H), 3.05 (dd, *J* = 12.5, 6.6 Hz, 2H), 2.86 (d, *J* = 5.3 Hz, 2H), 2.54 (s, 3H), 2.18 (t, *J* = 7.3 Hz, 2H), 1.52 – 1.44 (m, 2H), 1.41 – 1.20 (m, 14H).. ^13^C NMR (126 MHz, DMSO, ppm) δ 174.49, 165.95, 156.47, 152.01, 150.69, 148.40, 144.87, 113.19, 111.46, 108.93, 103.41, 97.67, 40.77, 38.52, 33.65, 28.97, 28.95, 28.88, 28.74, 28.55, 26.36, 24.49, 19.97. ESI-MS *m/z* calcd for C_24_H_31_NO_7_Br [M+H]^+^ 524.13; observed 524.1.

***(ii) Synthesis of NBD-labelled fatty acid:***

***tert*-butyl 6-((7-nitrobenzo[c][1,2,5]oxadiazol-4-yl)amino)hexanoate (15)**

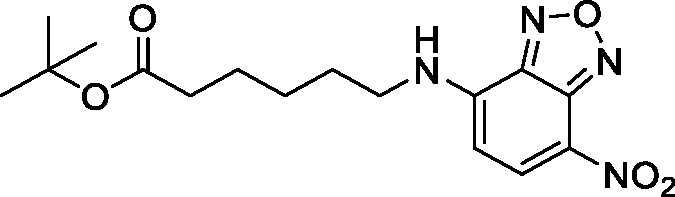

Under exclusion of light, a stirred solution of 4-chloro-7-nitrobenzo[*c*][1,2,5]oxadiazole (0.5g, 2.65 mmol) in methanol (8 mL) at 0°C was treated with *N,N*-diisopropylethylamine (2.5 mL, 15 mmol), followed by a dropwise addition of a solution of *tert*-butyl 6-aminohexanoate (0.5g, 2.9 mmol) in methanol (20 mL) over 20min. After the resulting reaction mixture was stirred for 12h at ambient temperature (as judged by TLC analysis for a complete reaction), the solvent was removed under reduced pressure. Purification of the obtained crude mixture by flash column chromatography on silica gel using pet. ether and ethylacetate as eluents (15-20% ethylacetate in pet. ether) furnished compound **15** as red solid.

Yield: 670 mg (73 %). R*_f_*: 0.45 (P.E/E.A 3:2, visualized with UV illumination at 254 nm). ^1^H NMR (400 MHz, CDCl_3_) δ 8.48 (d, *J* = 8.6 Hz, 1H), 6.39 (s, 1H), 6.17 (d, *J* = 8.7 Hz, 1H), 3.52 (dd, *J* = 13.1, 6.7 Hz, 2H), 2.27 (t, *J* = 7.2 Hz, 2H), 1.90 – 1.78 (m, 2H), 1.68 (dt, *J* = 14.8, 7.2 Hz, 2H), 1.55 – 1.46 (m, 2H), 1.44 (s, 9H). ^13^C NMR (101 MHz, CDCl_3_) δ 187.58, 172.95, 144.40, 144.03, 136.61, 124.11, 98.67, 80.57, 43.83, 35.21, 28.24, 26.39, 24.46. ESI-MS *m/z* calcd for C_16_H_23_N_4_O_5_ [M+H]^+^ 351.17; observed 351.2.

**Synthesis of 6-((7-nitrobenzo[c][1,2,5]oxadiazol-4-yl)amino)hexanoic acid (16)**

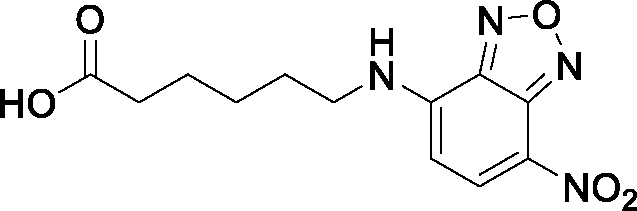

A stirred solution of compound **15** (0.4 g, 1.15 mmol) in CH_2_Cl_2_ (6 mL) at 0°C was dropwisely treated with triflouroacetic acid (3 mL). After the resulting reaction mixture was stirred at ambient temperature for 2h (as judged by TLC analysis for complete deprotection), the solvent was removed under reduced pressure. The residue was diluted with CH_2_Cl_2_ (50 mL) and washed with water (50 mL), dried over anhydrous NaSO_4_, filtered and concentered to provide compound **X** as a red solid. Compound **16** was obtained in a pure form (as judged by NMR and ESI-MS analysis) and was directly used in the next step without any further purifications. ^2, 5^

Yield: 300 mg (89 %). ^1^H NMR (400 MHz, DMSO, ppm) δ 9.52 (s, 1H), 8.49 (d, *J* = 8.8 Hz, 1H), 6.40 (d, *J* = 9.0 Hz, 1H), 2.21 (t, *J* = 7.3 Hz, 2H), 1.73 – 1.63 (m, 2H), 1.59 – 1.49 (m, 2H), 1.42 – 1.34 (m, 2H). ^13^C NMR (101 MHz, DMSO, ppm) δ 207.65, 174.38, 145.15, 144.41, 137.92, 99.08, 43.21, 33.54, 27.34, 25.93, 24.14. ESI-MS *m/z* calcd for C_12_H_15_N_4_O_5_ [M+H]^+^ 295.09; observed 295.1.

##### c) Synthesis of FRET-ceramide probes

**6-(2-(7-bromo-8-methyl-6-oxo-6*H*-[1,3]dioxolo[4,5-*g*]chromen-2-yl)acetamido)-*N*-((2*S*,3*R*, *E*)-1,3-dihydroxy-14-((7-nitrobenzo[*c*][1,2,5]oxadiazol-4-yl)amino)tetradec-4-en-2-yl)hexanamide (17, FRETcer1)**

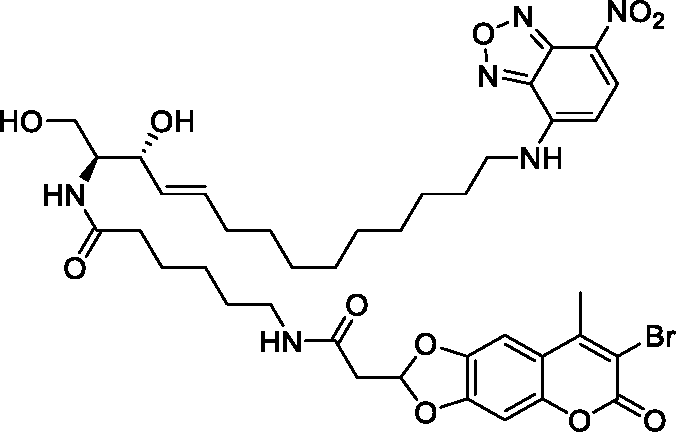

To a stirred solution of compound **13** (32 mg, 69 µmol) in a dry DMF (1 mL) at 0°C under argon atmosphere was successively added DIPEA (38 µL, 0.21 mmol), followed by HOBt (11.2 mg, 83 µmol) and EDCI (16 mg, 83 µmol). The resulting mixture was allowed to stir at the same conditions for 5 min and for additional 15 min at ambient temperature; during while a clear solution was formed. To this clear mixture, a solution of NBD-sphingosine **4** (27 mg, 62 µmol) in dry DMF (0.5 mL) under argon atmosphere was slowly added. After the resulting reaction mixture was stirred under exclusion of light at ambient temperature for 16h (as monitored by TLC analysis), the solvent was removed under reduced pressure. The obtained residue was dissolved in DCM (40 mL) and was washed with saturated NaHCO_3_ solution (35 mL) and brine solution (30 mL), dried over anhydrous NaSO_4_, filtered and concentered under reduced pressure. Flash column chromatography of the crude mixture on silica gel using cyclohexane, ethylacetate, and methanol as eluents (from CY:EA:MeOH 1:0:0 to CY:EA:MeOH 0:9:1) afforded the desired NBD-ceramide-C_6_CUM FRET probe **17** a as red-waxy solid.

Yield: 23 mg (43 %). R*_f_*: 0.35 (EA/MeOH 4:1, visualized with UV illumination at 366 nm). ^1^H NMR (CDCl_3_/CD_3_OD, 500 MHz, ppm): δ 8.45 (d, *J* = 8.2 Hz, 1H). 7.12 (s, 1H), 6.79 (s, 1H), 6.62 (t, *J* = 5.2 Hz, 1H), 6.28 (d, *J* = 8.8 Hz, 1H), 5.71 – 5.64 (m, 1H), 5.45 (dd, *J* = 7.2, 15.4 Hz, 1H), 4.09 (dt, *J* = 7.0, 13.7 Hz, 1H), 3.87 (dd, *J* = 5.3, 11.5 Hz, 1H), 3.64 – 3.74 (m, 2H), 3.20 (t, *J* = 6.9 Hz, 2H), 2.91 (t, *J* = 5.8 Hz, 2H), 2.54 (s, 3H), 2.22 (t, *J* = 7.4 Hz, 2H), 1.96–2.03 (m, 2H), 1.72 – 1.81 (m, 2H), 1.58 – 1.65 (m, 2H), 1.51 (dd, *J* = 7.3, 14.8 Hz, 2H), 1.40 – 1.49 (m, 2H), 1.23– 1.40 (m, 14H). ^13^C NMR (CDCl_3_/CD_3_OD, 125 MHz, ppm): δ 175.88, 168.96, 158.88, 153.36, 152.30, 146.63, 134.50, 130.94, 114.79, 112.44, 110.26, 103.90, 98.64, 73.59, 62.11, 56.52, 49.85, 42.30, 40.20, 36.97, 33.22, 30.44, 30.38, 30.25, 30.17, 30.10, 29.86, 27.95, 27.35, 26.46, 20.28. ESI-MS *m/z* calcd for C_39_H_50_N_6_O_11_Br [M+H]^+^ 857.27; observed 857.2.

**11-(2-(7-bromo-8-methy-6-oxo-6*H*-[1,3]dioxolo[4,5-*g*]chromen-2-yl)acetamido)-*N*-((2*S*,3*R*,*E*)-1,3-dihydroxy-14-((7-nitrobenzo[*c*][1,2,5]oxadiazol-4-yl)amino)tetradec-4-en-2-yl)undecane-amide (18, FRETcer2)**

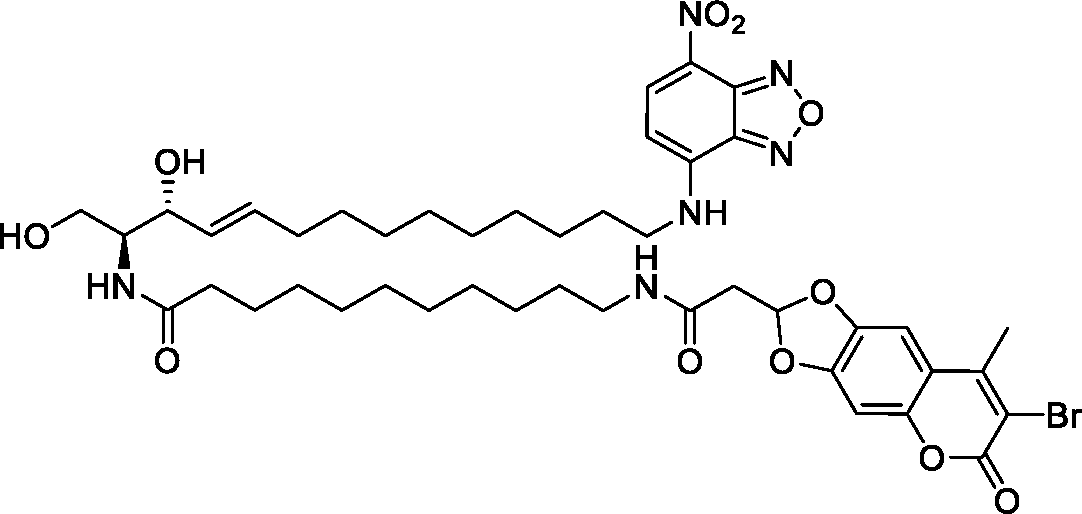

A stirred solution of coumarin-labeled C_11_-fatty acid **14** (41 mg, 0.077mmol) in dry DCM (2mL) under argon atmosphere at 0°C was treated with DMAP (38 mg, 0.31mmol), followed by *N*-hydroxysuccinimide (11 mg, 0.09mmol). The resulting reaction mixture was stirred at ambient temperature for 6h, before the solvent was removed under reduced pressure. The obtained residue was purified by flash column chromatography using ethylacetate as eluent (100%) to furnish the corresponding activated ester of the coumarin fatty acid which was used directly in the next step. The resultant activated acid was dissolved in dry DCM (2mL) under argon atmosphere and the obtained solution was cooled to 0°C. Subsequently, the solution was treated with DIPEA (41 µL, 0.23mmol), followed by a solution of NBD-labeled sphingosine **4** (27mg, 0.064 mmol) in dry DCM (1mL). After the resulting reaction mixture was stirred for 16h at ambient temperature, the solvent was removed under reduced pressure and the obtained residue was purified by flash column chromatography (from PE:EA:MeOH 1:0:0 to PE:EA:MeOH 0:10:1) to furnish NBD-ceramide-C_11_CUM FRET probe **18** a as red-waxy oil.

Yield: 17 mg (29 %). R*_f_*: 0.41 (EA/MeOH 5:1, visualized with UV illumination at 366 nm). ^1^H NMR (500 MHz, CDCl_3_, ppm) δ ^1^H NMR (500 MHz, CDCl_3_) δ 8.35 (s, 1H), 6.90 (d, *J* = 5.1 Hz, 1H), 6.67 (d, *J* = 4.0 Hz, 1H), 6.50 (d, *J* = 5.3 Hz, 1H), 6.06 (s, 1H), 5.59 (dt, *J* = 10.4, 6.1 Hz, 1H), 5.35 (dd, *J* = 14.8, 5.0 Hz, 1H), 4.03 (br.s, 1H), 3.52 (dd, *J* = 10.4, 7.0 Hz, 2H), 3.46 – 3.31 (m, 3H), 3.08 (dd, *J* = 11.9, 5.3 Hz, 2H), 2.77 (d, *J* = 5.0 Hz, 2H), 2.44 (s, *J* = 4.4 Hz, 3H), 2.11 – 2.04 (m, 2H), 1.96 – 1.86 (m, 2H), 1.69 – 1.61 (m, 2H), 1.51 – 1.42 (m, 2H), 1.36 – 1.11 (m, 26H). ^13^C NMR (126 MHz, CDCl_3_, ppm) δ 174.63, 167.21, 157.87, 151.79, 150.95, 148.58, 145.34, 135.30, 133.45, 128.98, 127.49, 113.56, 110.98, 109.41, 102.46, 97.69, 72.89, 61.23, 54.73, 39.34, 36.23, 31.98, 29.38, 29.20, 29.10, 29.07, 28.96, 28.89, 28.84, 26.72, 26.57, 25.54, 19.61. ESI-MS *m/z* calcd for C_44_H_60_N_6_O_11_Br [M+H]^+^ 927.35; observed 927.3.

***N*-((2*S*,3*R*,*E*)-1,3-dihydroxy-14-((2-oxo-2*H*-chromen-6-yl)oxy)-tetradec-4-en-2-yl)-6-((7 nitrobenzo[*c*][1,2,5]oxadiazol-4-yl)amino)-hexanamide (19, FRETcer3)**

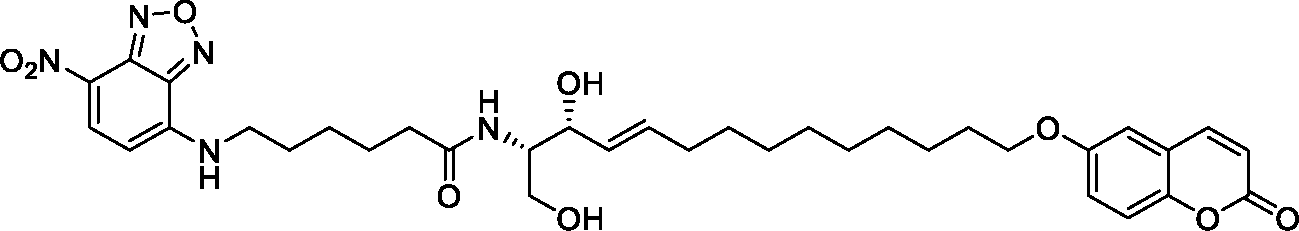

To a stirred solution of NBD-labelled C_6_-fatty acid **16** (36mg, 0.12mmol) in dry DCM (3 mL) under argon atmosphere at 0°C was added DIPEA (63µL, 0.36 mmol), followed by EDCI (30 mg, 0.16 mmol) and HOBt (22 mg, 0.16 mmol). The resulting mixture was stirred for 10 min at ambient temperature, before a solution of coumarin-labeled sphingosine **7** (40 mg, 0.1mmol) in dry DCM (1mL) was added. After the resulting reaction mixture was stirred for 14h at ambient temperature, the reaction was quenched with 0.1M HCl (20mL). The resulting mixture was extracted with DCM, and the combined organic extracts was dried over anhydrous Na_2_SO_4_, filtered, and concentrated. Flash column chromatography of resultant residue over silica gel (from PE:EA 1:0 to PE:EA 1:10) provided the desired FRET ceramide probe **19** as a faint red waxy-solid.

Yield: 37 mg (53 %). R*_f_*: 0.32 (EA/MeOH 5:1, visualized with UV illumination at 366 nm). ^1^H NMR (500 MHz, CDCl_3_, ppm) δ 8.37 (d, *J* = 7.7 Hz, 1H), 7.57 (d, *J* = 9.4 Hz, 1H), 7.29 (d, *J* = 8.6 Hz, 1H), 6.76 (dd, *J* = 8.6, 1.8 Hz, 1H), 6.58 (s, 1H), 6.16 (d, *J* = 9.4 Hz, 1H), 6.09 (s, 1H), 5.75 – 5.67 (m, 1H), 5.47 (dd, *J* = 15.3, 5.0 Hz, 1H), 4.28 (s, 1H), 3.94 (t, *J* = 6.4 Hz, 2H), 3.90 (d, *J* = 6.9 Hz, 1H), 3.66 (d, *J* = 7.9 Hz, 1H), 2.26 (m, 2H), 1.97 (dd, *J* = 13.1, 6.3 Hz, 2H), 1.80 – 1.66 (m, 6H), 1.46 (s, 2H), 1.41 – 1.34 (m, 2H), 1.31 – 1.17 (m, 12H). ^13^C NMR (126 MHz, CDCl_3_, ppm) δ 162.61, 161.68, 155.97, 144.40, 143.83, 136.82, 134.23, 128.92, 122.14, 113.14, 112.90, 112.50, 101.52, 74.58, 68.79, 62.33, 54.65, 36.20, 32.39, 29.48, 29.40, 29.28, 29.21, 28.98, 26.32, 25.96, 24.91. ESI-MS *m/z* calcd for C_35_H_45_N_5_O_9_ [M+H]^+^ 680.33; observed 680.3.

#### II. 2. Synthesis of NSC405020 analogues

**3,4-Dichloro-*N*-isopentylbenzenesulfonamide (ES_ACR01):**

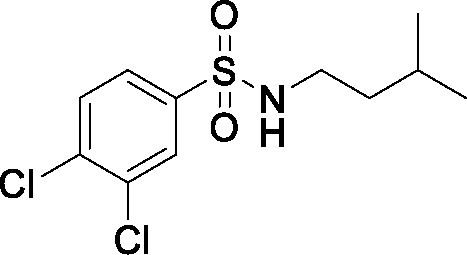

To a stirred solution of 3,4-Dichlorobenzenesulfonyl chloride (250 mg, 1.02 mmol) in acetonitrile (10 mL) at 0°C was added *iso*-pentylamine (360 µL, 3.05 mmol). After the resulting reaction mixture was stirred for 2h at ambient temperature, the reaction was quenched with 1M HCl (20 mL). The resulting mixture was extracted with ethylacetate (3×50mL), and the combined organic layers were washed with brine, dried over anhydrous NaSO_4_, filtered and concentered under reduced pressure. Purification of the resultant residue by flash column chromatography over silica gel using pet. ether and ethylacetate as eluents (20-25% ethylacetate in pet. ether) provided compound **ES_ACR01** as a white solid.

Yield: (77%). R*_f_*: 0.45 (PE/EA 4:1, visualized with UV illumination at 254 nm). ^1^H NMR (500 MHz, CDCl_3_, ppm) δ 7.95 (d, *J* = 2.1 Hz, 1H), 7.69 (dd, *J* = 8.4, 2.1 Hz, 1H), 7.60 (d, *J* = 8.4 Hz, 1H), 4.50 (t, *J* = 5.4 Hz, 1H), 3.04 – 2.95 (m, 2H), 1.58 (td, *J* = 13.4, 6.7 Hz, 1H), 1.37 (dd, *J* = 14.5, 7.1 Hz, 2H), 0.86 (d, *J* = 6.6 Hz, 3H). ^13^C NMR (75 MHz, CDCl_3_, ppm) δ 140.04, 133.90, 131.31, 129.20, 126.26, 41.75, 38.55, 25.54, 22.35. ESI-MS *m/z* calcd for C_11_H_16_NO_2_SCl_2_ [M+H]^+^ 296.02; observed 296.0.

***N*-Butyl-3,4-dichlorobenzenesulfonamide (ES_ACR02):**

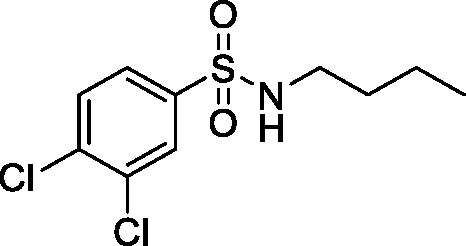

A stirred solution of 3,4-Dichlorobenzenesulfonyl chloride (250 mg, 1.02 mmol) in acetonitrile (10 mL) at 0°C was treated with *n*-butylamine (220 mg, 3.05 mmol). The resulting reaction mixture was allowed to stir for 3h at ambient temperature (as indicated by TLC analysis for complete reaction), before it was quenched by 1M HCl (50 mL). The resulting mixture was diluted with ethylacetate (60 mL) and the layers were separated. The aqueous layer was extracted with ethylacetate (3×60mL), and the combined organic layers were washed with brine, dried over anhydrous NaSO_4_, filtered and concentered under reduced pressure. The resultant crude mixture was purified by flash column chromatography over silica gel using ethylacetate and pet. ether (15-20% ethylacetate in pet. ether) to provide compound **ES_ACR02** as a white solid.

Yield: (84%). R*_f_*: 0.44 (PE/EA 4:1, visualized with UV illumination at 254 nm). ^1^H NMR (300 MHz, CDCl_3_, ppm) δ 7.95 (d, *J* = 2.1 Hz, 1H), 7.69 (dd, *J* = 8.4, 2.1 Hz, 1H), 7.60 (d, *J* = 8.4 Hz, 1H), 4.57 (t, *J* = 5.7 Hz, 1H), 2.98 (dd, *J* = 13.2, 6.9 Hz, 2H), 1.46 (ddd, *J* = 11.8, 5.1, 3.0 Hz, 2H), 1.35 – 1.26 (m, 2H), 0.87 (t, *J* = 7.3 Hz, 2H). ^13^C NMR (75 MHz, CDCl_3_, ppm) δ 140.08, 137.56, 133.89, 131.31, 129.19, 126.25, 43.18, 31.77, 19.79, 13.64. ESI-MS *m/z* calcd for C_10_H_14_NO_2_SCl_2_ [M+H]^+^ 282.01; observed 282.0.

***N*-(3,4-dichlorophenyl)-2-methylpentanamide (ES_ACR03):**

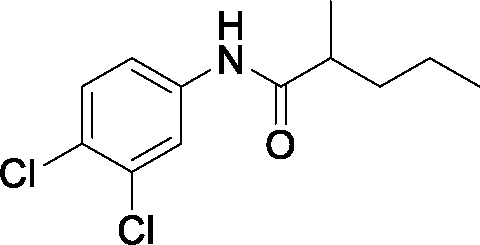

To a stirred solution of 2-methyl-valeric acid (430 mg, 3.7 mmol) in dry DMF (35 mL) under argon atmosphere was added DIPEA (1.6 mL, 9.3 mmol), followed by EDCI.HCl (890 mg, 4.65 mmol) and HOBt (630 mg, 4.65 mmol). The resulting mixture was stirred for 10 min at the same conditions, before it was treated with 3,4-dichloro-aniline (0.5 g, 3.1 mmol). After the resulting reaction mixture was stirred at ambient temperature for 16h (as indicated by TLC analysis), the reaction was quenched with 1M HCl (100 mL). The resulting mixture was extracted with ethylacetate (3×100mL), and the combined layers were washed with brine, dried over anhydrous NaSO_4_, filtered and concentered under reduced pressure. Flash column chromatography of the obtained residue over silica gel using ethylacetate and pet. ether as eluents (10-15% ethylacetate in pet. ether) provided compound **ES_ACR03** as a white solid.

Yield: (62%). R*_f_*: 0.44 (PE/EA 4:1, visualized with UV illumination at 254 nm). ^1^H NMR (300 MHz, CDCl_3_, ppm) δ 7.79 (s, 1H), 7.35 (d, *J* = 1.3 Hz, 2H), 7.29 (s, 1H), 2.40 – 2.26 (m, 1H), 1.78 – 1.64 (m, 1H), 1.51 – 1.31 (m, 3H), 1.22 (d, *J* = 6.8 Hz, 3H), 0.92 (t, *J* = 7.2 Hz, 3H). ^13^C NMR (75 MHz, CDCl_3_, ppm) δ 175.33, 137.55, 132.89, 130.58, 127.46, 121.69, 119.17, 42.59, 36.63, 20.78, 17.90, 14.17. ESI-MS *m/z* calcd for C_12_H_16_NOCl_2_ [M+H]^+^ 260.06; observed 260.1.

**Pentan-2-yl-3,4-dichlorobenzoate (ES_ACR04):**

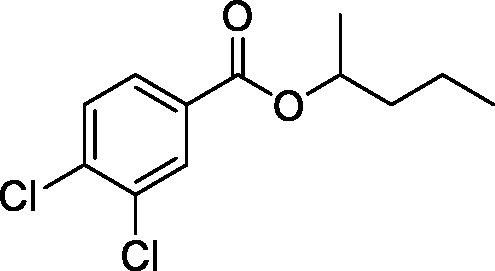

3,4-dichloro-benzoic acid (0.5 g, 2.62 mmol) under argon atmosphere at ambient temperature was treated with thionyl chloride (2 mL, 26.2 mmol). After the resulting reaction mixture was stirred at the same conditions for 2h, the unreacted thionyl chloride was removed under reduced pressure. The resulting 3-4-dichloro-benzoylchloride was suspended in 2-pentanol (4mL) under argon atmosphere and subsequently treated with sodium hydrogen carbonate (1.26 g, 15 mmol). The resulting reaction mixture was allowed to stir for 12h at ambient temperature (as judged by TLC analysis for complete esterification reaction). The solvent was removed under reduced pressure, and the obtained residue was portioned between water and ethylacetate. The layers were separated and the aqueous layer was extracted several times with ethylacetate. The combined organic layers were washed with brine, dried over anhydrous NaSO_4_, filtered and concentered. Flash column chromatography of the resultant crude product over silica gel using ethylacetate and pet. ether as eluents (5% ethylacetate in pet. ether) afforded the desired product **ES_ACR04** as colorless oil.

Yield: 525 mg (77%). R*_f_*: 0.44 (PE/EA 4:1, visualized with UV illumination at 254 nm). ^1^H NMR (300 MHz, MeOD, ppm) δ 8.09 (d, *J* = 1.8 Hz, 1H), 7.90 (dd, *J* = 8.4, 2.0 Hz, 1H), 7.66 (d, *J* = 8.3 Hz, 1H), 5.22 – 5.09 (m, 1H), 1.80 – 1.71 (m, 1H), 1.67 – 1.57 (m, 1H), 1.50 – 1.37 (m, 2H), 1.34 (d, *J* = 6.3 Hz, 3H), 0.96 (t, *J* = 7.3 Hz, 3H). ^13^C NMR (75 MHz, MeOD, ppm) δ 132.20, 131.99, 129.96, 73.78, 39.12, 20.21, 19.74, 14.20. ESI-MS *m/z* calcd for C_12_H_15_O_2_Cl_2_ [M+H]^+^ 261.04; observed 261.0.

***N*-Butyl-3,4-dichlorobenzamide (ES_ACR05):**

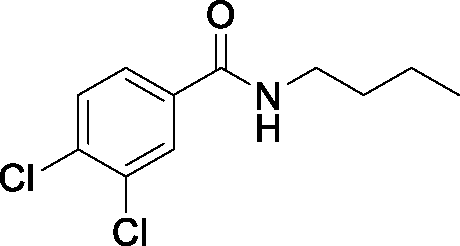

To a stirred solution of 3,4-dichloro-benzoic acid (0.5 g, 2.62 mmol) in dry DCM (20 mL) under argon atmosphere at ambient temperature was added DIPEA (1.3 mL, 7.86 mmol) followed by HOBt (420 mg, 3.1 mmol), EDCI.HCl (600 mg, 3.1 mmol), and n-butylamine (775 µL, 7.85 mmol). After the resulting reaction mixture was allowed to stir at the same conditions for 16h, the reaction was quenched with 1M HCl (100mL). The mixture was extracted with DCM (3×100mL), and the combined organic layers was washed with sat. NaHCO_3_ and brine, dried over anhydrous NaSO_4_, filtered and concentered under reduced pressure. Purification of the obtained crude product by flash column chromatography over silica gel using ethylacetate and per. ether as eluents (25% ethylacetate in pet. ether) afforded the desired product **ES_ACR05** as a white solid.

Yield: 340 mg (53%). R*_f_*: 0.45 (PE/EA 4:2, visualized with UV illumination at 254 nm). ^1^H NMR (300 MHz, CDCl_3_, ppm) δ 7.84 (d, *J* = 2.0 Hz, 1H), 7.58 (dd, *J* = 8.3, 2.0 Hz, 1H), 7.49 (d, *J* = 8.3 Hz, 1H), 6.16 (br.s, 1H), 3.44 (ddd, *J* = 13.0, 8.5, 4.9 Hz, 2H), 1.65 – 1.54 (m, 2H), 1.40 (dq, *J* = 9.9, 7.1 Hz, 2H), 0.95 (t, *J* = 7.3 Hz, 3H). ^13^C NMR (75 MHz, CDCl_3_, ppm) δ 165.46, 163.82, 135.87, 134.77, 133.17, 130.73, 129.22, 126.18, 40.17, 31.76, 20.27, 13.89. ESI-MS *m/z* calcd for C_11_H_14_NOCl_2_ [M+H]^+^ 246.04; observed 246.0.

**3,4-Dichloro-*N*-isopentylbenzamide (ES_ACR06):**

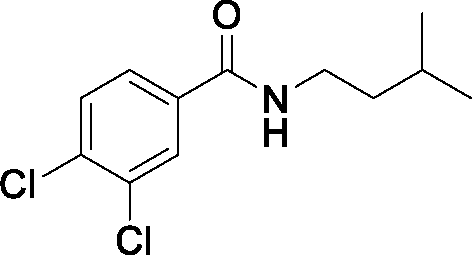

A solution of 3,4-dichloro-benzoic acid (0.5 g, 2.62 mmol) in dry DMF:DCM mixture (1:3, 20mL) under argon atmosphere at ambient temperature was treated with EDCI.HCl (600 mg, 3.1 mmol), followed by DIPEA (1.3 mL, 7.86 mmol) iso-pentylamine (920 µL, 7.85 mmol), and HOBt (420 mg, 3.1 mmol). The resulting reaction mixture was allowed to stir at the same conditions for 6h, and subsequently the reaction was quenched by 1M HCl (100mL). The obtained mixture was extracted with DCM (3×100mL) and the organic extracts were washed with brine, dried over anhydrous NaSO_4_, filtered and concentered. The obtained residue was purified by column chromatography over silica using ethaylacetate and pet ether as eluents (10-25% ethylacetate in pet. ether) to provide compound **ES_ACR06** as a white solid, tlc 8:2 0.45,

Yield: 460 mg (68%). R*_f_*: 0.47 (PE/EA 4:2, visualized with UV illumination at 254 nm). ^1^H NMR (300 MHz, CDCl_3_, ppm) δ 7.84 (d, *J* = 2.0 Hz, 1H), 7.57 (dd, *J* = 8.3, 2.0 Hz, 1H), 7.49 (d, *J* = 8.3 Hz, 1H), 6.10 (br.s, 1H), 3.50 – 3.41 (m, 2H), 1.75 – 1.60 (m, 1H), 1.50 (dd, *J* = 14.7, 7.1 Hz, 2H), 0.95 (d, *J* = 6.6 Hz, 6H). ^13^C NMR (75 MHz, CDCl_3_, ppm) δ 165.41, 135.87, 134.77, 133.17, 130.75, 129.22, 126.16, 38.76, 38.57, 26.09, 22.60. ESI-MS *m/z* calcd for C_12_H_16_NOCl_2_ [M+H]^+^ 260.06; observed 260.1.

**1-Butyl-3-(3,4-dichlorophenyl)-urea (ES_ACR07):**

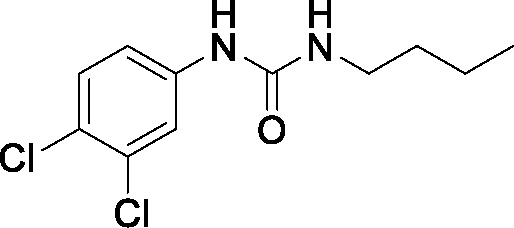

To a stirred solution of 3,4-dichloro-aniline (0.5 g, 3.1 mmol) in dry DCM (15 mL) under argon atmosphere at 0°C was added *n*-butyl isocyanate (740 µL, 6.8 mmol). The resulting reaction mixture was allowed to stir at 0°C for 1h and for additionally 16h at ambient temperature (as indicated by TLC analysis), before it was quenched by sat. NaHCO_3_ solution (100 mL). The resulting mixture was extracted by DCM (3×100mL). The combined extracts were washed with brine, dried over anhydrous NaSO_4_, filtered and concentered under reduced pressure. Flash column chromatography for the crude product using ethylacetate and pet ether as eluents (10-20% ethylacetate in pet. ether) furnished compound **ES_ACR07** as a white solid.

Yield: 380 mg (47%). R*_f_*: 0.43 (PE/EA 4:1, visualized with UV illumination at 254 nm). ^1^H NMR (300 MHz, CDCl_3_, ppm) δ 7.71 (br.s, 1H), 7.47 (d, *J* = 2.5 Hz, 1H), 7.29 (d, *J* = 1.1 Hz, 1H), 7.09 (dd, *J* = 8.7, 2.5 Hz, 1H), 3.21 (t, *J* = 7.0 Hz, 2H), 1.52 – 1.41 (m, 2H), 1.39 – 1.28 (m, 2H), 0.92 (t, *J* = 7.2 Hz, 3H). ^13^C NMR (75 MHz, CDCl_3_, ppm) δ 156.23, 138.52, 132.81, 130.56, 126.37, 121.59, 119.27, 40.27, 32.20, 20.18, 13.86. ESI-MS *m/z* calcd for C_11_H_15_N_2_OCl_2_ [M+H]^+^ 261.06; observed 2611.

***N*-(3,4-dichlorophenyl)-benzamide (ES_ACR08):**

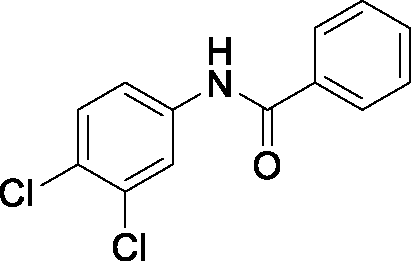

A stirred solution of benzoyl chloride (400 µL, 3.4 mmol) in dry DCM (30 mL) under argon atmosphere at 0°C was treated with DIPEA (1.6 mL, 9.3 mmol), followed by 3,4-dichloro-aniline (0.5g, 3.1 mmol). After the resulting reaction mixture was stirred at ambient temperature for 2h (as judged by TLC analysis), the reaction was quenched by 1M HCl (100mL). The obtained mixture was extracted with DCM, and the combined organic extracts were dried over anhydrous Na_2_SO_4_, filtered, and concentrated. Purification of the crude residue by column chromatography over silica gel using ethylacetate and pet ether as eluents (0-5% ethylacetate in pet. ether) provided compound **ES_ACR08** as a white solid.

Yield: 720 mg (88%). R*_f_*: 0.44 (PE/EA 4:1, visualized with UV illumination at 254 nm). ^1^H NMR (500 MHz, CDCl_3_, ppm) δ 7.97 (br.s, 1H), 7.88 (d, *J* = 2.5 Hz, 1H), 7.83 (dd, *J* = 8.3, 1.2 Hz, 2H), 7.58 – 7.53 (m, 1H), 7.50 – 7.45 (m, 3H), 7.39 (d, *J* = 8.7 Hz, 1H)., ppm ESI-MS *m/z* calcd for C_13_H_10_NOCl_2_ [M+H]^+^ 266.01; observed 266.0.

***N*-(3,4-dichlorophenyl)-4-heptylbenzamide (ES_ACR09):**

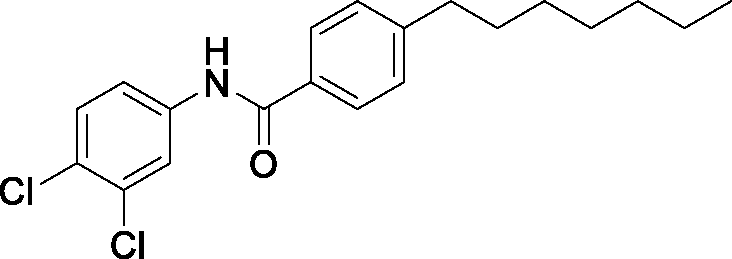

4-heptylbenzoic acid (755 mg, 3.7 mmol) was treated with thionyl chloride (3 mL, 37 mmol) under argon atmosphere. After the resulting mixture was stirred at the same conditions for 2h, the excess thionyl chloride was removed under vacuum to afford the corresponding acid chloride. The obtained acid chloride was dissolved in dry DCM (30 mL) under argon atmosphere and the resulting solution was treated with DIPEA (1.6 mL, 9.3 mmol), followed by 3,4-dichloro-aniline (0.5g, 3.1 mmol). The reaction mixture was stirred for 3h at ambient temperature (as indicated by TLC analysis), before it was quenched by 1M HCl (100mL). The mixture was extracted with DCM and the combined organic layers were washed with brine, dried over anhydrous Na_2_SO_4_, filtered, and concentrated. The obtained residue was purified by column chromatography over silica gel using ethylacetate and pet ether as eluents (0-5% ethylacetate in pet. ether) to provide compound **ES_ACR09** as a white solid.

Yield: 745 mg (66%). R*_f_*: 0.51 (PE/EA 5:1, visualized with UV illumination at 254 nm). ^1^H NMR (500 MHz, CDCl_3_, ppm) δ 7.88 (d, *J* = 2.4 Hz, 1H), 7.76 (d, *J* = 8.3 Hz, 2H), 7.47 (dd, *J* = 8.7, 2.5 Hz, 1H), 7.39 (d, *J* = 8.7 Hz, 1H), 7.28 (d, *J* = 8.3 Hz, 2H), 2.70 – 2.63 (m, 2H), 1.69 – 1.56 (m, 2H), 1.36 – 1.30 (m, 4H), 1.30 – 1.23 (m, 4H), 0.88 (t, *J* = 7.0 Hz, 3H). ^13^C NMR (126 MHz, CDCl_3_, ppm) δ 165.83, 148.06, 137.68, 133.01, 131.69, 130.68, 129.07, 127.72, 127.20, 121.96, 119.49, 36.02, 31.92, 31.31, 29.34, 29.26, 22.78, 14.22. ESI-MS *m/z* calcd for C_20_H_24_NOCl_2_ [M+H]^+^ 364.12; observed 364.1.

***N*-(3,4-dichlorophenyl)-2,3,4,5,6-pentafluorobenzamide (ES_ACR10):**

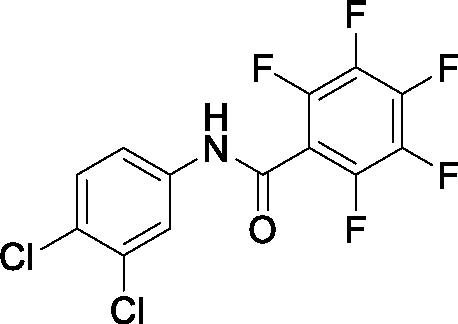

To a stirred solution of pentafluorobenzoic acid (755 mg, 3.7 mmol) in dry DMF (40 mL) under argon atmosphere at ambient temperature was added DIPEA (1.6 mL, 9.3 mmol), followed by EDCI.HCl (890 mg, 4.65 mmol), HOBt (630 mg, 4.65 mmol) and 3,4-dichloro-aniline (0.5g, 3.1 mmol). After the reaction mixture was stirred for 16h at the same conditions, the reaction was quenched with 1M HCl (100mL). The resulting mixture was extracted with ethylacetate (3×100) and the combined extracts were washed with brine, dried over anhydrous Na_2_SO_4_, filtered, and concentrated. Flash column chromatography for the obtained residue over silica gel using ethylacetate and pet ether as eluents (0-5% ethylacetate in pet. ether) furnished compound **ES_ACR10** as a pale-yellow solid.

Yield: 515 mg (47%). R*_f_*: 0.38 (PE/EA 5:1, visualized with UV illumination at 254 nm). ^1^H NMR (500 MHz, CDCl_3_, ppm) δ 7.83 (d, *J* = 2.2 Hz, 1H), 7.80 (br.s, 1H), 7.44 (d, *J* = 8.7 Hz, 1H), 7.41 (dd, *J* = 8.7, 2.3 Hz, 1H). ^13^C NMR (126 MHz, CDCl_3_, ppm) δ 155.55, 136.10, 133.37, 130.95, 129.36, 122.25, 119.65. ESI-MS *m/z* calcd for C_13_H_5_NOF_5_Cl_2_ [M+H]^+^ 355.97; observed 356.0.

***N*-(3,4-dichlorophenyl)-oleamide (ES_ACR11):**

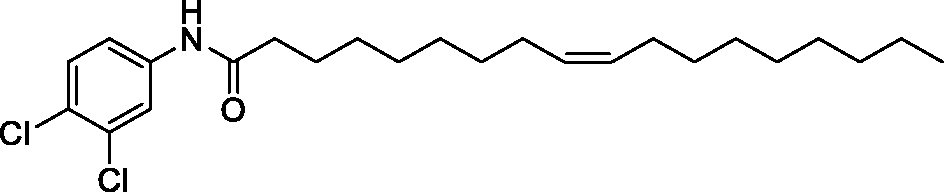

To a stirred solution of oleoyl chloride (1 mL, 3.4 mmol) in dry DCM (30 mL) under argon atmosphere at 0°C was added DIPEA (1.6 mL, 9.3 mmol), followed by 3,4-dichloro-aniline (0.5g, 3.1 mmol). The reaction mixture was stirred for 3h at the same conditions (as judged by TLC analysis), and subsequently quenched by 1M HCl. The obtained mixture was extracted with DCM and the combined organic layers were washed with brine, dried over anhydrous Na_2_SO_4_, filtered and concentrated. Purification of the obtained crude residue over silica gel using ethylacetate and pet ether as eluents (0-3% ethylacetate in pet. ether) afforded compound **ES_ACR11** as a white solid.

Yield: 740 mg (58%). R*_f_*: 0.58 (PE/EA 4:1, visualized with UV illumination at 254 nm). ^1^H NMR (500 MHz, CDCl_3_, ppm) δ 7.76 (d, *J* = 1.7 Hz, 1H), 7.36 (d, *J* = 8.6 Hz, 1H), 7.33 (dd, *J* = 8.8, 2.1 Hz, 1H), 7.22 (br.s, 1H), 5.40 – 5.30 (m, 2H), 2.34 (t, *J* = 7.6 Hz, 2H), 2.07 – 1.93 (m, 4H), 1.74 – 1.67 (m, 2H), 1.41 – 1.23 (m, 14H), 0.88 (t, *J* = 7.0 Hz, 3H). ^13^C NMR (126 MHz, CDCl_3_, ppm) δ 171.59, 137.53, 132.93, 130.62, 130.21, 129.83, 121.56, 119.04, 37.86, 32.05, 29.91, 29.84, 29.67, 29.47, 29.46, 29.40, 29.34, 29.24, 27.38, 27.30, 25.58, 22.83, 14.26. ESI-MS *m/z* calcd for C_24_H_38_NOCl_2_ [M+H]^+^ 426.23; observed 426.2.

***N*-(3,4-dichlorophenyl)-dodecanamide (ES_ACR12):**

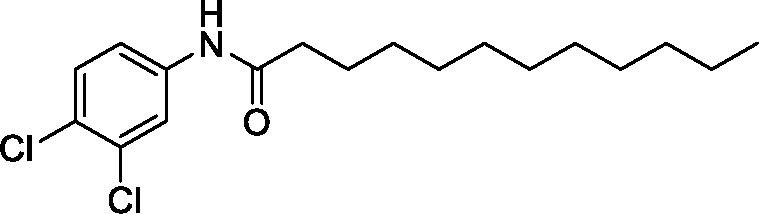

A stirred solution of lauryl chloride (790 µL, 3.4 mmol) in dry DCM (35 mL) under argon atmosphere was treated with DIPEA (1.6 mL, 9.3 mmol), followed by 3,4-dichloro-aniline (0.5g, 3.1 mmol). After stirring at ambient temperature for 2h, the reaction was quenched with 1M HCl and the resultant mixture was extracted with DCM (3×80mL). The organic extracts were combined and washed with brine, dried over anhydrous Na_2_SO_4_, and filtered. After the solvent was removed under reducing pressure, the residue was purified by flash column chromatography using ethylacetate and pet ether as eluents (0-5% ethylacetate in pet. ether) to give compound **ES_ACR12** as a white solid.

Yield: 730 mg (69%). R*_f_*: 0.54 (PE/EA 4:1, visualized with UV illumination at 254 nm). ^1^H NMR (500 MHz, CDCl_3_, ppm) δ 7.77 (d, *J* = 1.3 Hz, 1H), 7.35 (d, *J* = 8.6 Hz, 1H), 7.33 (dd, *J* = 9.0, 1.8 Hz, 1H), 2.34 (t, *J* = 7.6 Hz, 2H), 1.74 – 1.67 (m, 2H), 1.40 – 1.22 (m, 18H), 0.88 (t, *J* = 6.9 Hz, 3H). ^13^C NMR (126 MHz, CDCl_3_, ppm) δ 171.67, 137.54, 132.92, 130.61, 127.46, 121.57, 119.06, 37.88, 32.05, 29.74, 29.60, 29.49, 29.47, 29.37, 25.60, 22.82, 14.25. ESI-MS *m/z* calcd for C_18_H_28_NOCl_2_ [M+H]^+^ 344.15; observed 344.1.

***N*-(3,4-dichlorophenyl)-2-methylpentanamide (ES_ACR13):**

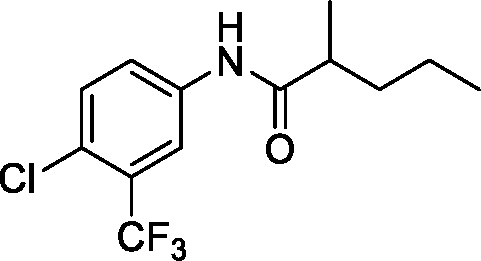

2-methyl-valeric acid (430 mg, 3.7 mmol) was treated with thionyl chloride (3 mL, 37 mmol) under argon atmosphere and the resulting reaction mixture was stirred at the same conditions for 2h. Subsequently, the excess thionyl chloride was removed under reduced pressure to afford the corresponding acid chloride as pale-yellow residue, which was used directly in the next step. After the obtained acid chloride was dissolved in dry DCM (35 mL) under argon atmosphere, DIPEA (1.6 mL, 9.3 mmol) was added at ambient temperature, followed by 4-chloro-3-(trifluoromethyl)aniline (565 mg, 2.9 mmol). The resulting reaction mixture was stirred at the same conditions for 6h, before it was quenched with 1M HCl. The obtained mixture was extracted with DCM, and the combined organic extracts were washed with brine, dried over anhyd. Na_2_SO_4_, filtered and concentrated. Flash column chromatography for the obtained crude product over silica gel (10-20% ethylacetate in pet. ether) yielded compound **ES_ACR13** as a white solid.

Yield: 325 mg (43%). R*_f_*: 0.39 (PE/EA 4:1, visualized with UV illumination at 254 nm). ^1^H NMR (500 MHz, CDCl_3_, ppm) δ 7.87 (d, *J* = 2.3 Hz, 1H), 7.75 (dd, *J* = 8.7, 2.3 Hz, 1H), 7.42 (d, *J* = 8.6 Hz, 2H), 2.37 (dt, *J* = 14.0, 6.8 Hz, 1H), 1.76 – 1.67 (m, 1H), 1.50 – 1.42 (m, 1H), 1.40 – 1.32 (m, 2H), 1.23 (d, *J* = 6.8 Hz, 3H), 0.92 (t, *J* = 7.3 Hz, 3H). ^13^C NMR (126 MHz, CDCl_3_, ppm) δ 175.52, 136.95, 132.08, 129.02, 128.77, 126.87, 123.90, 123.77, 121.60, 118.92, 118.88, 42.60, 36.61, 20.78, 17.86, 14.15. ESI-MS *m/z* calcd for C_12_H_16_NOCl_2_ [M+H]^+^ 260.06; observed 260.1.

***N*-(4-cyanophenyl)-cinnamamide (ES_ACR14):**

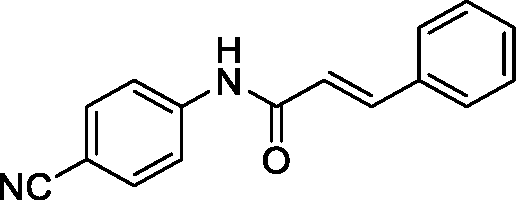

To a cinnamic acid (550 mg, 3.7 mmol) under argon atmosphere at ambient temperature was added thionyl chloride (3 mL, 37 mmol). After the resulting reaction mixture was stirred for 2h at the same conditions, the unreacted thionyl chloride was removed under reduced pressure to afford the corresponding acid chloride. The obtained acid chloride was dissolved in dry DCM (30 mL) under argon atmosphere and subsequently treated with DIPEA (1.6 mL, 9.3 mmol), followed by 4-aminobenzonitrile (365 mg, 3.1 mmol). The reaction mixture was allowed to stir at the same conditions for 3h, before it was quenched with 1M HCl (100mL). The resultant mixture was extracted with DCM (3×80mL), and the collected extracts were dried over anhyd. Na_2_SO_4_, filtered and concentrated under reduced pressure. Purification of obtained residue by flash column chromatography using ethylacetate and pet. ether as eluents (15-20% ethylacetate in pet. ether) provided compound **ES_ACR14** as a white solid.

Yield: 530 mg (79%). R*_f_*: 0.56 (PE/EA 4:2, visualized with UV illumination at 254 nm).^1^H NMR (500 MHz, MeOD, ppm) δ 7.88 – 7.85 (m, 1H), 7.70 (d, *J* = 14.1 Hz, 1H), 7.68 (dd, *J* = 5.6, 3.6 Hz, 1H), 7.60 (dd, *J* = 7.5, 2.0 Hz, 1H), 7.44 – 7.39 (m, 2H), 6.78 (d, *J* = 15.7 Hz, 1H). ^13^C NMR (126 MHz, MeOD, ppm) δ 166.84, 144.58, 143.92, 136.02, 134.25, 131.31, 130.05, 129.09, 121.68, 120.97, 119.86, 107.62. ESI-MS *m/z* calcd for C_16_H_13_N_2_O [M+H]^+^ 249.10; observed 249.1.

***N*-(3,4-dichlorophenyl)-4-isopropylcyclohexane-1-carboxamide (ES_ACR15):**

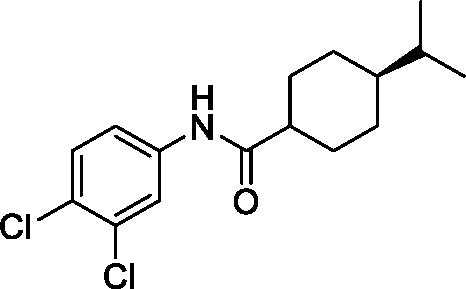

To a stirred solution of *trans*-4-*iso*-propylcyclohexane-1-carboxylic acid (630 mg, 3.7 mmol) in dry DMF under argon atmosphere was added *N*-methyl-2-pyrrolidone (NMP) (15 mL, 9.3 mmol), followed by HOBt (630 mg, 4.65 mmol), *N*,*N’*-diisopropylcarbodiimide (DIC) (700 µL, 4.65 mmol), and 3,4-dichloro-aniline (0.5 g, 3.1 mmol). The resulting reaction mixture was allowed to stir at ambient temperature for 16h (as indicated by TLC analysis), and subsequently the reaction was quenched with 1M HCl (80 mL). The obtained mixture was extracted with ethylacetate and the organic extracts were combined and washed with brine. The combined extracts were dried over anhydrous NaSO_4_, filtered and concentered under reduced pressure. The crude product was purified by flash column chromatography using ethylacetate and pet. ether as eluent (10-20% ethylacetate in pet. ether) afforded compound **ES_ACR15** as a white solid.

Yield: 495 mg (51%). R*_f_*: 0.44 (PE/EA 4:1, visualized with UV illumination at 254 nm).^1^H NMR (500 MHz, CDCl_3_, ppm) δ 7.79 (s, 1H), 7.31 (d, *J* = 30.8 Hz, 2H), 2.15 (tt, *J* = 12.1, 3.4 Hz, 1H), 2.02 (dd, *J* = 25.4, 8.7 Hz, 2H), 1.86 – 1.77 (m, 2H), 1.53 (qd, *J* = 12.7, 3.0 Hz, 1H), 1.45 – 1.39 (m, 1H), 1.15 – 1.06 (m, 1H), 1.04 – 0.96 (m, 2H), 0.88 – 0.85 (m, 6H). ^13^C NMR (126 MHz, CDCl_3_, ppm) δ 180.98, 174.74, 137.67, 132.89, 130.58, 127.37, 121.61, 119.09, 46.86, 43.38, 43.31, 43.25, 32.89, 29.87, 29.21, 29.04, 28.94, 19.86. ESI-MS *m/z* calcd for C_16_H_22_NOCl_2_ [M+H]^+^ 314.11; observed 314.1.

***N*-(3,4-dichlorophenyl)-4’-(trifluoromethyl)-[1,1’-biphenyl]-2-carboxamide (ES_ACR16):**

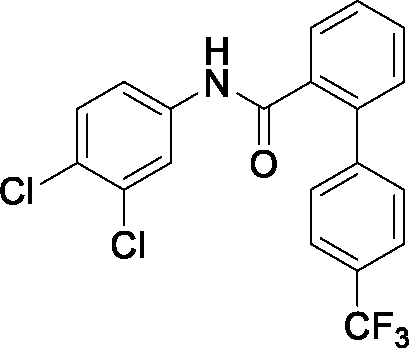

4′-(Trifluoromethyl)-2-biphenylcarboxylic acid (980 mg, 3.7 mmol) under argon atmosphere was treated with thionyl chloride (3 mL, 37 mmol). After the resulting reaction solution was allowed to stir at ambient temperature for 3h, the excess thionyl chloride was removed under vacuum. The resultant residue was dissolved in dry DCM (30mL) under argon atmosphere and the obtained solution was treated with DIPEA (1.6 mL, 9.3 mmol), followed by 3,4-dichloroaniline (500 mg, 3.1 mmol). The resulting reaction mixture was stirred for 2h at ambient temperature (as indicated by TLC analysis), before it was quenched with 1M HCl (100mL). The mixture was extracted with DCM (80mL) and the combined organic extracts was washed with brine, dried over anhydrous NaSO_4_, filtered and concentered under reduced pressure. Flash column chromatography over silica gel for the resultant crude residue (5-10% ethylacetate in pet. ether) provided compound **ES_ACR16** as a white solid.

Yield: 810 mg (64%). R*_f_*: 0.39 (PE/EA 9:1, visualized with UV illumination at 254 nm). ^1^H NMR (500 MHz, CDCl_3_, ppm) δ 7.78 (d, *J* = 6.9 Hz, 1H), 7.69 (d, *J* = 8.1 Hz, 2H), 7.60 (td, *J* = 7.6, 1.3 Hz, 1H), 7.56 (d, *J* = 8.0 Hz, 2H), 7.52 (td, *J* = 7.6, 1.3 Hz, 1H), 7.48 (d, *J* = 2.3 Hz, 1H), 7.44 (d, *J* = 7.5 Hz, 1H), 7.29 (d, *J* = 8.7 Hz, 1H), 6.96 (br.s, 1H), 6.92 (dd, *J* = 8.7, 2.4 Hz, 1H). ^13^C NMR (126 MHz, CDCl_3_, ppm) δ 167.14, 143.58, 138.48, 136.88, 135.23, 133.05, 131.36, 130.63, 130.56, 129.28, 129.23, 128.83, 128.20, 126.02, 125.99, 125.16, 123.00, 121.65, 119.04. ESI-MS *m/z* calcd for C_20_H_13_NOF_3_Cl_2_ [M+H]^+^ 410.03; observed 410.0.

**(*E*)-2-(4-(tetradecyloxy)benzylidene)hydrazine-1-carboximidamide (ES_ACR17)**

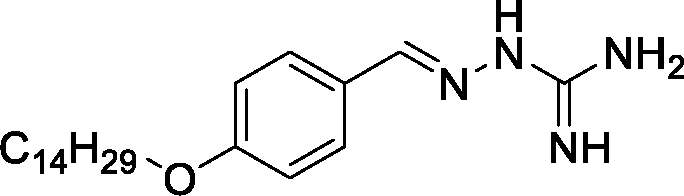

The titled compound was synthesized in 4-steps following our previously reported route.^6^

**Phenyl (3,4-dichlorophenyl)carbamate (ES_ACR18):**

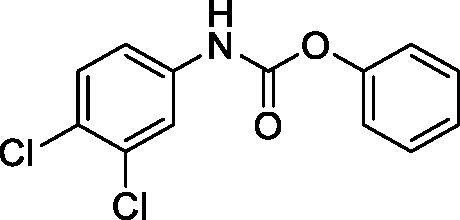

(*General procedure for carbamate formation*): To a stirred solution of 3,4-dichloro-aniline (500 mg, 3.1 mmol) in dry DCM (8 mL) under argon atmosphere at 0°C was added triethylamine (620 µL, 4.4 mmol) followed by phenyl chloroformate (470 µL, 3.7 mmol). The resulting reaction mixture was allowed to stir at the same conditions for 1h and for additional 3h at ambient temperature (as monitored by TLC analysis for complete carbamate formation). The reaction mixture was cooled again to 0°C, subsequently diluted with DCM (50 mL) and quenched with 1M HCl (50 mL). The layers were separated, and the aqueous layer was extracted again with DCM (2×50mL). The organic layers were combined and washed with brine, dried over anhydrous NaSO_4_, filtered and concentered under reduced pressure. Purification of the obtained crude mixture by flash column chromatography over silica gel using ethylacetate and pet. ether as eluents (5-10% ethylacetate in pet. ether) yielded the desired product **ES_ACR18** as a white solid.

Yield: 550 mg (63%). R*_f_*: 0.51 (PE/EA 4:1, visualized with UV illumination at 254 nm). ^1^H NMR (500 MHz, CDCl_3_, ppm) δ 7.66 (br.s, 1H), 7.40 (ddd, *J* = 12.4, 7.4, 5.4 Hz, 3H), 7.29 – 7.24 (m, 2H), 7.18 (dd, *J* = 9.8, 2.2 Hz, 2H), 7.00 (br.s, 1H). ^13^C NMR (126 MHz, CDCl_3_, ppm) δ 151.52, 150.43, 137.04, 133.19, 130.82, 129.65, 127.39, 126.16, 121.65, 120.63, 118.13. ESI-MS *m/z* calcd for C_13_H_10_NO_2_Cl_2_ [M+H]^+^ 282.01; observed 282.0.

**Phenyl (4-chloro-3-(trifluoromethyl)phenyl)carbamate (ES_ACR19):**

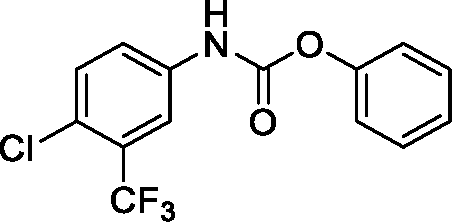

The titled compound was synthesized following the general procedure described for compound **ESCA18**. Accordingly, 4-chloro-3-(trifluoromethyl)aniline (580 mg, 3.1 mmol) was reacted with phenyl chloroformate (470 µL, 3.7 mmol) in presence of triethylamine (620 µL, 4.4 mmol) to afford compound **ESCA19** as a faint brown solid after purification by flash column chromatography (5-10% ethylacetate in pet. ether).

Yield: 710 mg (72%). R*_f_*: 0.43 (PE/EA 9:1, visualized with UV illumination at 254 nm). ^1^H NMR (500 MHz, CDCl_3_, ppm) δ 7.81 (d, *J* = 1.6 Hz, 1H), 7.60 (dd, *J* = 8.7, 2.1 Hz, 1H), 7.47 – 7.39 (m, 3H), 7.29 – 7.24 (m, 1H), 7.18 (dt, *J* = 9.0, 1.9 Hz, 2H), 7.13 (br.s, 1H). ^13^C NMR (126 MHz, CDCl_3_, ppm) δ 151.63, 150.37, 136.47, 132.31, 129.70, 129.29, 129.04, 126.26, 123.72, 122.77, 121.64, 117.97. ESI-MS *m/z* calcd for C_14_H_10_NO_2_F_3_Cl [M+H]^+^ 316.03; observed 316.0.

**Phenyl octadecylcarbamate (ES_ACR20):**

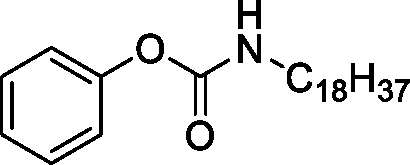

The titled compound was synthesized following the general procedure described for compound **ESCA18**. Accordingly, octadecylamine (950 mg, 3.1 mmol) was reacted with phenyl chloroformate (470 µL, 3.7 mmol) in presence of triethylamine (620 µL, 4.4 mmol) to afford compound **ESCA20** as a white solid after purification by flash column chromatography (5-10% ethylacetate in pet. ether).

Yield: 830 mg (69%). R*_f_*: 0.53 (PE/EA 9:1, visualized with UV illumination at 254 nm). ^1^H NMR (500 MHz, CDCl_3_, ppm) δ 7.37 – 7.33 (m, 2H), 7.20 – 7.16 (m, 1H), 7.12 (dd, *J* = 8.6, 0.9 Hz, 2H), 3.33 (d, *J* = 4.6 Hz, 1H), 3.26 (dd, *J* = 13.3, 6.9 Hz, 1H), 1.57 (dt, *J* = 14.6, 7.2 Hz, 2H), 1.38 – 1.23 (m, 26H), 0.88 (t, *J* = 7.0 Hz, 3H). ^13^C NMR (126 MHz, CDCl_3_, ppm) δ 154.73, 151.25, 129.39, 129.33, 125.33, 125.14, 121.94, 121.73, 41.44, 32.07, 29.98, 29.85, 29.80, 29.74, 29.69, 29.51, 29.42, 26.91, 22.84, 14.26. ESI-MS *m/z* calcd for C_25_H_44_NO_2_ [M+H]^+^ 390.34; observed 390.3.

**Phenyl undecylcarbamate (ES_ACR21):**

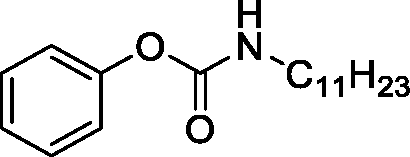

Following the general procedure described for compound **ES_ACR18**, the titled compound **ES_ACR21** was synthesized starting from undecylamine (660 µL, 3.1 mmol) and obtained as a white solid after flash column chromatography (10-15% ethylacetate in pet. ether).

Yield: 700 mg (78%). R*_f_*: 0.49 (PE/EA 4:1, visualized with UV illumination at 254 nm). ^1^H NMR (500 MHz, CDCl_3_, ppm) δ 7.37 – 7.33 (m, 2H), 7.19 (t, *J* = 7.4 Hz, 1H), 7.13 (d, *J* = 7.7 Hz, 1H), 5.01 (br.s, 1H), 3.32 (d, *J* = 4.5 Hz, 1H), 3.26 (dd, *J* = 13.3, 6.9 Hz, 1H), 1.60 – 1.53 (m, 2H), 1.39 – 1.24 (m, 16H), 0.89 (t, *J* = 7.0 Hz, 3H). ^13^C NMR (126 MHz, CDCl_3_, ppm) δ 154.72, 151.26, 129.38, 129.32, 125.31, 125.13, 121.93, 121.73, 41.43, 32.05, 29.98, 29.74, 29.73, 29.68, 29.47, 29.41, 26.90, 22.82, 14.25. ESI-MS *m/z* calcd for C_18_H_30_NO_2_ [M+H]^+^ 292.23; observed 292.2.

**Phenyl pentylcarbamate (ESCA22):**

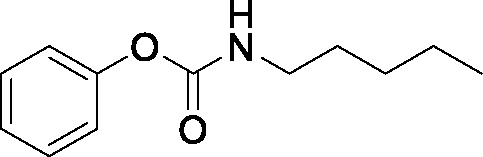

Following the general procedure described for compound **ES_ACR18**, the titled compound **ES_ACR22** was synthesized starting from *n*-pentylamine (360 µL, 3.1 mmol) and obtained as a pale-yellow solid after flash column chromatography (10-20% ethylacetate in pet. ether).

Yield: 390 mg (61%). R*_f_*: 0.46 (PE/EA 4:1, visualized with UV illumination at 254 nm). ^1^H NMR (500 MHz, CDCl_3_, ppm) δ 7.35 (t, *J* = 7.9 Hz, 2H), 7.19 (t, *J* = 7.4 Hz, 1H), 7.13 (d, *J* = 7.7 Hz, 2H), 5.02 (br.s, 1H), 3.26 (dd, *J* = 13.3, 6.9 Hz, 2H), 1.62 – 1.53 (m, 2H), 1.39 – 1.30 (m, 4H), 0.92 (t, *J* = 7.0 Hz, 3H). ^13^C NMR (126 MHz, CDCl_3_, ppm) δ 154.73, 151.25, 129.38, 125.32, 121.73, 41.39, 29.66, 29.03, 22.47, 14.12. ESI-MS *m/z* calcd for C_12_H_18_NO_2_ [M+H]^+^ 208.13; observed 208.1.

**Phenyl (4-acetylphenyl)-carbamate (ES_ACR23):**

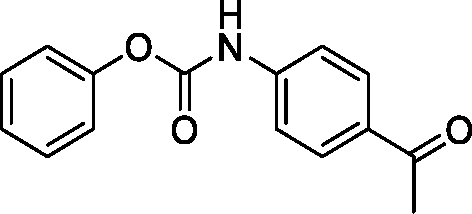

Following the general procedure described for compound **ES_ACR18**, the titled compound **ES_ACR23** was synthesized starting from 4-amino-acetophenone (400 mg, 3.1 mmol) and obtained as a pale yellow solid after flash column chromatography (5-15% ethylacetate in pet. ether).

Yield: 420 mg (53%). R*_f_*: 0.43 (PE/EA 5:1, visualized with UV illumination at 254 nm). ^1^H NMR (500 MHz, CDCl_3_, ppm) δ 8.06 (s, 1H), 7.74 (d, *J* = 7.7 Hz, 1H), 7.70 (d, *J* = 8.0 Hz, 1H), 7.45 – 7.38 (m, 3H), 7.32 (br.s, 1H), 7.27 – 7.23 (m, 1H), 7.20 (dd, *J* = 8.6, 1.1 Hz, 2H), 2.60 (s, 3H). ^13^C NMR (126 MHz, CDCl_3_, ppm) δ 198.01, 151.89, 150.55, 138.16, 138.09, 129.62, 126.03, 123.87, 123.34, 121.74, 118.54, 26.83. ESI-MS *m/z* calcd for C_15_H_14_NO_3_ [M+H]^+^ 256.09; observed 256.1.

**Methyl (phenoxycarbonyl)-*D*-phenylalaninate (ES_ACR24):**

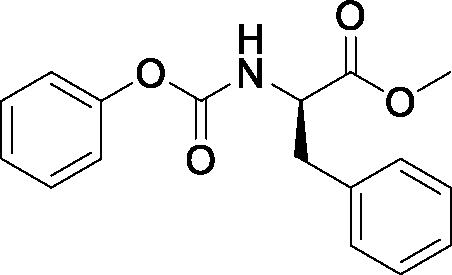

Following the general procedure described for compound **ES_ACR18**, the titled compound **ES_ACR24** was synthesized starting from *D*-phenylalanine methylester (600 mg, 3.1 mmol) and obtained as a white waxy solid after flash column chromatography (15-25% ethylacetate in pet. ether).

Yield: 750 mg (81%). R*_f_*: 0.47 (PE/EA 4:1, visualized with UV illumination at 254 nm). ^1^H NMR (500 MHz, CDCl_3_, ppm) δ 7.34 (dd, *J* = 14.7, 7.7 Hz, 4H), 7.29 (d, *J* = 7.2 Hz, 1H), 7.21 (d, *J* = 7.4 Hz, 1H), 7.17 (d, *J* = 7.1 Hz, 1H), 7.10 (d, *J* = 7.7 Hz, 2H), 5.51 (d, *J* = 8.0 Hz, 1H), 4.72 (dt, *J* = 8.2, 5.9 Hz, 1H), 3.76 (s, 3H), 3.22 (dd, *J* = 13.9, 5.8 Hz, 1H), 3.15 (dd, *J* = 13.9, 6.1 Hz, 1H). ^13^C NMR (126 MHz, CDCl_3_, ppm) δ 171.91, 154.04, 150.95, 135.72, 129.45, 129.44, 128.84, 127.41, 125.60, 121.64, 55.04, 52.58, 38.32. ESI-MS *m/z* calcd for C_17_H_18_NO_4_ [M+H]^+^ 300.12; observed 300.1.

**Phenyl (3,4-dimethoxyphenethyl)-carbamate (ES_ACR25):**

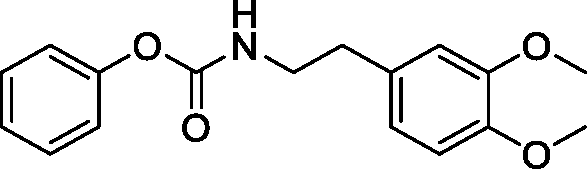

Following the general procedure described for compound **ES_ACR18**, the titeled compound **ES_ACR25** was synthesized strarting from 3,4-dimethoxy-phenylethylamine (522 µL, 3.1 mmol) and obtained as a white solid after flash column chromatography (10-15% ethylacetate in pet. ether).

Yield: 610 mg (66%). R*_f_*: 0.46 (PE/EA 5:1, visualized with UV illumination at 254 nm). ^1^H NMR (500 MHz, CDCl_3_, ppm) δ 7.35 (t, *J* = 7.9 Hz, 2H), 7.19 (t, *J* = 7.4 Hz, 1H), 7.10 (d, *J* = 7.6 Hz, 2H), 6.84 (d, *J* = 8.1 Hz, 1H), 6.79 – 6.74 (m, 2H), 5.05 (br.s, 1H), 3.88 (d, *J* = 4.8 Hz, 6H), 3.51 (dd, *J* = 13.2, 6.8 Hz, 2H), 2.84 (t, *J* = 7.0 Hz, 2H). ^13^C NMR (126 MHz, CDCl_3_, ppm) δ 154.69, 151.14, 149.26, 147.95, 131.17, 129.41, 125.42, 121.68, 120.85, 112.10, 111.59, 56.09, 56.02, 42.60, 35.63. ESI-MS *m/z* calcd for C_17_H_20_NO_4_ [M+H]^+^ 302.14; observed 302.1.

**Phenyl (furan-2-ylmethyl)-carbamate (ES_ACR26):**

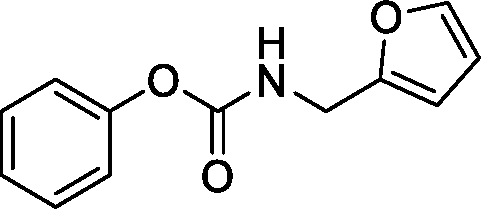

Following the general procedure described for compound **ES_ACR18**, the titeled compound **ES_ACR26** was synthesized strarting from furfuryl amine (280 µL, 3.1 mmol) and obtained as a white solid after flash column chromatography (10-15% ethylacetate in pet. ether).

Yield: 400 mg (59%). R*_f_*: 0.43 (PE/EA 4:1, visualized with UV illumination at 254 nm). ^1^H NMR (500 MHz, CDCl_3_, ppm) δ 7.39 (d, *J* = 1.1 Hz, 1H), 7.36 (t, *J* = 7.9 Hz, 2H), 7.20 (t, *J* = 7.4 Hz, 1H), 7.14 (dd, *J* = 8.6, 1.0 Hz, 2H), 6.37 – 6.34 (m, 1H), 6.29 (dd, *J* = 3.2, 0.6 Hz, 1H), 5.37 (br.s, 1H), 4.48 (d, *J* = 11.9 Hz, 1H), 4.45 (d, *J* = 5.7 Hz, 2H). ^13^C NMR (126 MHz, CDCl_3_, ppm) δ 154.52, 151.19, 151.10, 142.53, 129.42, 125.52, 121.69, 110.63, 107.79, 38.37. ESI-MS *m/z* calcd for C_12_H_12_NO_3_ [M+H]^+^ 218.08; observed 218.1.

***N*-(4-chloro-3-(trifluoromethyl)-phenyl)-4-isopropylcyclohexane-1-carboxamide (ES_ACR27):**

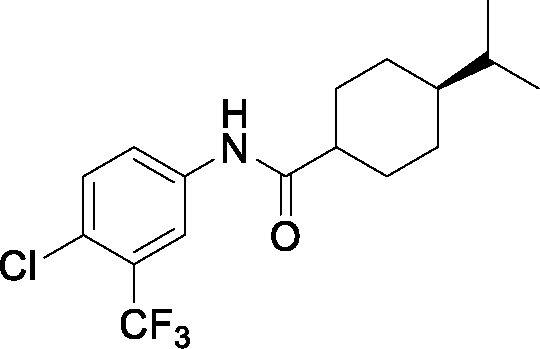

A stirred solution of *trans*-4-*iso*-propylcyclohexane-1-carboxylic acid (330 mg, 2mmol) in dry DMF (3mL) under argon atmosphere at ambient temperature was treated with *N*-methylmorpholine (380 µL, 3.4 mmol), followed by HCTU (1.4 g, 3.4 mmol) and 4-chloro-3-(trifluoromethyl)aniline (0.33 g, 1.7 mmol). The reaction mixture was allowed to stir at 80°C, and the progress of the reaction was followed by TLC analysis. After 6h, the TLC analysis showed a complete reaction. The reaction mixture was cooled to 0°C and quenched with 1M HCL (50 mL). The resulting mixture was extracted with ethylactetate, and the combined organic layers was washed with brine, dried over anhydrous NaSO_4_, filtered and concentered. The obtained residue was purified by flash column chromatography over silica gel using ethylacetate and per. ether as eluents (10-20% ethylacetate in pet. ether) to yield compound **ES_ACR27** as a white solid.

Yield: 475 mg (81%). R*_f_*: 0.44 (PE/EA 4:1, visualized with UV illumination at 254 nm). ^1^H NMR (500 MHz, CDCl_3_, ppm) δ 7.85 (d, *J* = 2.4 Hz, 1H), 7.74 (dd, *J* = 8.7, 2.4 Hz, 1H), 7.43 (s, 1H), 7.41 (s, 1H), 2.17 (tt, *J* = 12.1, 3.5 Hz, 1H), 2.00 (dd, *J* = 14.5, 2.2 Hz, 2H), 1.84 (dd, *J* = 12.9, 2.5 Hz, 2H), 1.54 (qd, *J* = 12.7, 2.9 Hz, 2H), 1.43 (td, *J* = 13.4, 6.7 Hz, 1H), 1.14 – 1.06 (m, 1H), 1.01 (ddd, *J* = 15.1, 9.7, 3.2 Hz, 2H), 0.87 (d, *J* = 6.8 Hz, 6H). ^13^C NMR (126 MHz, CDCl_3_, ppm) δ 174.94, 137.04, 132.07, 128.97, 126.75, 123.81, 121.59, 118.80, 46.84, 43.28, 32.88, 29.85, 29.01, 19.85. ESI-MS *m/z* calcd for C_17_H_22_NOF_3_Cl [M+H]^+^ 348.13; observed 348.1.

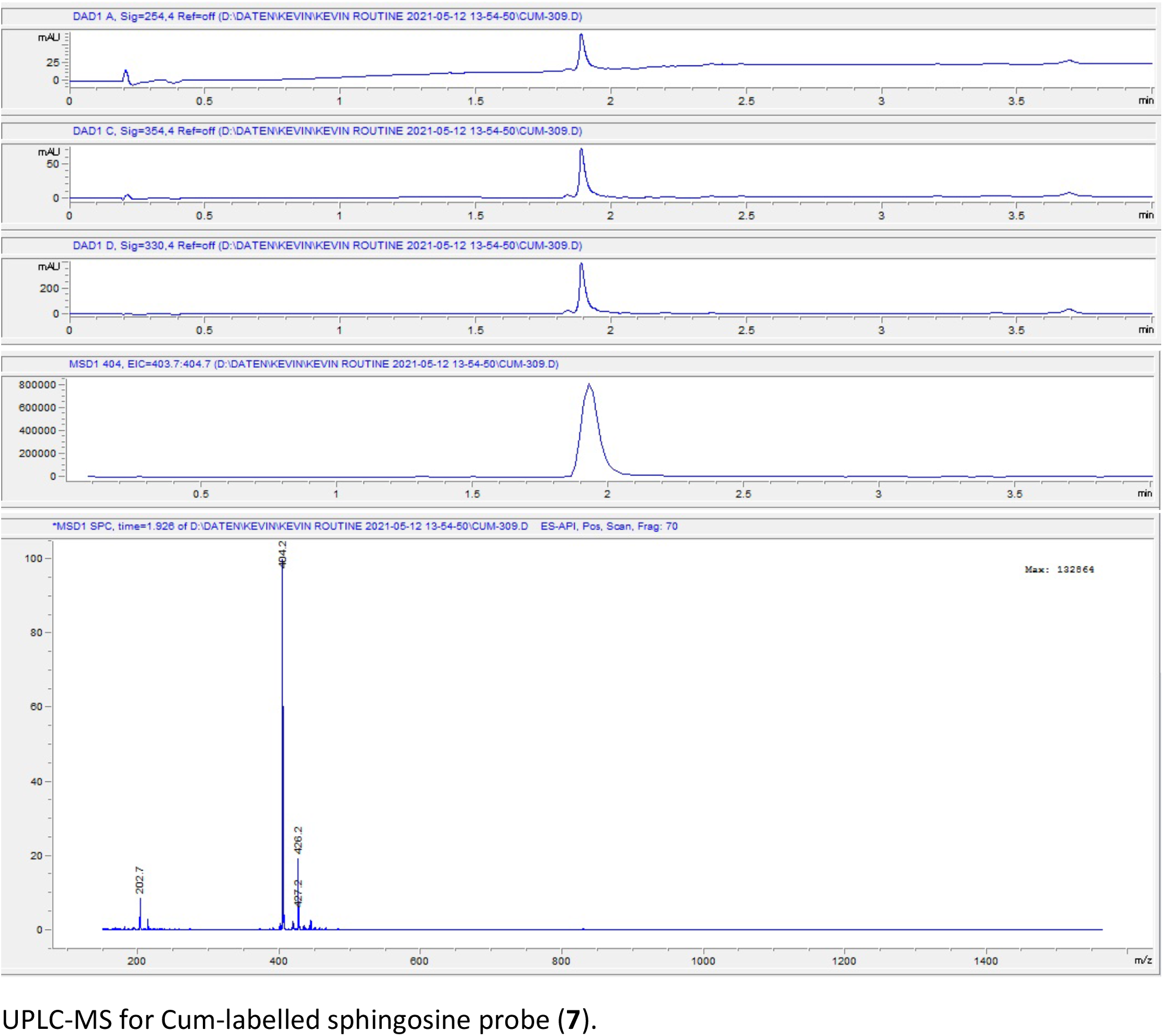

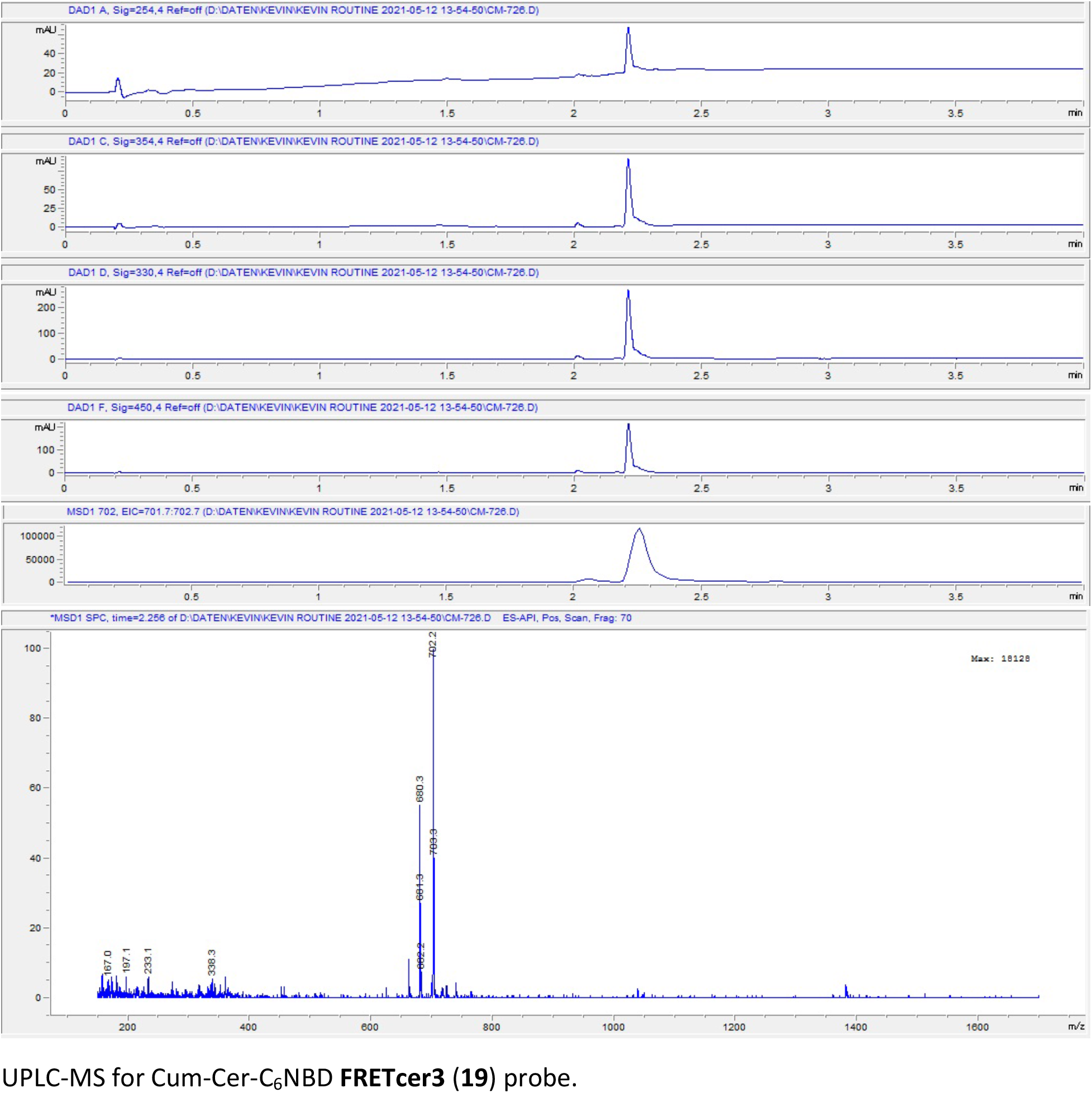

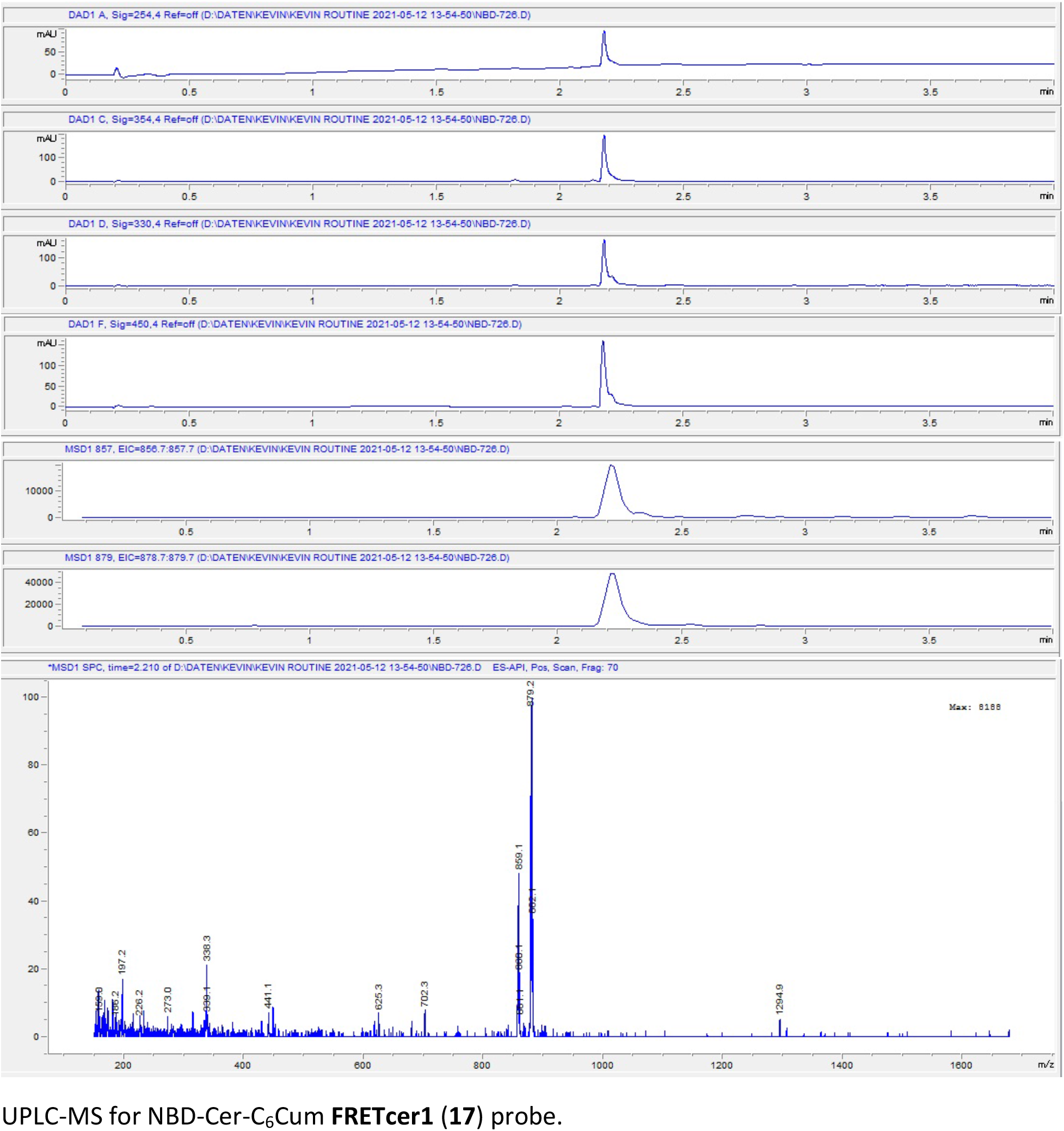

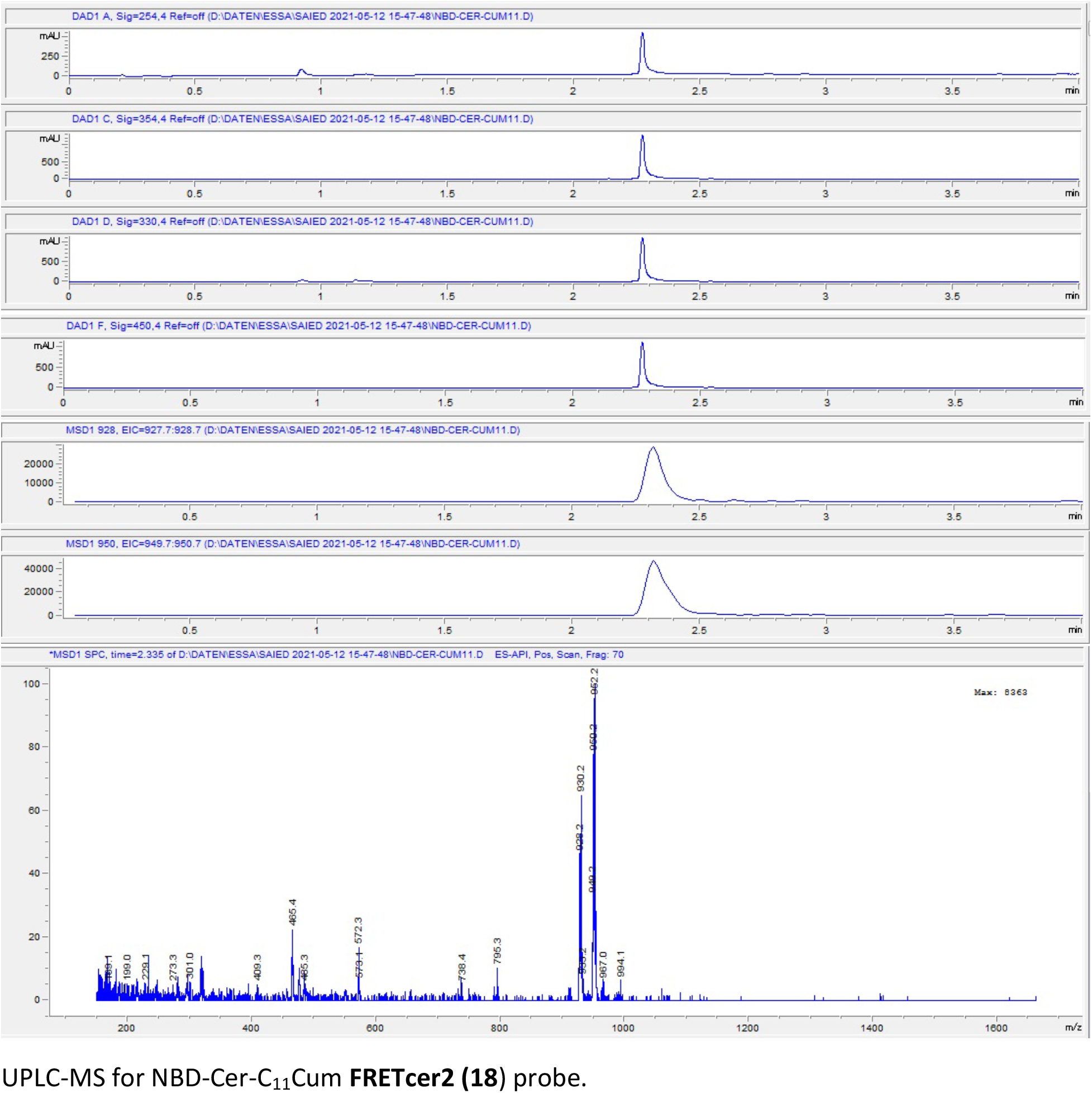

### III. NMR Spectra of the synthesized compounds

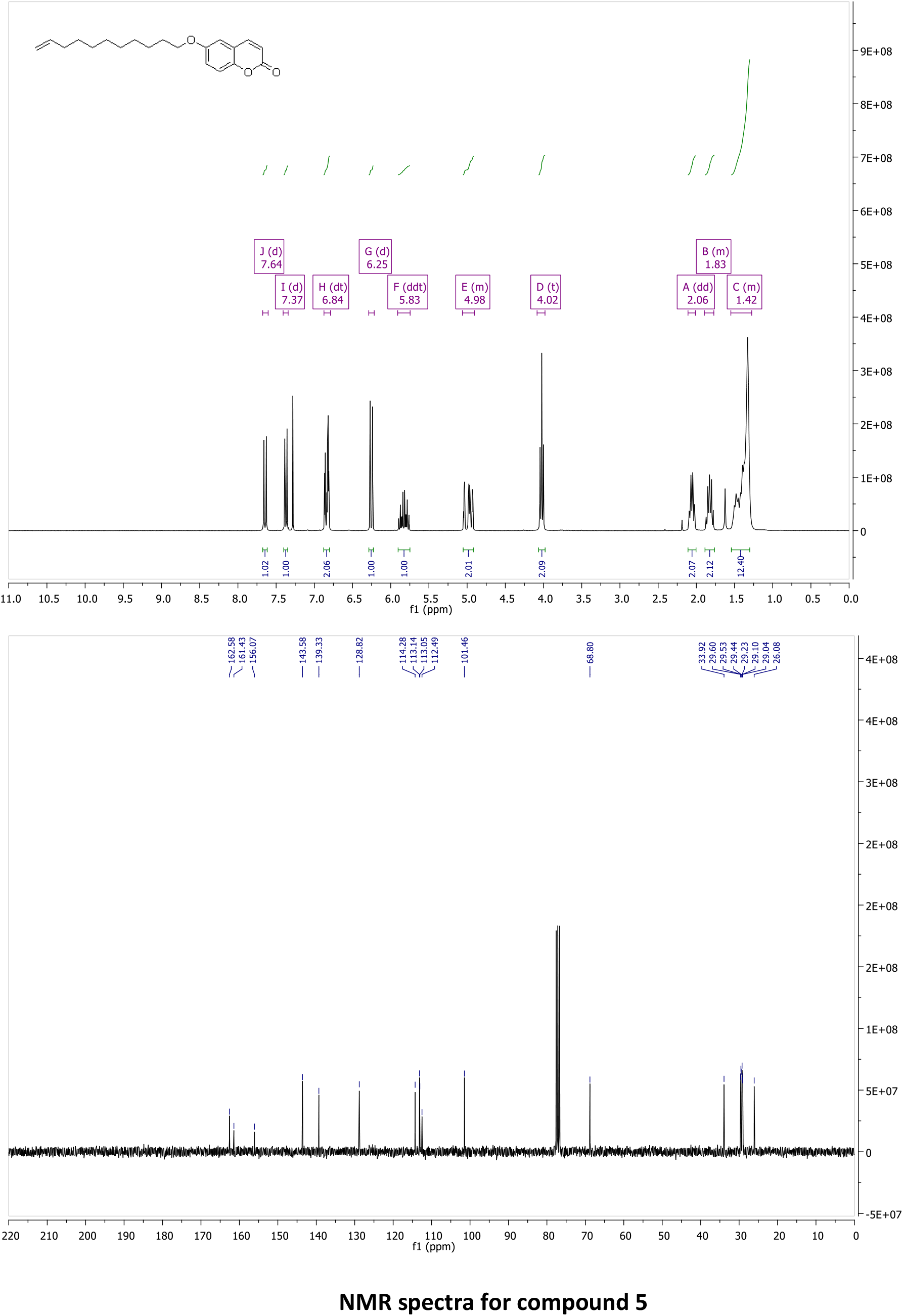
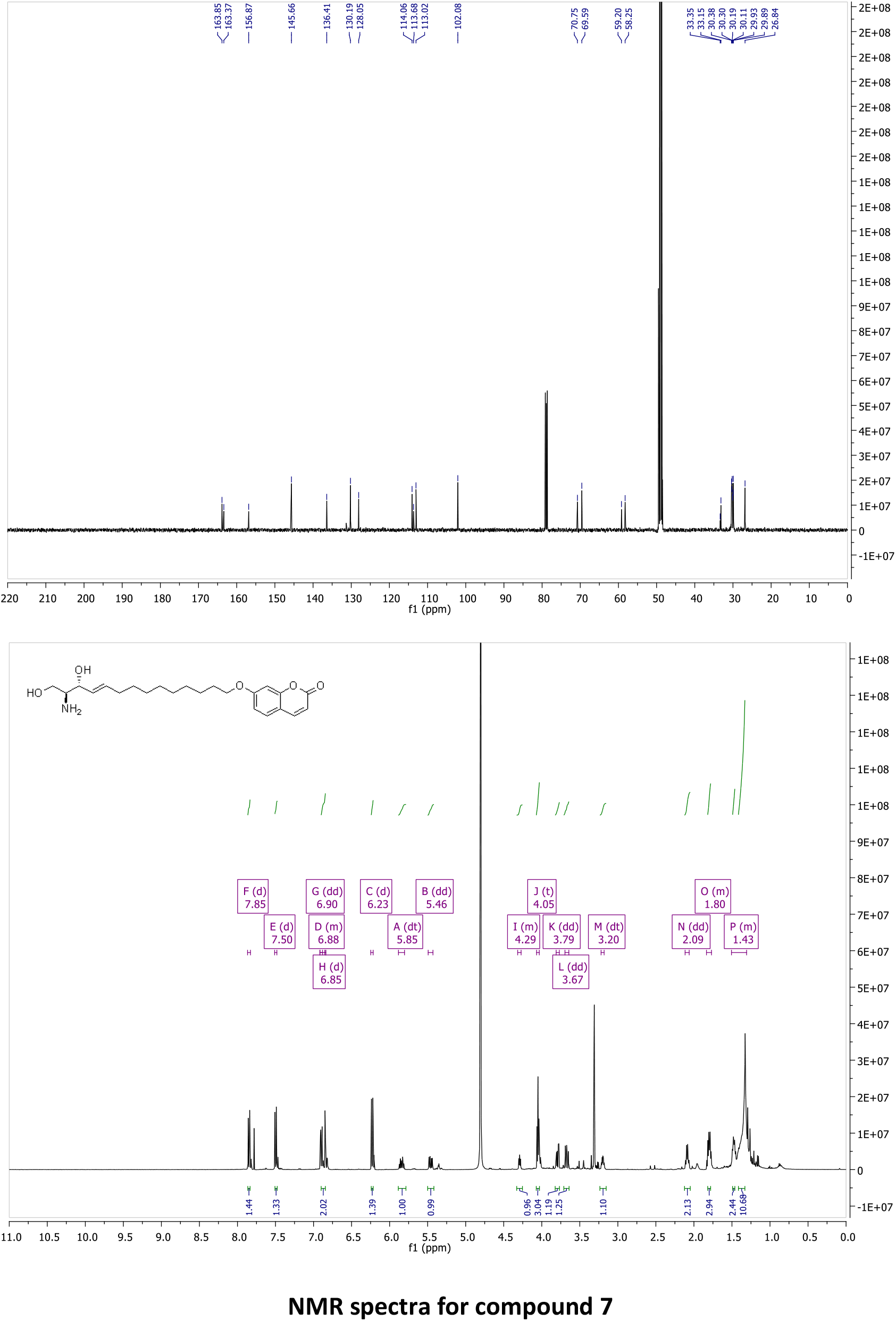
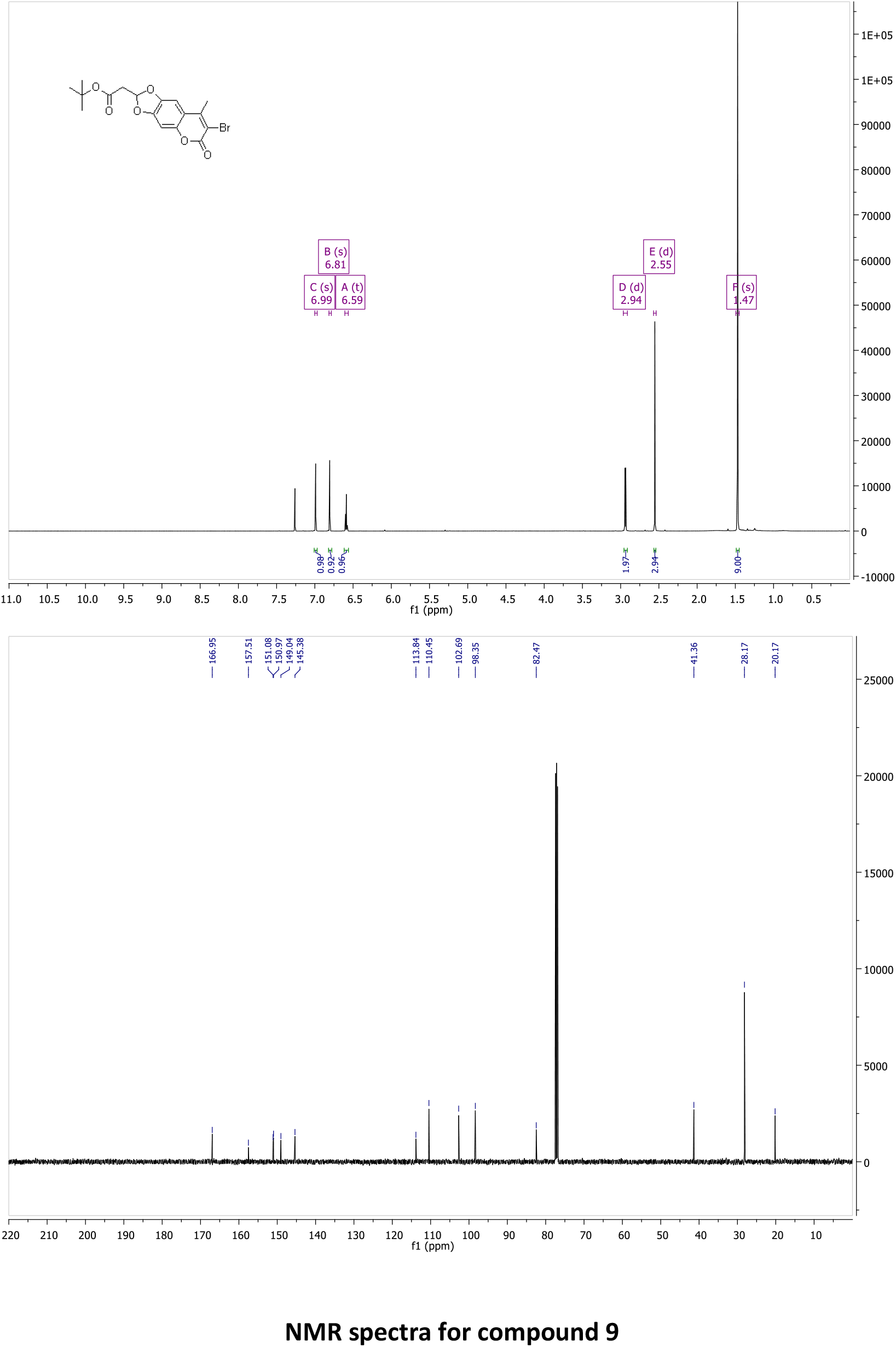
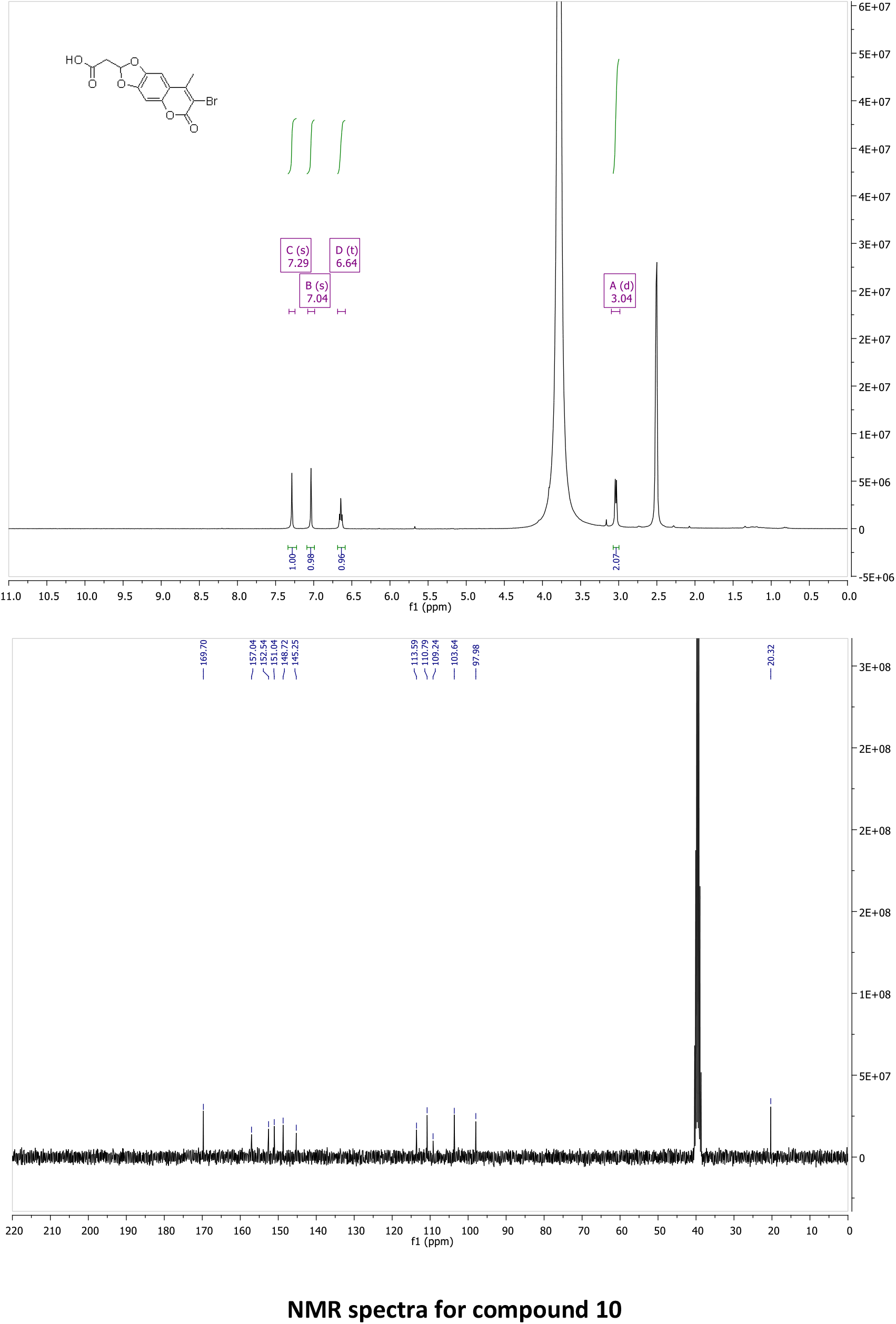
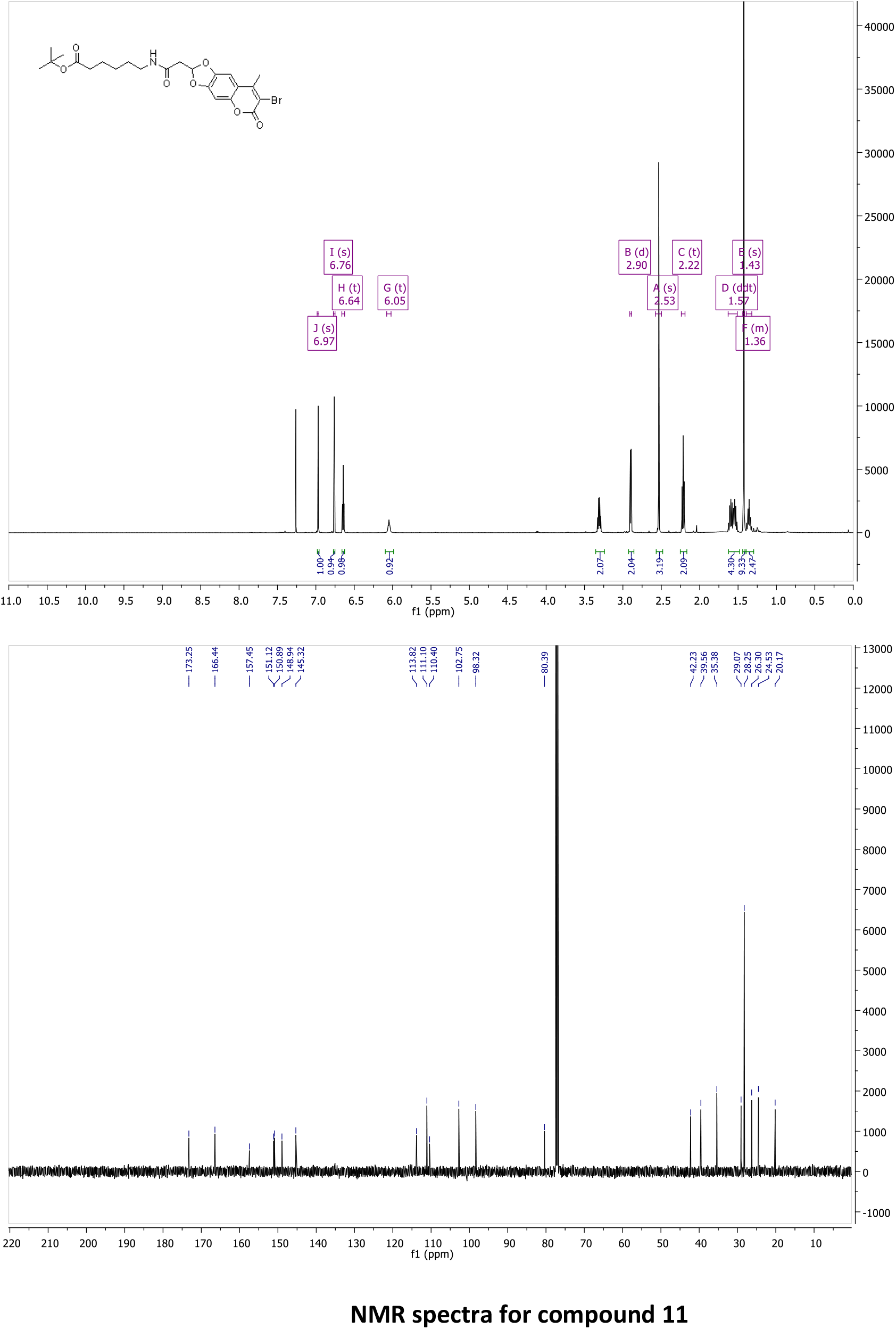
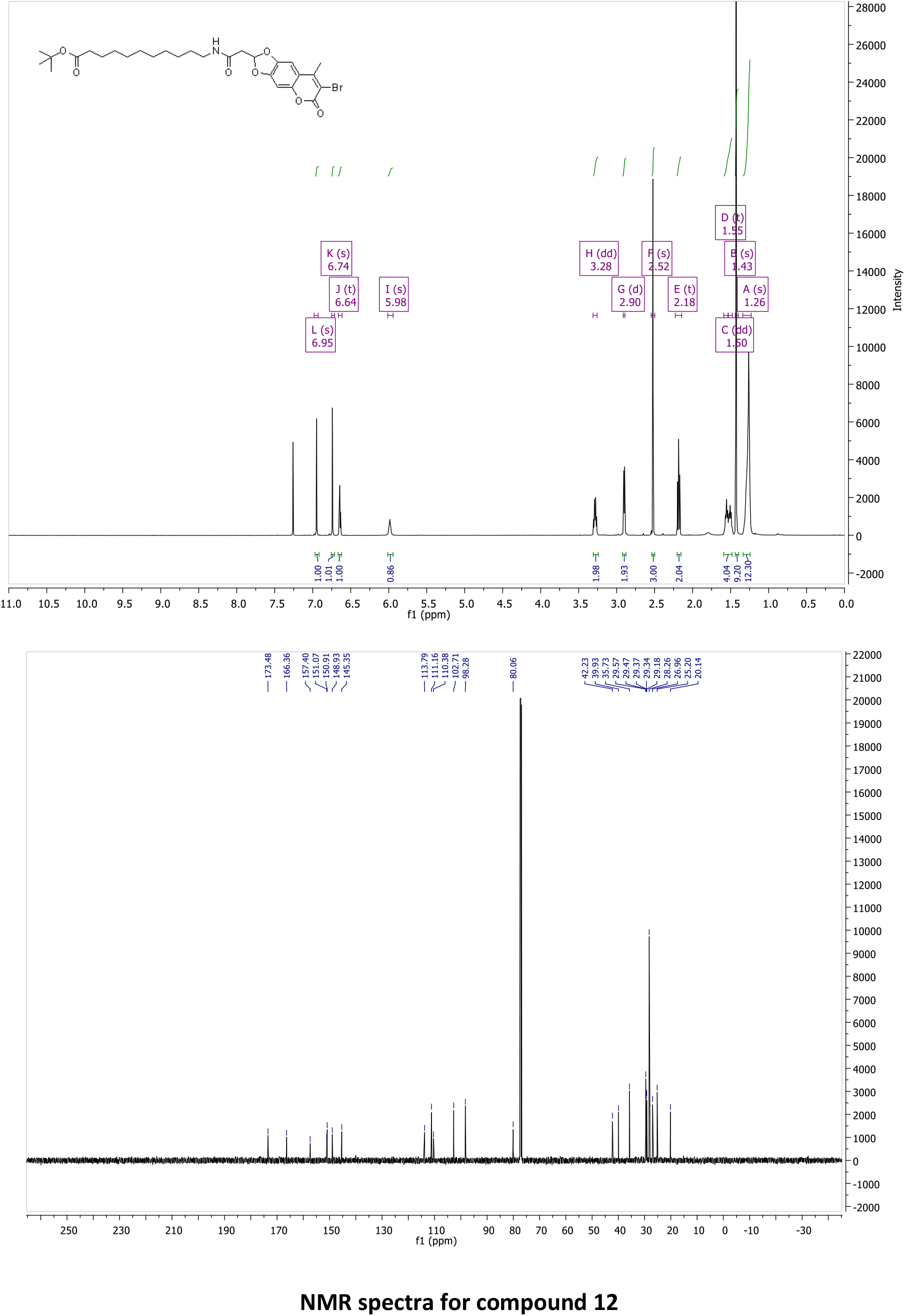
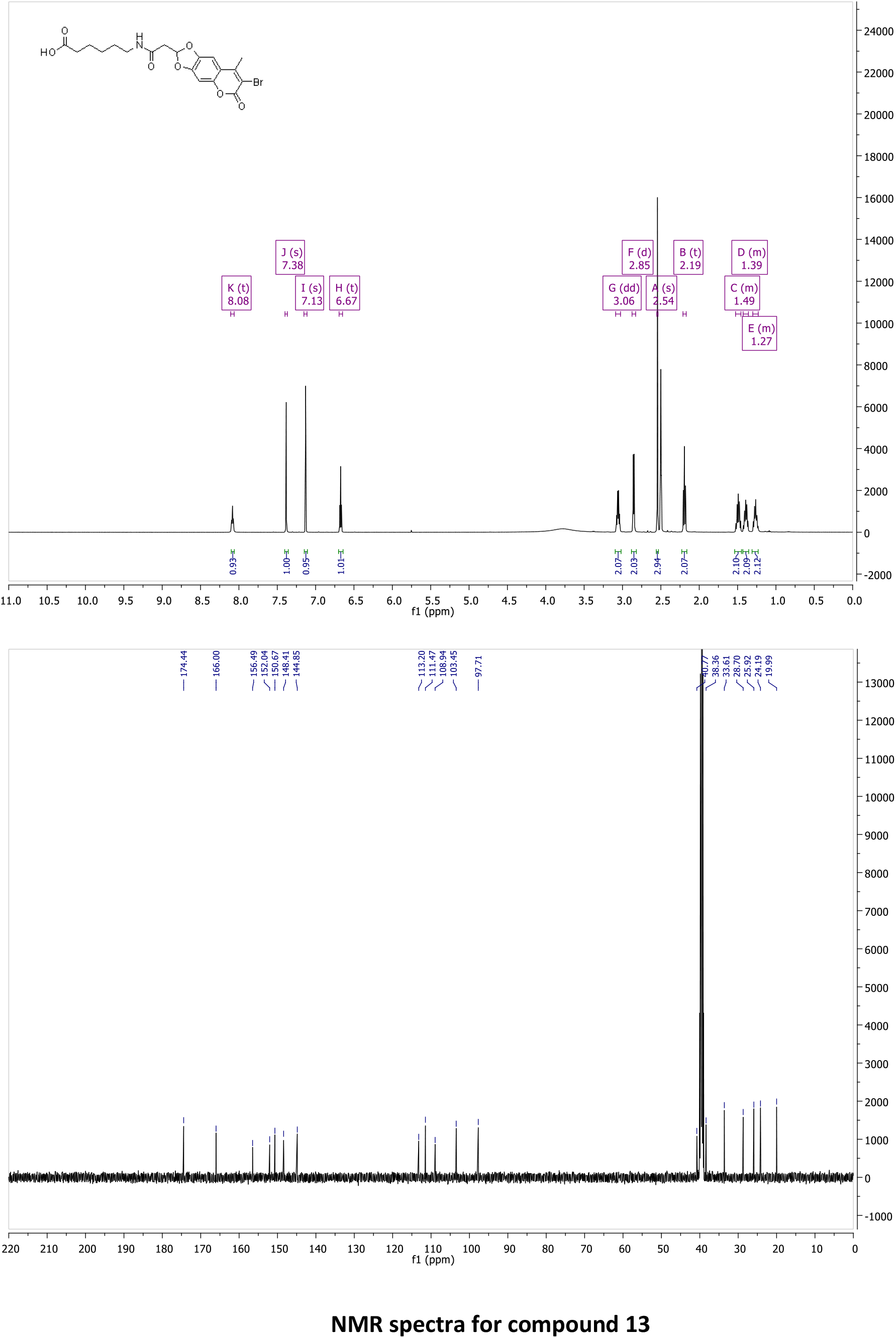
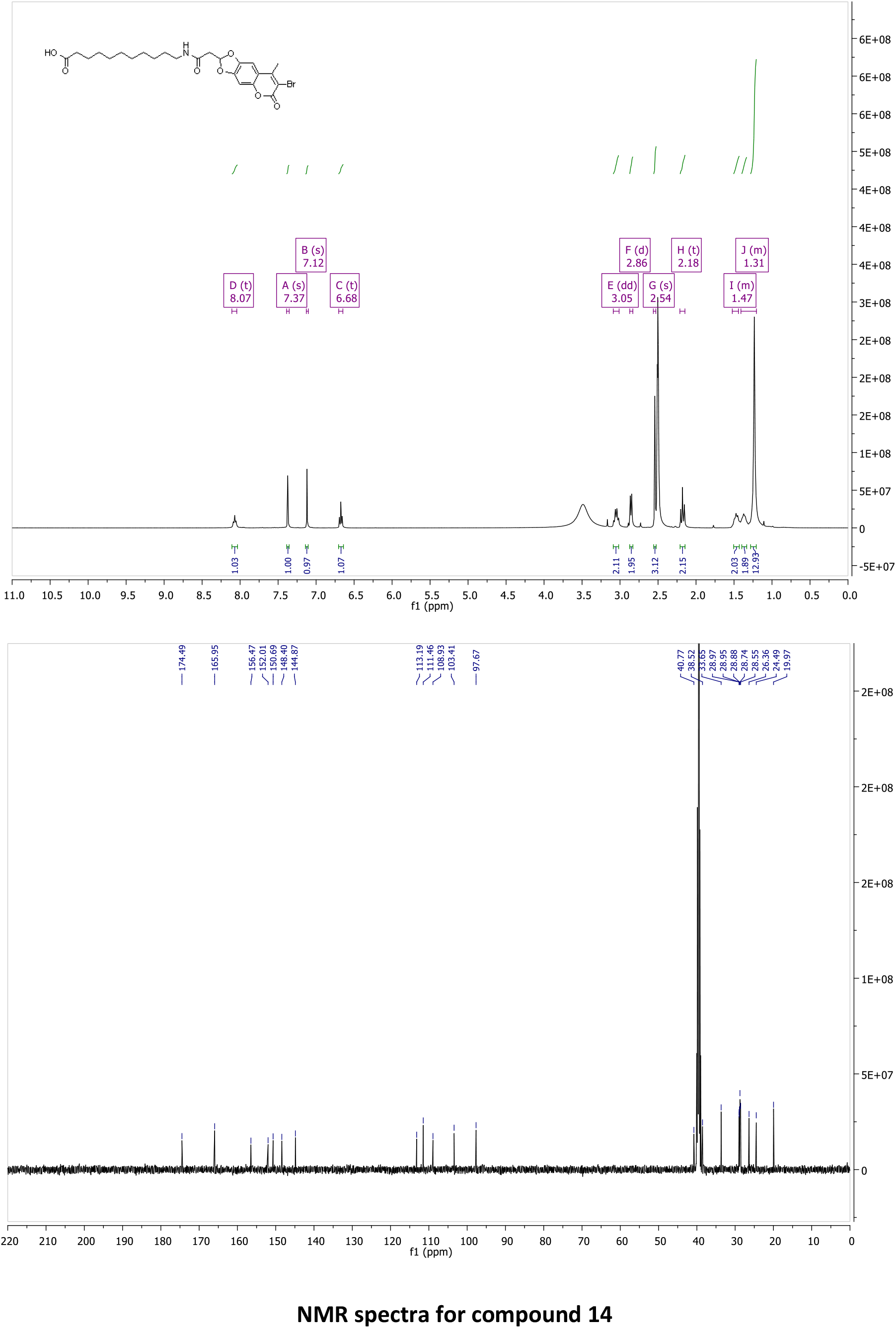
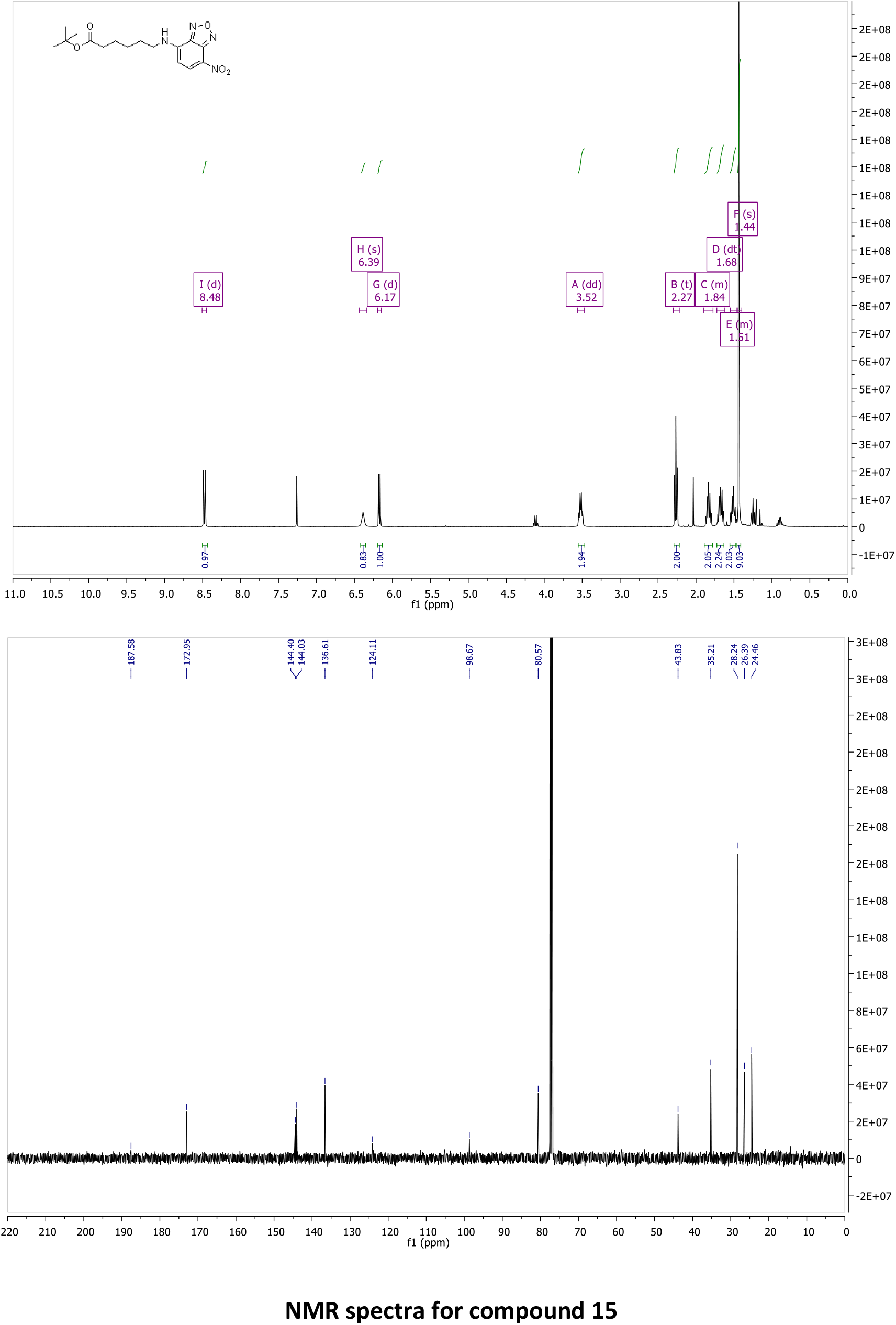
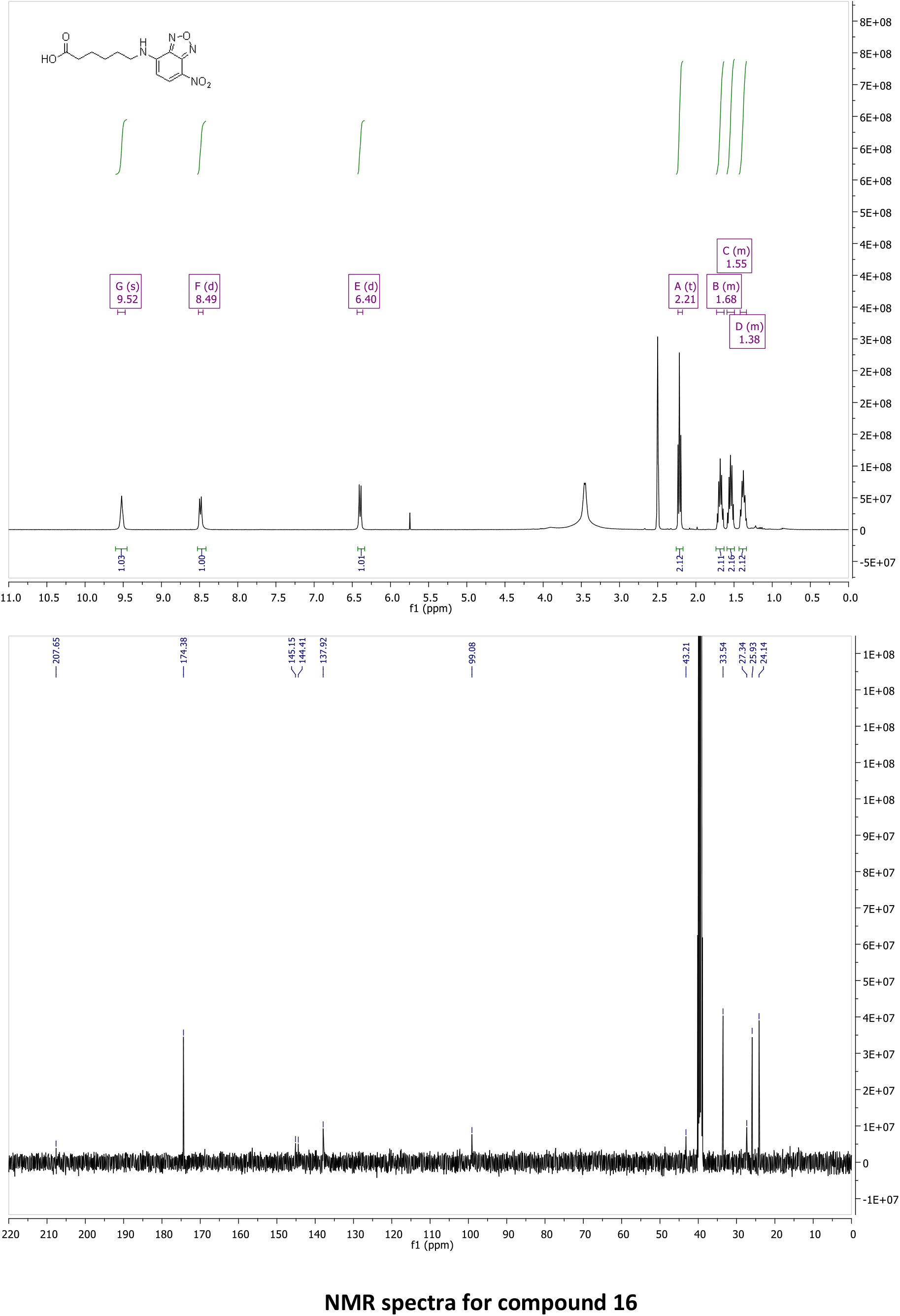
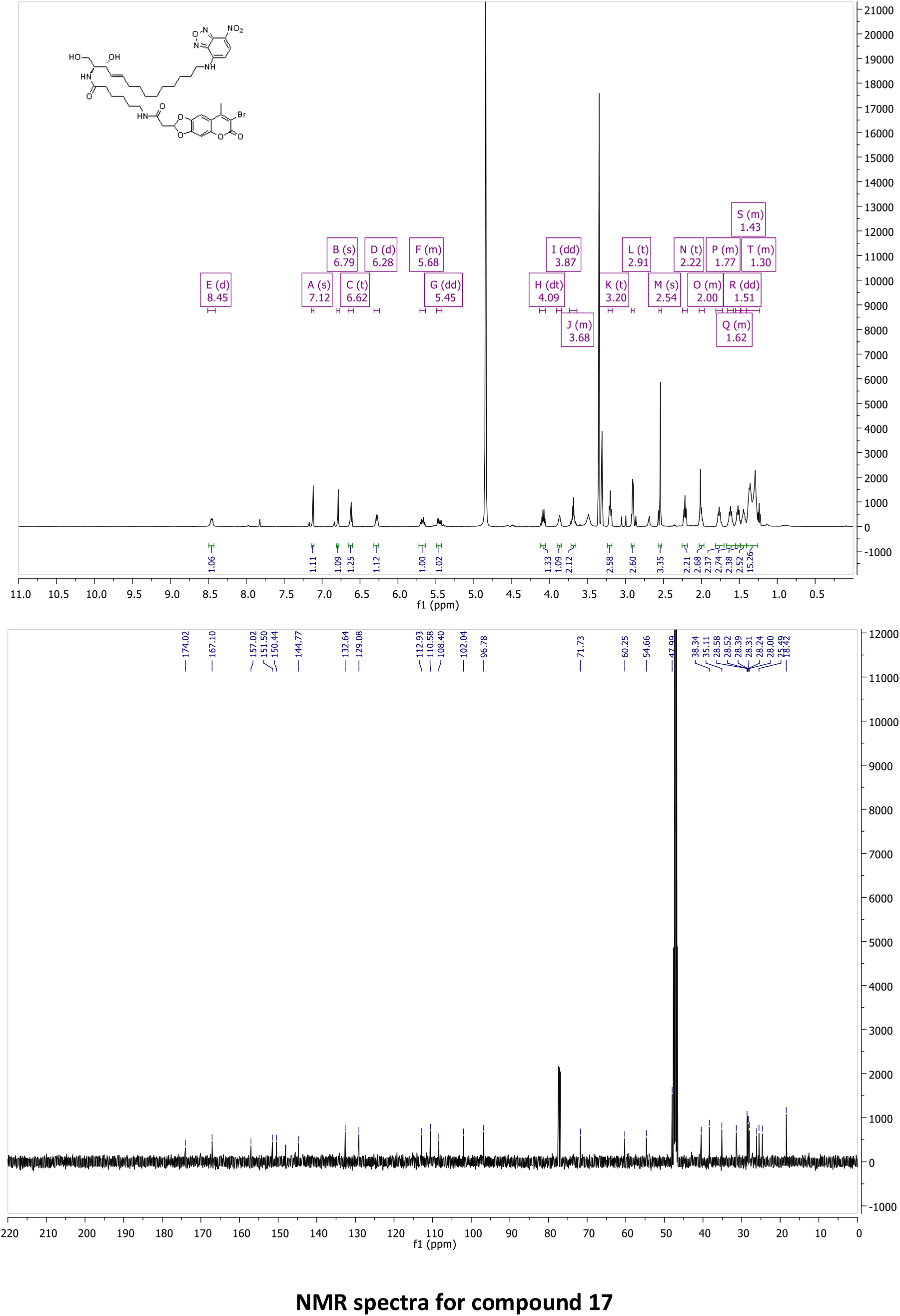
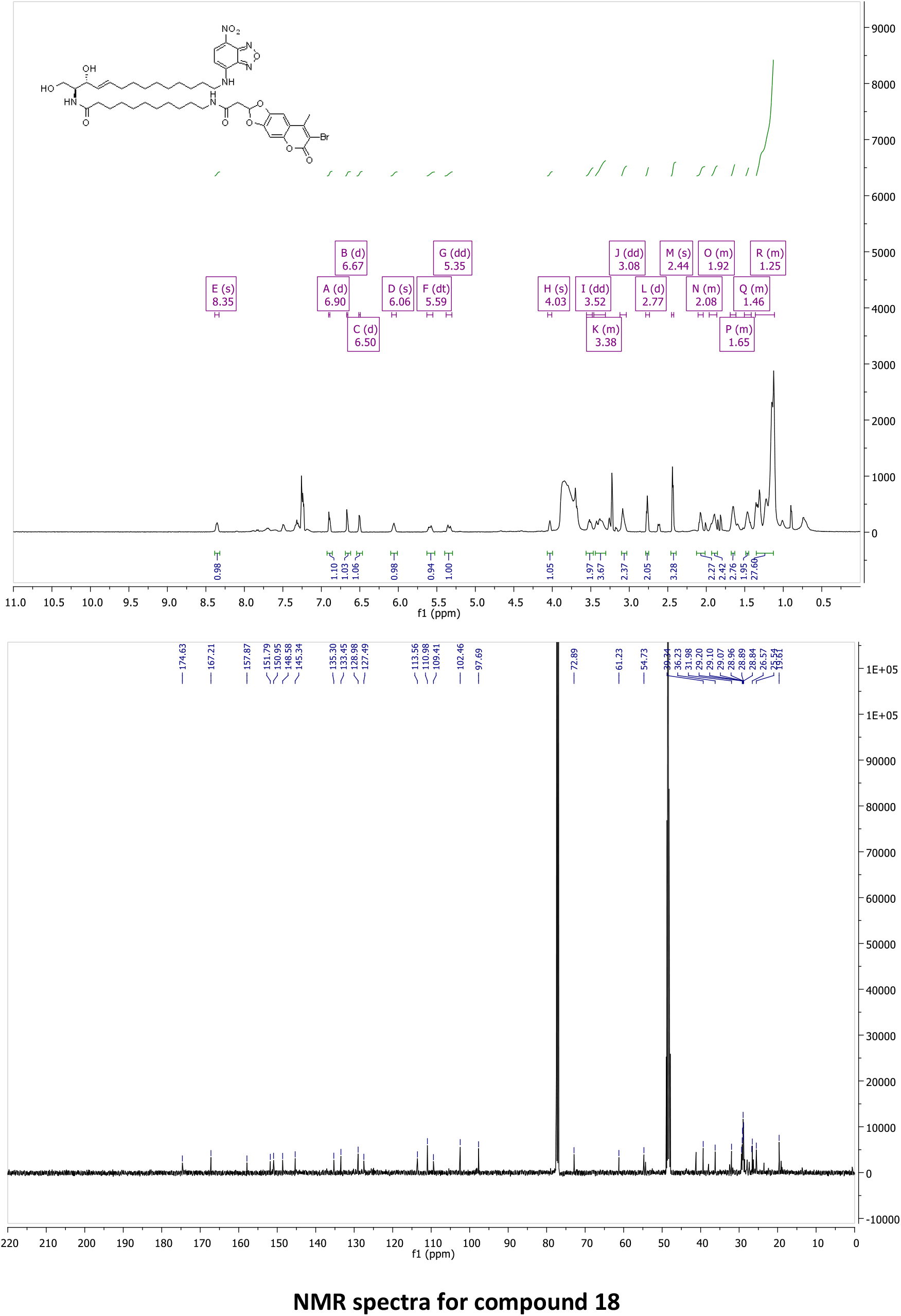
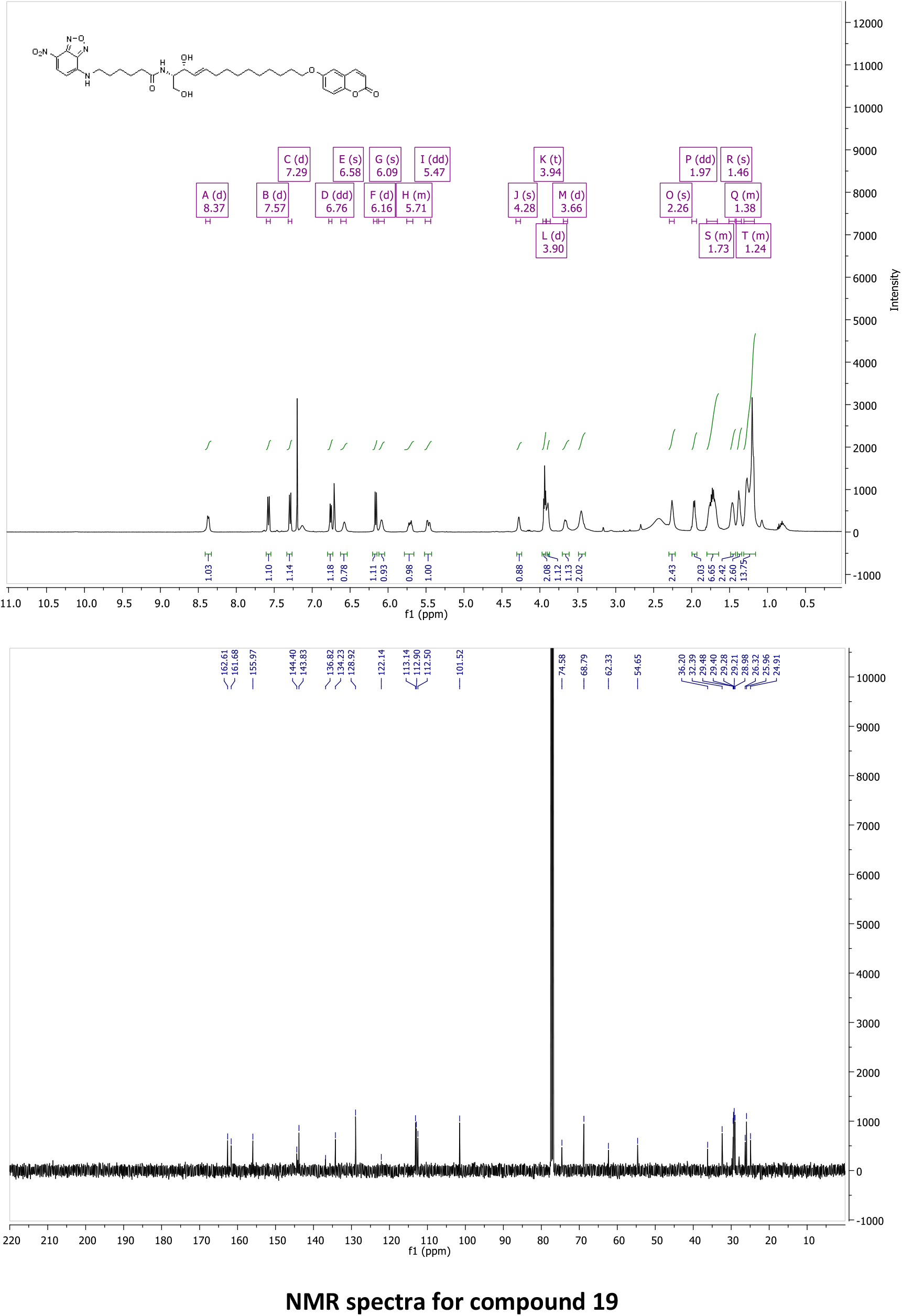
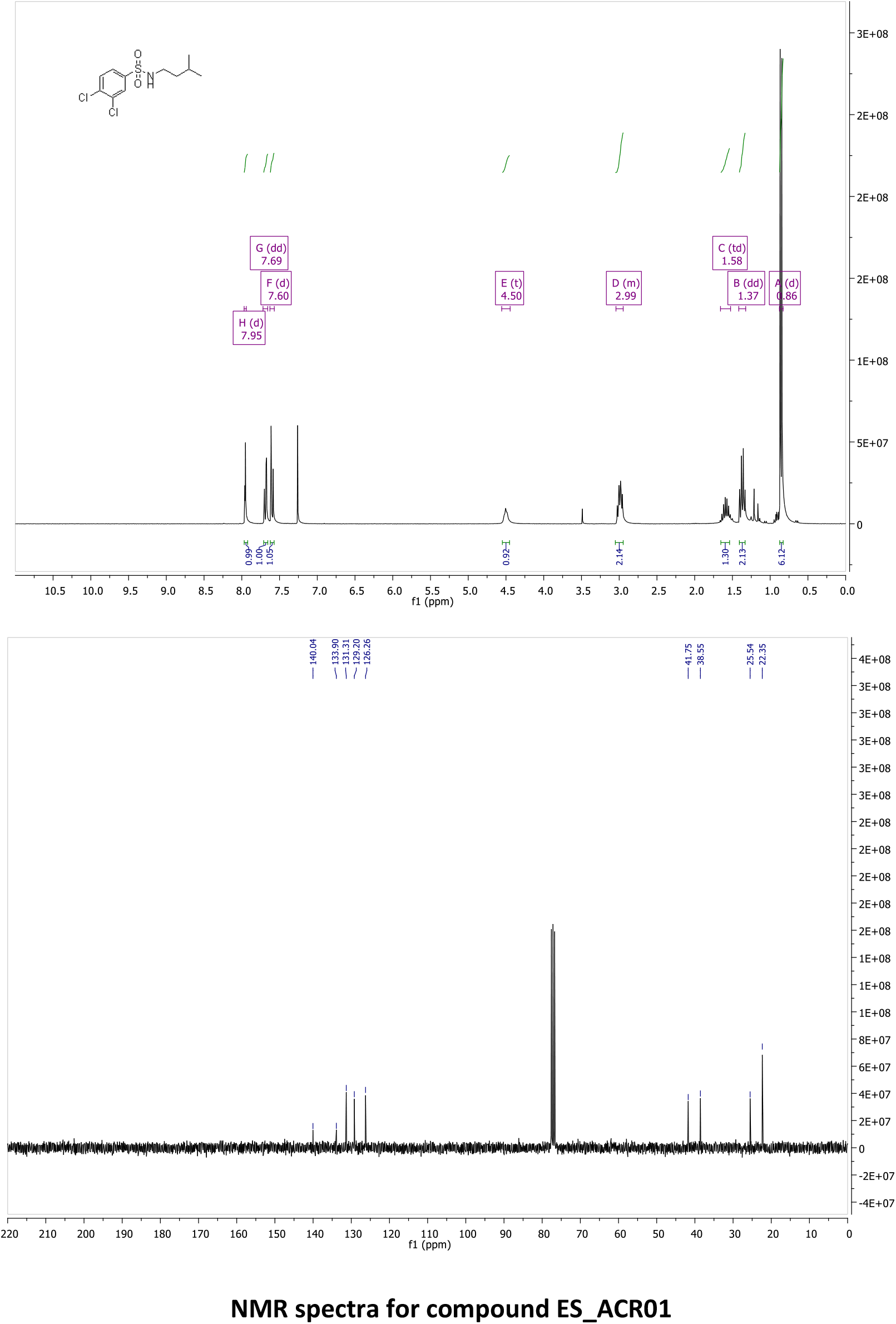
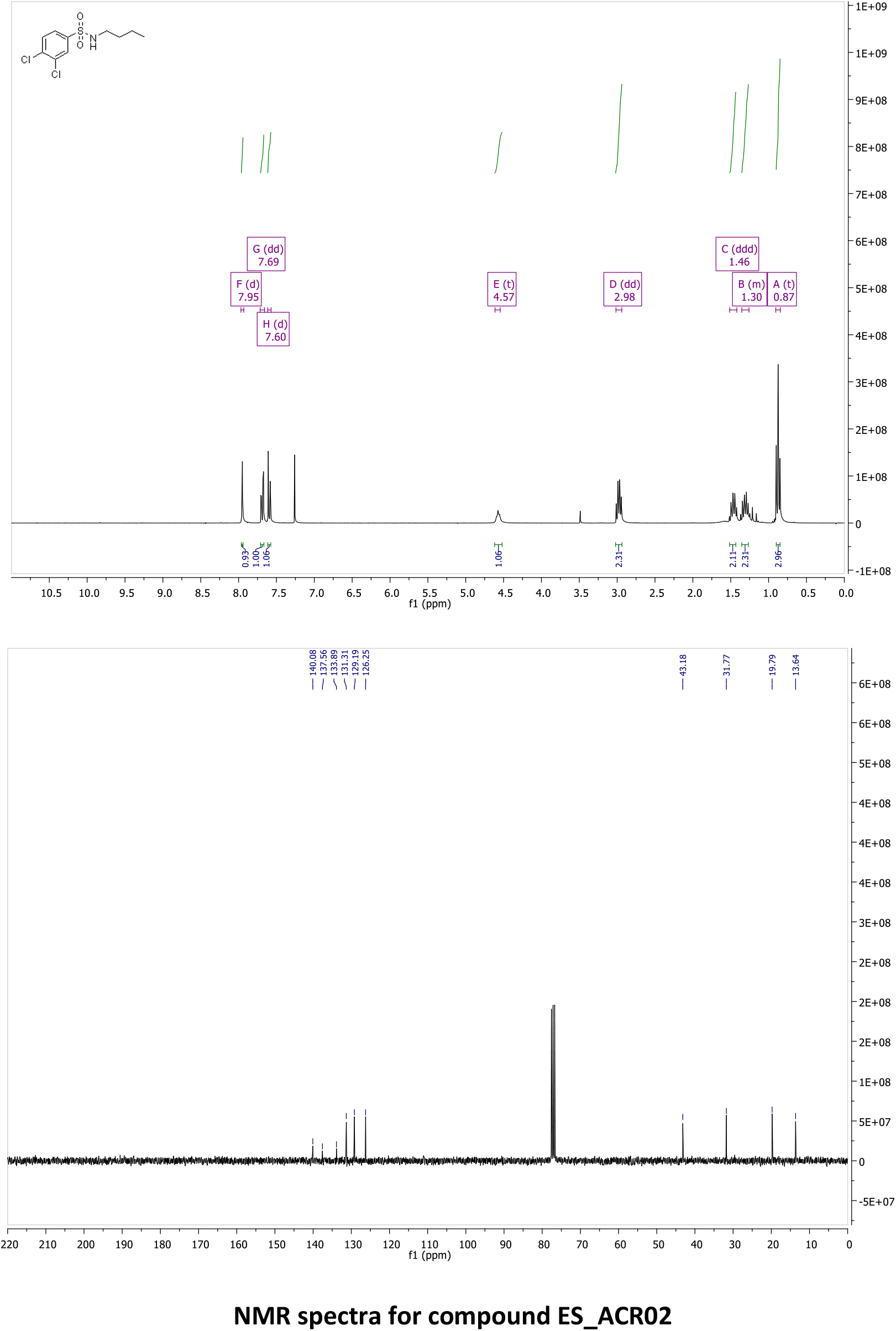
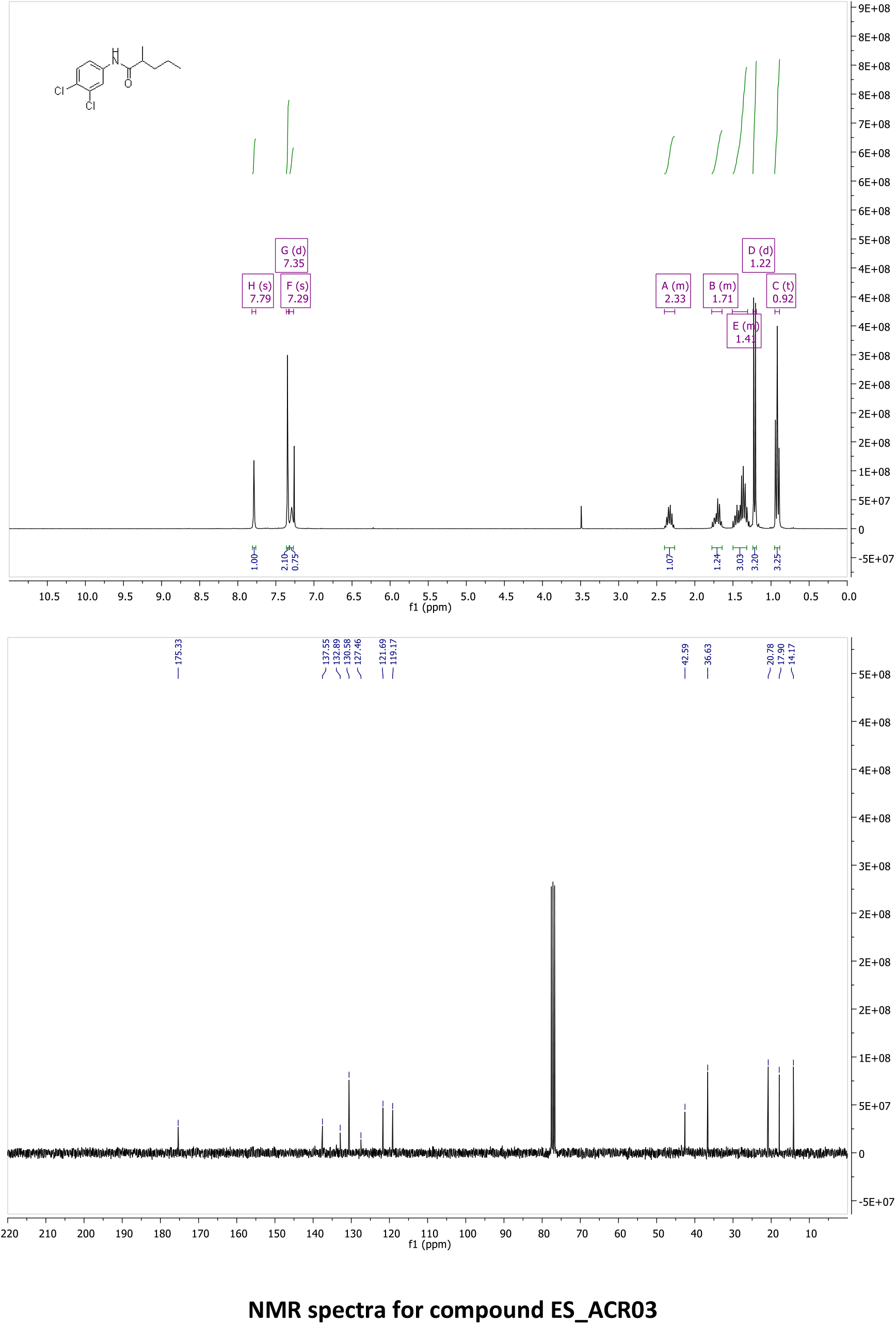
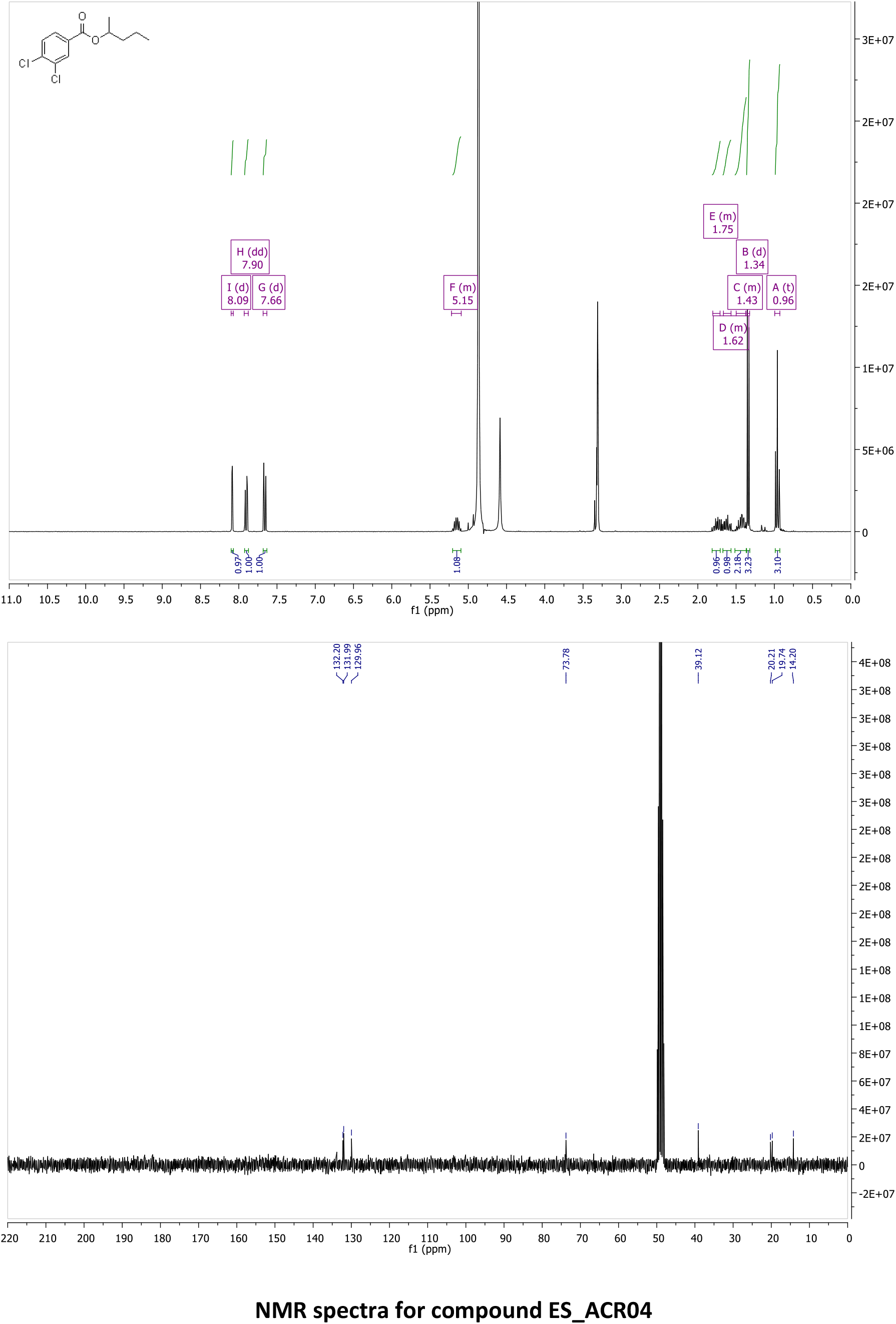
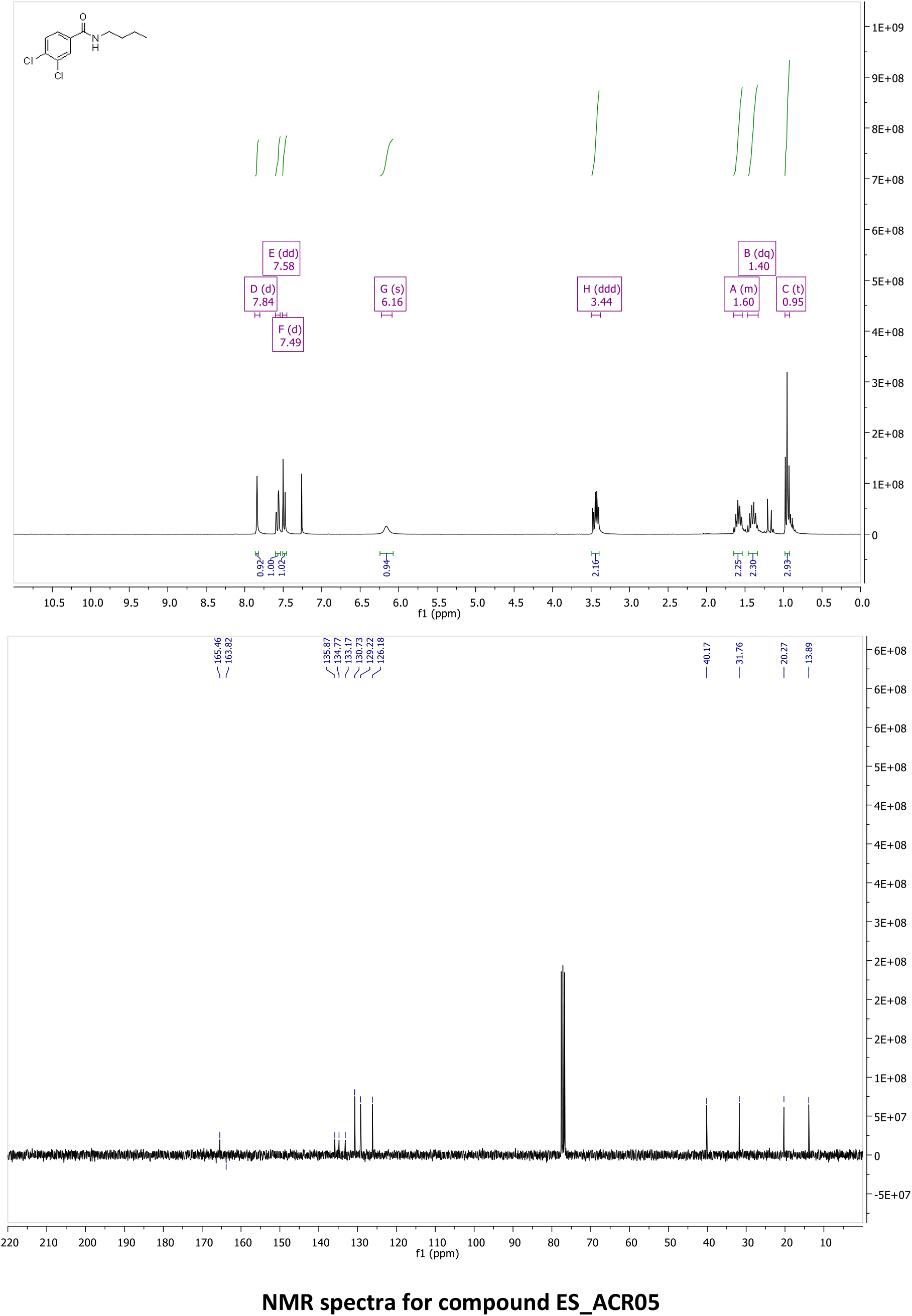
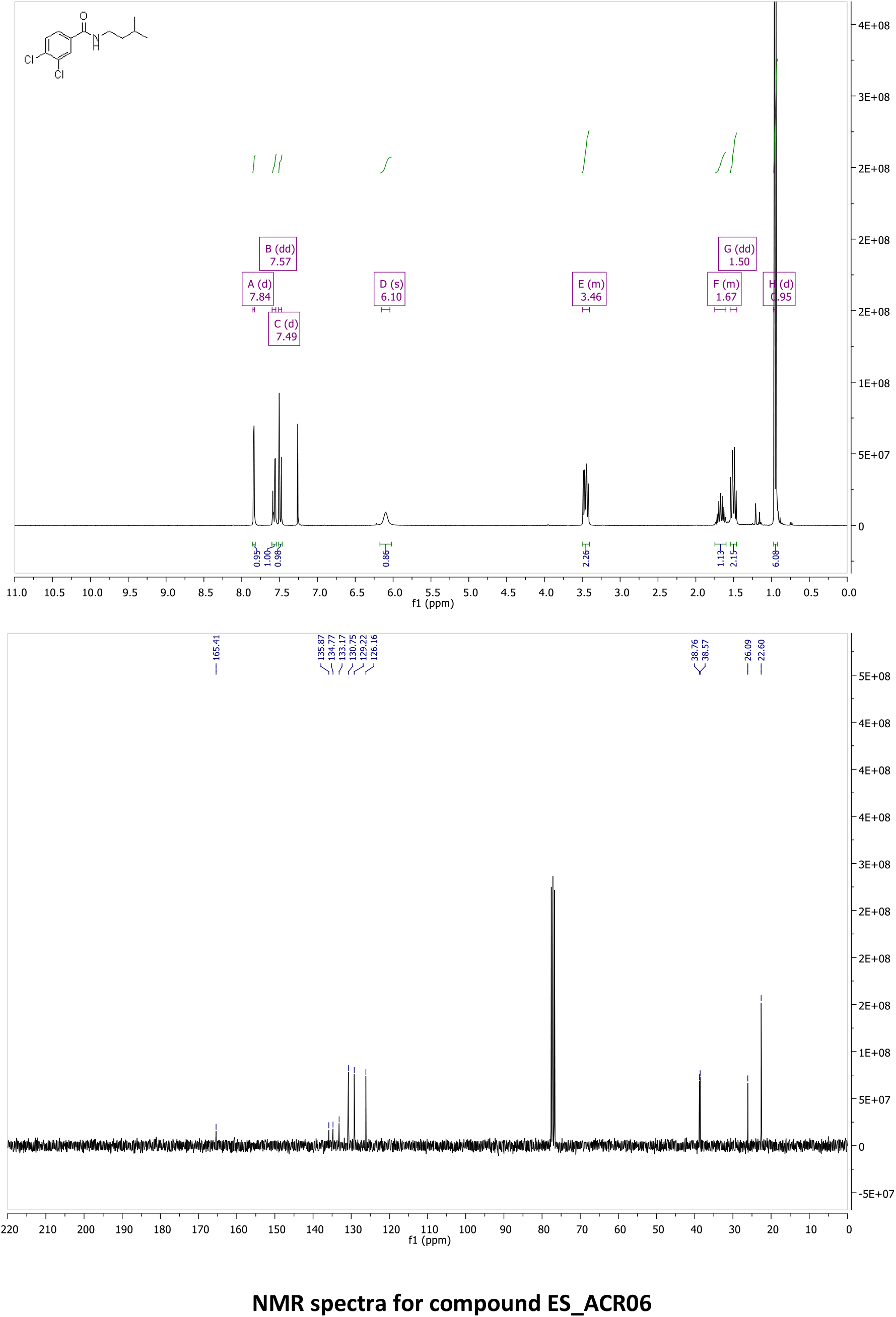
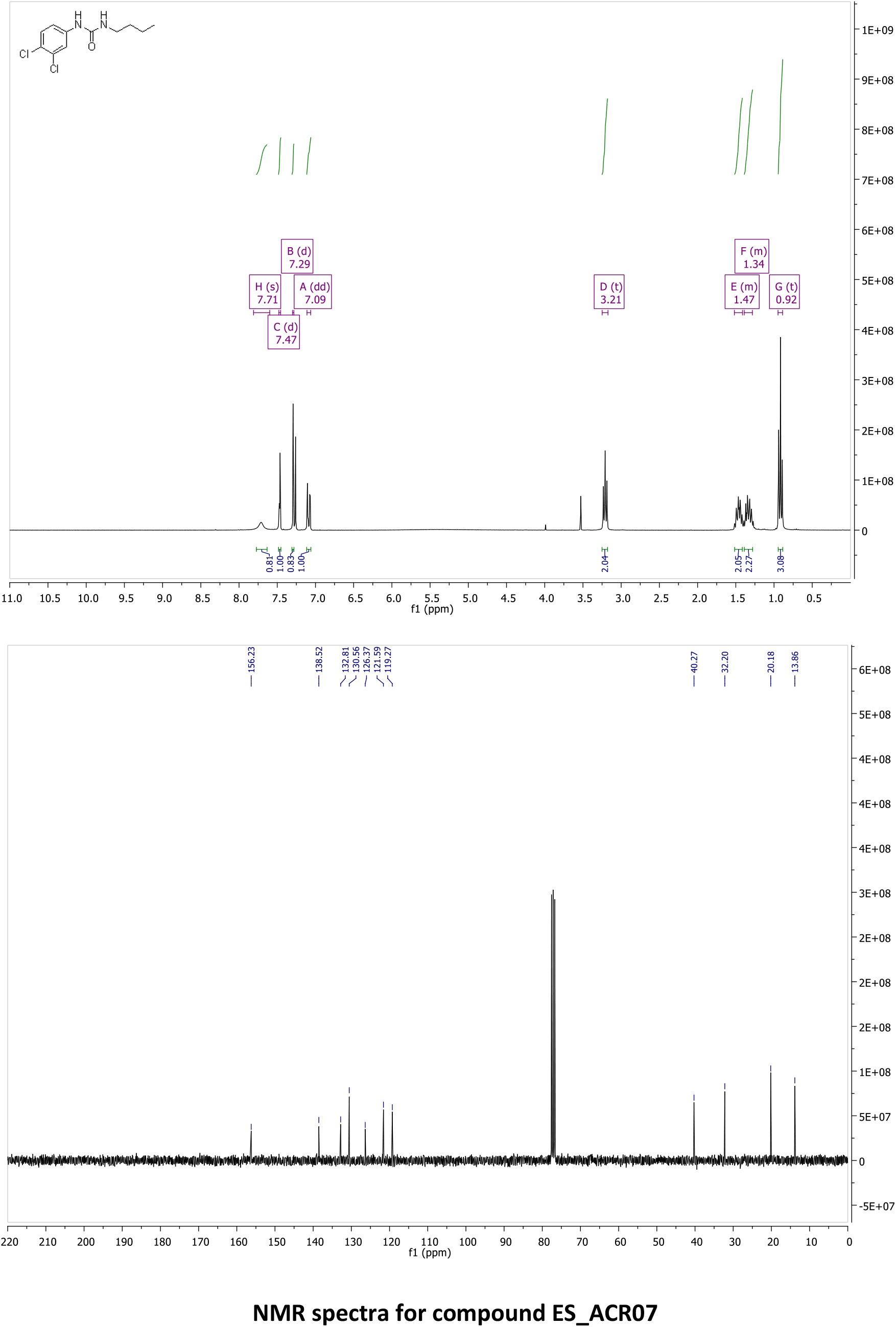
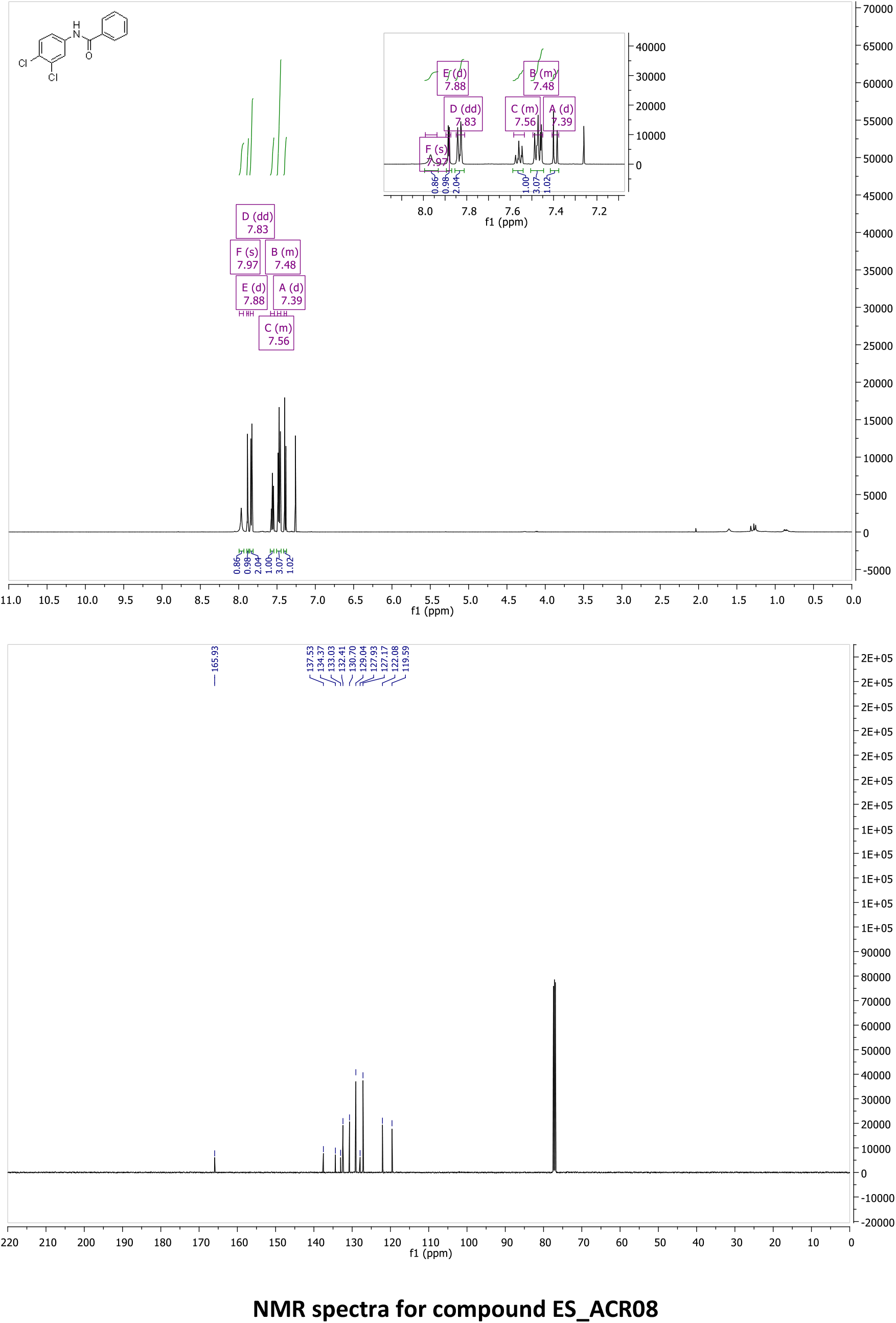
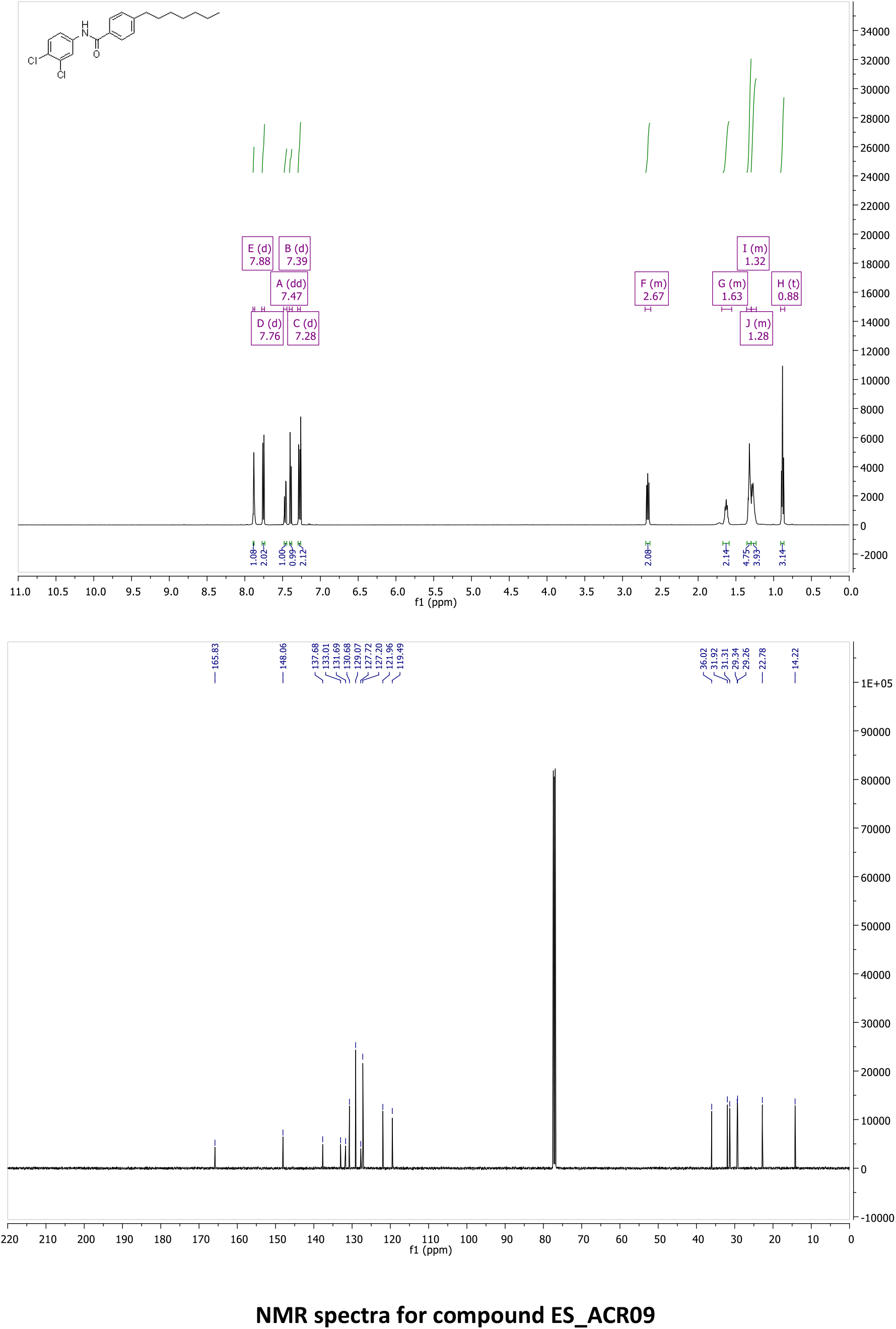
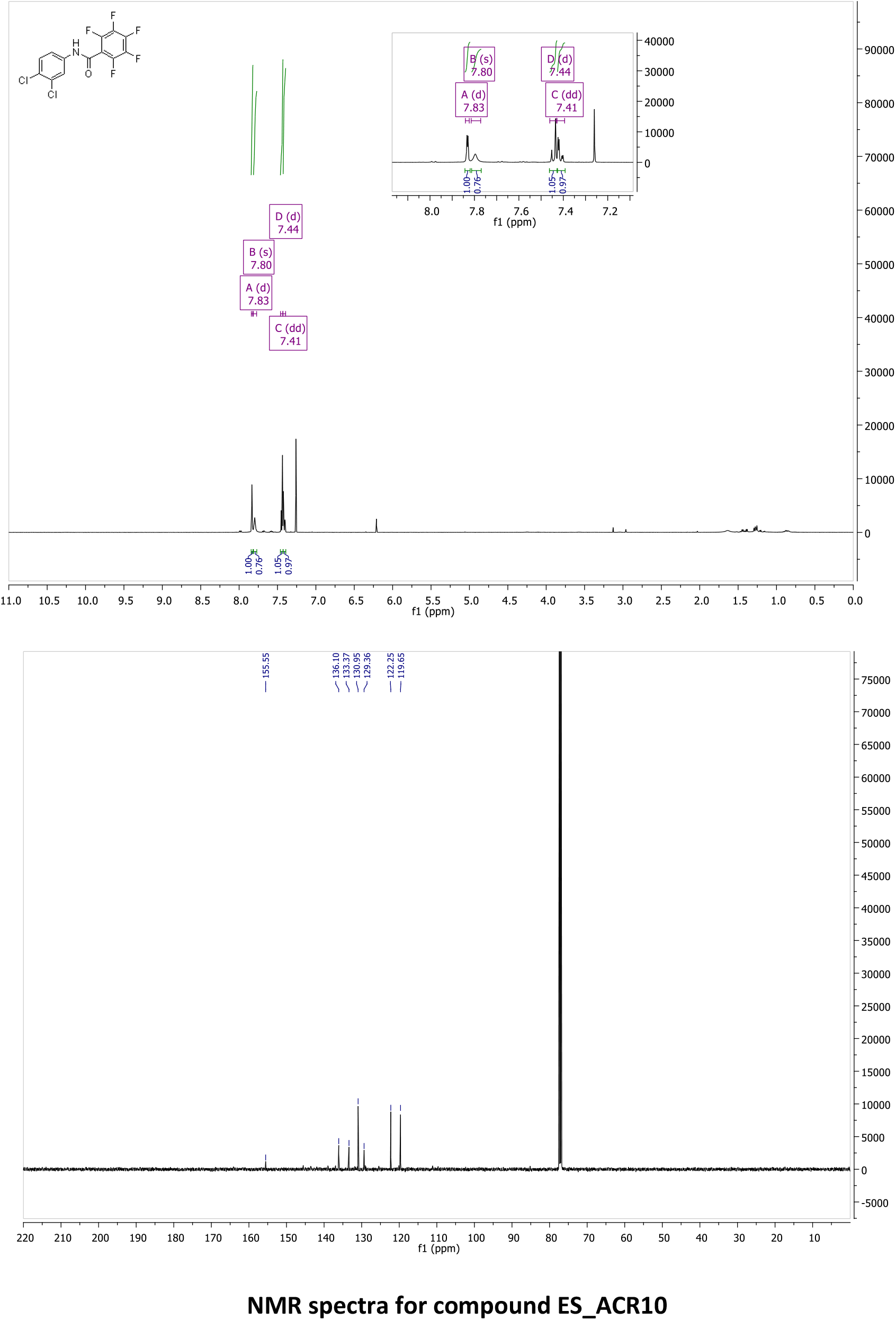
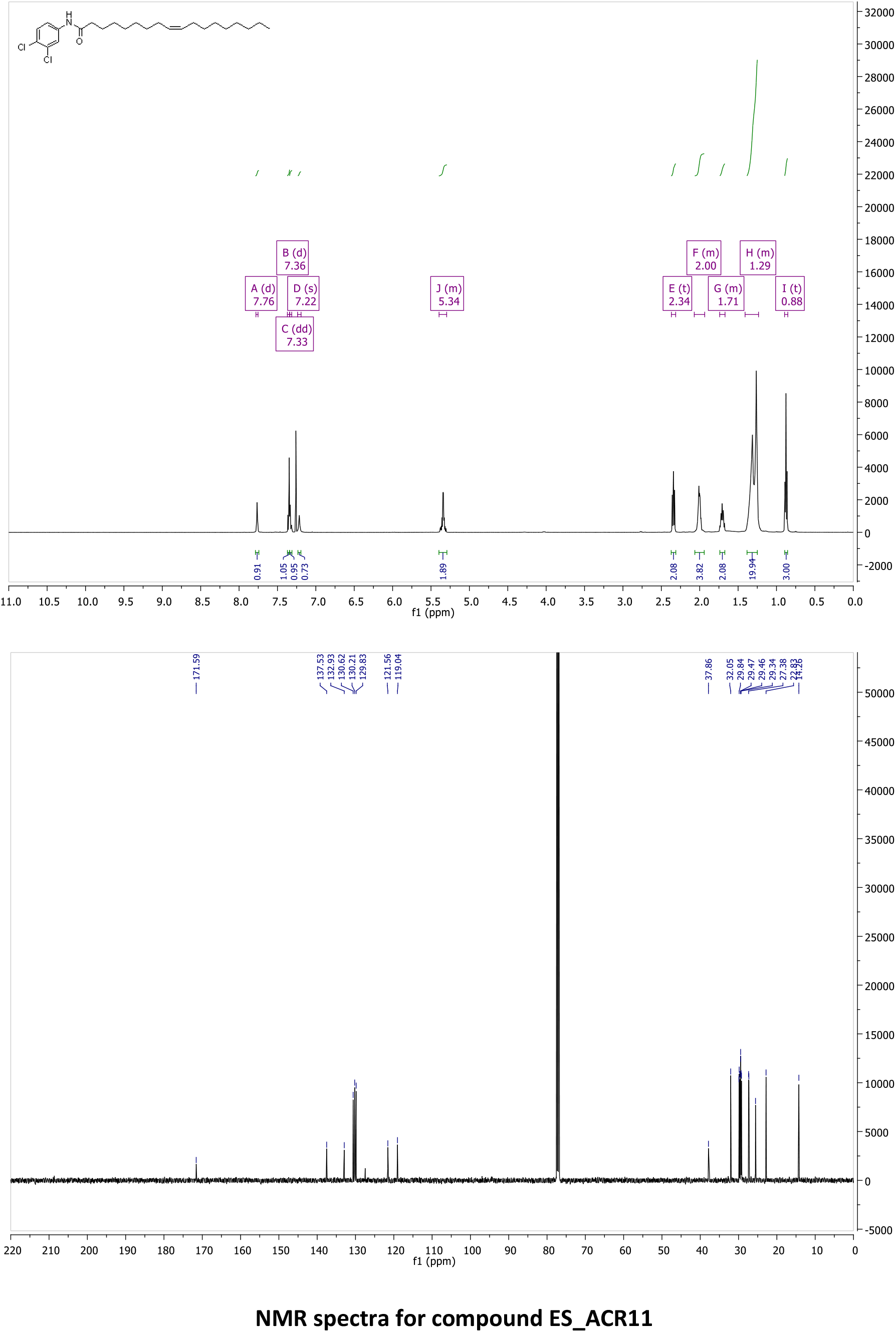
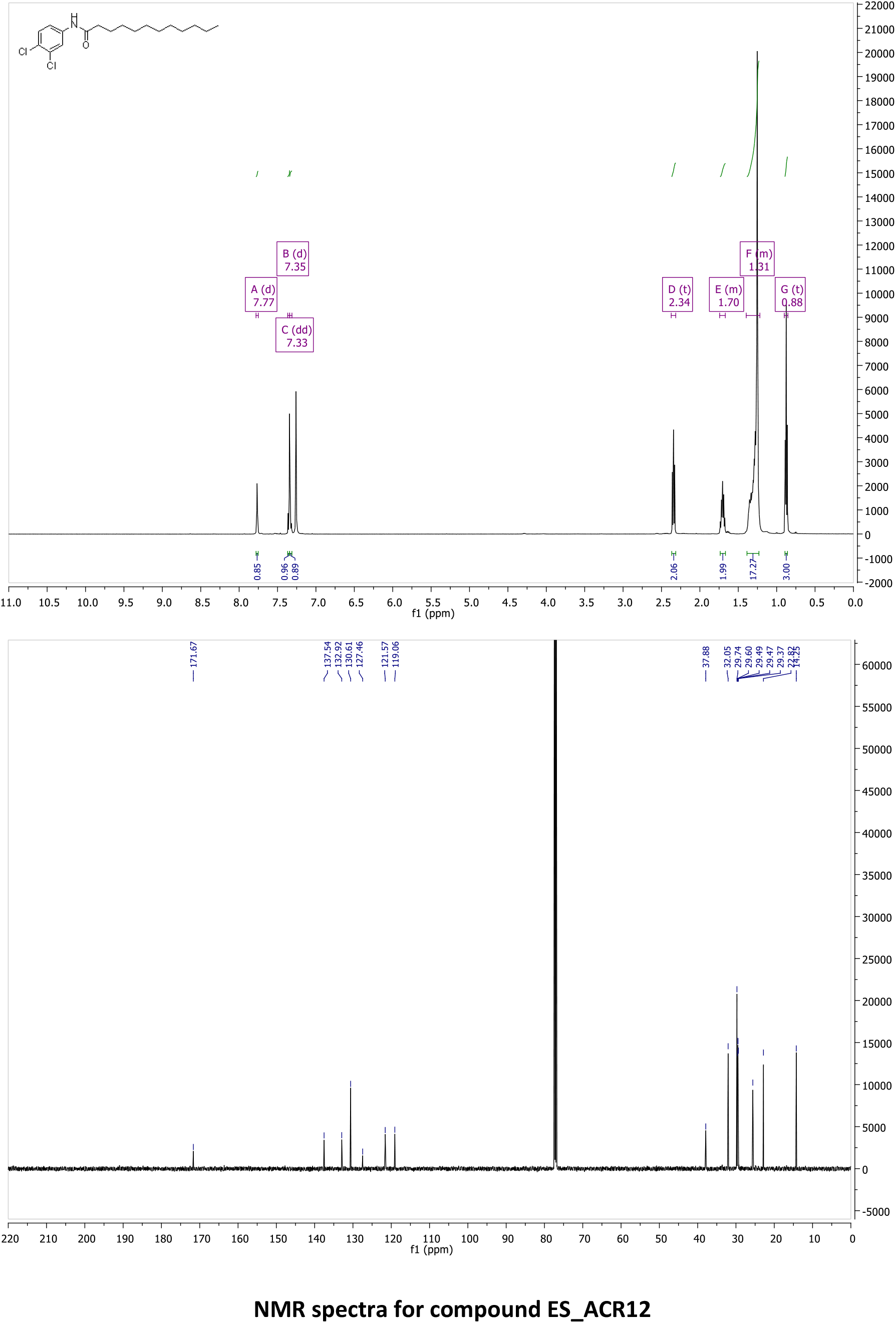
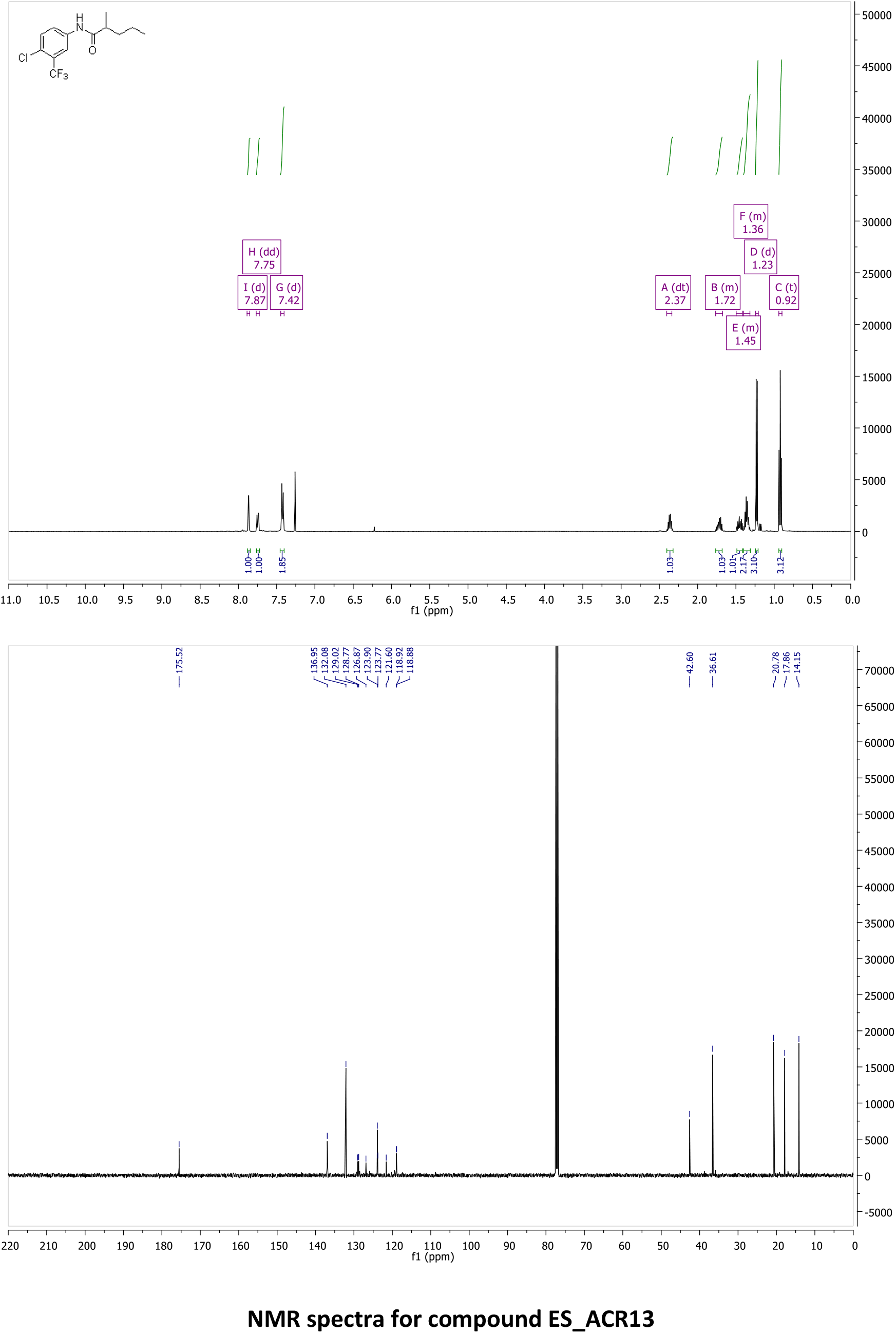
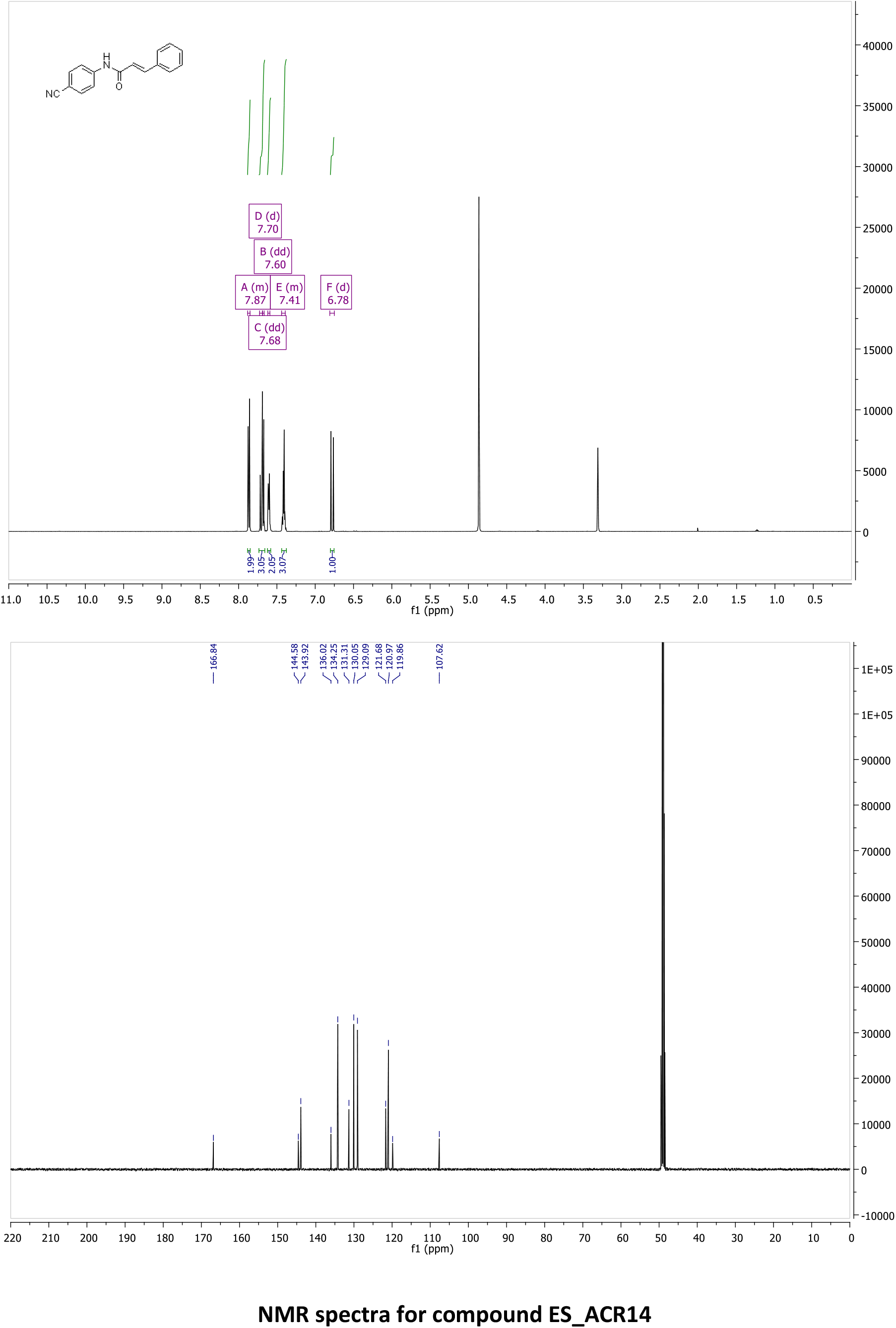
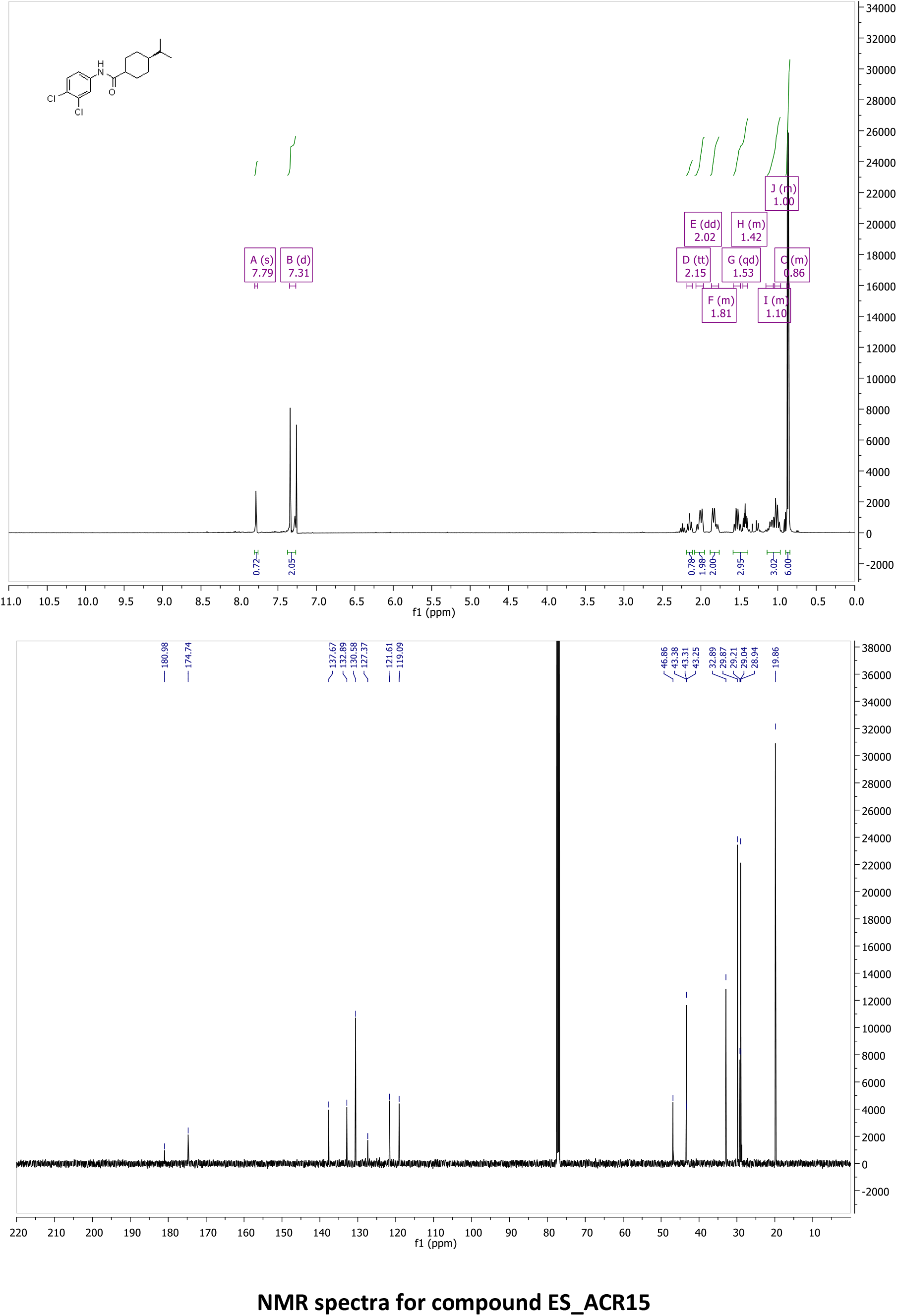
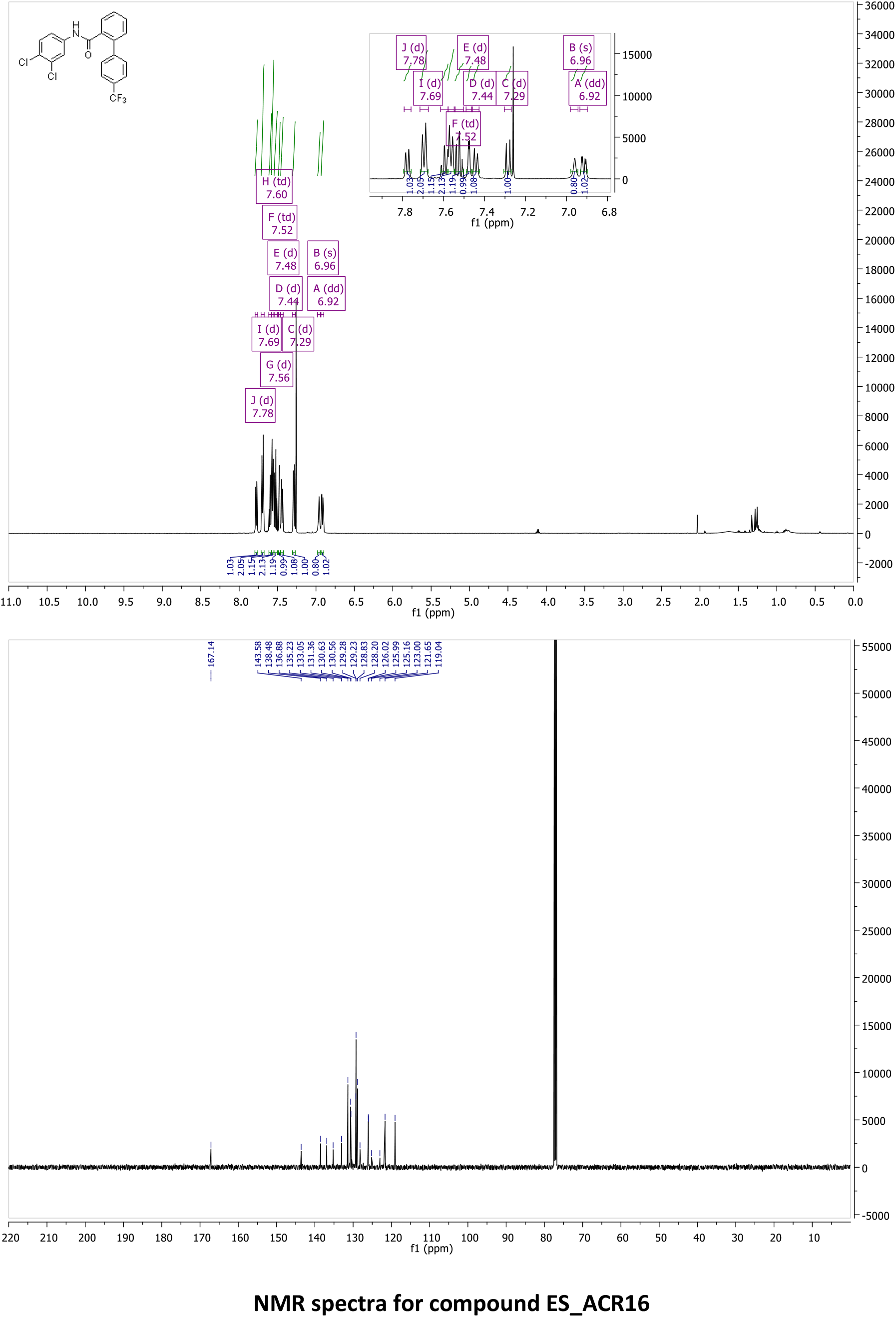
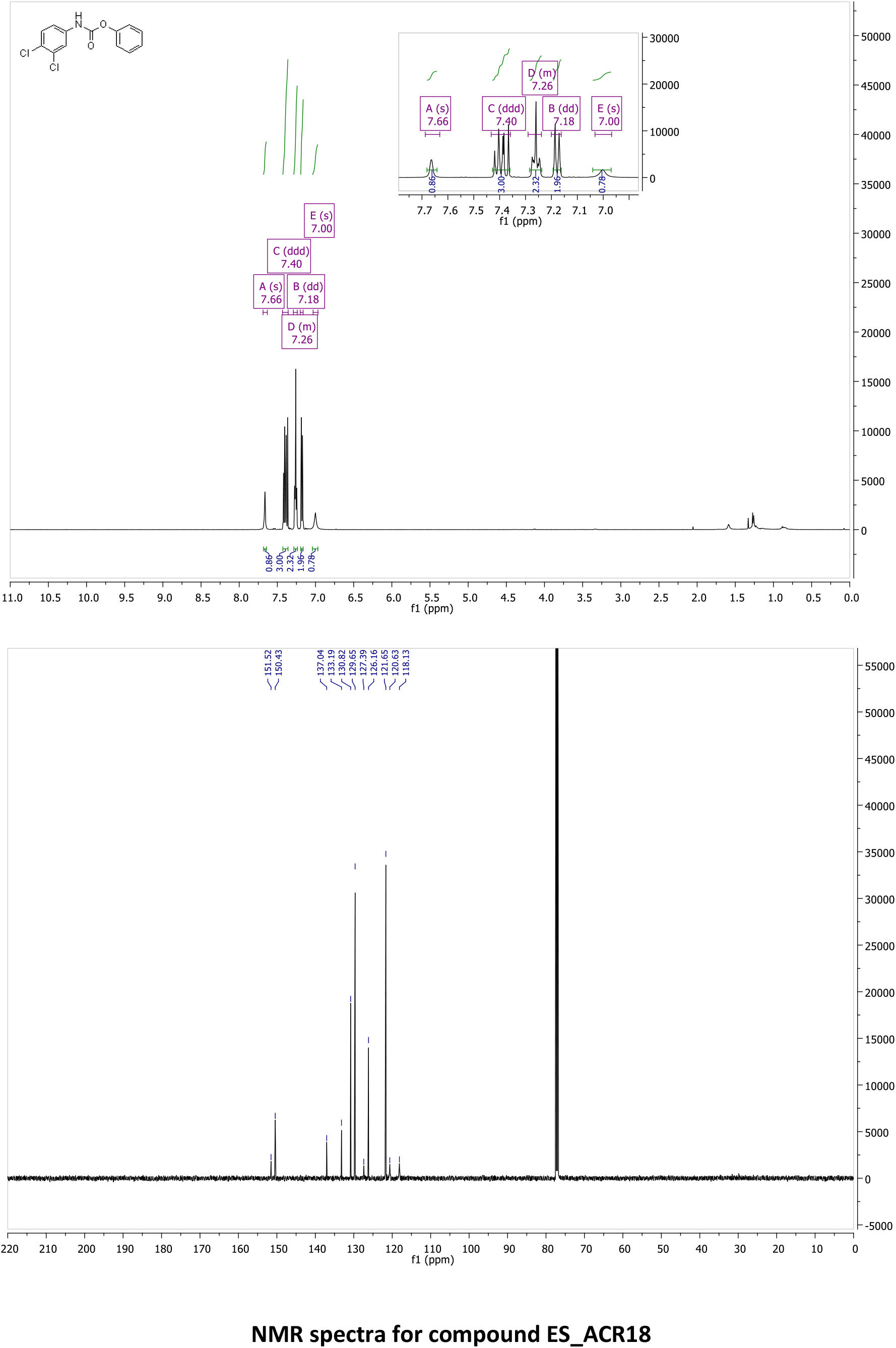
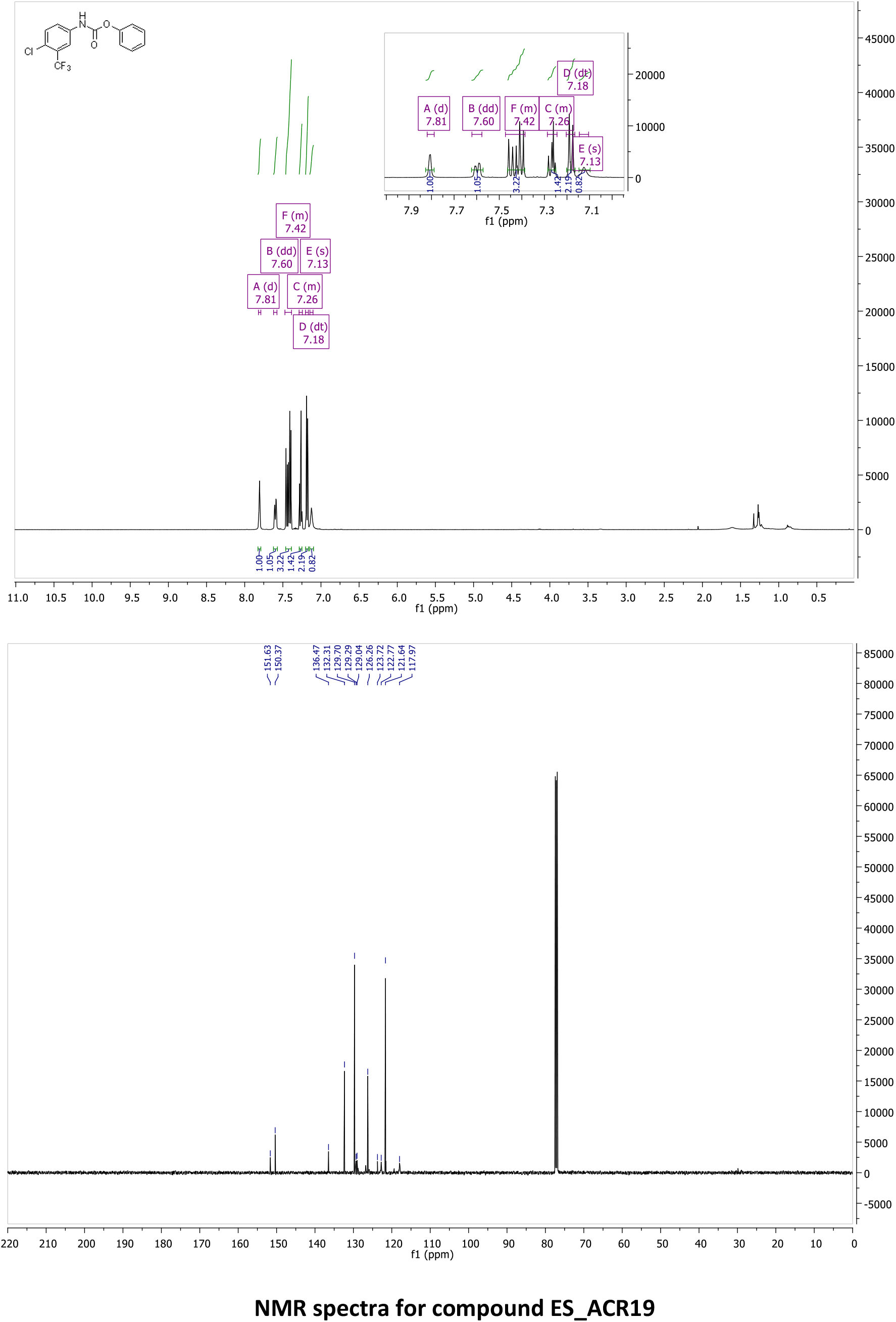
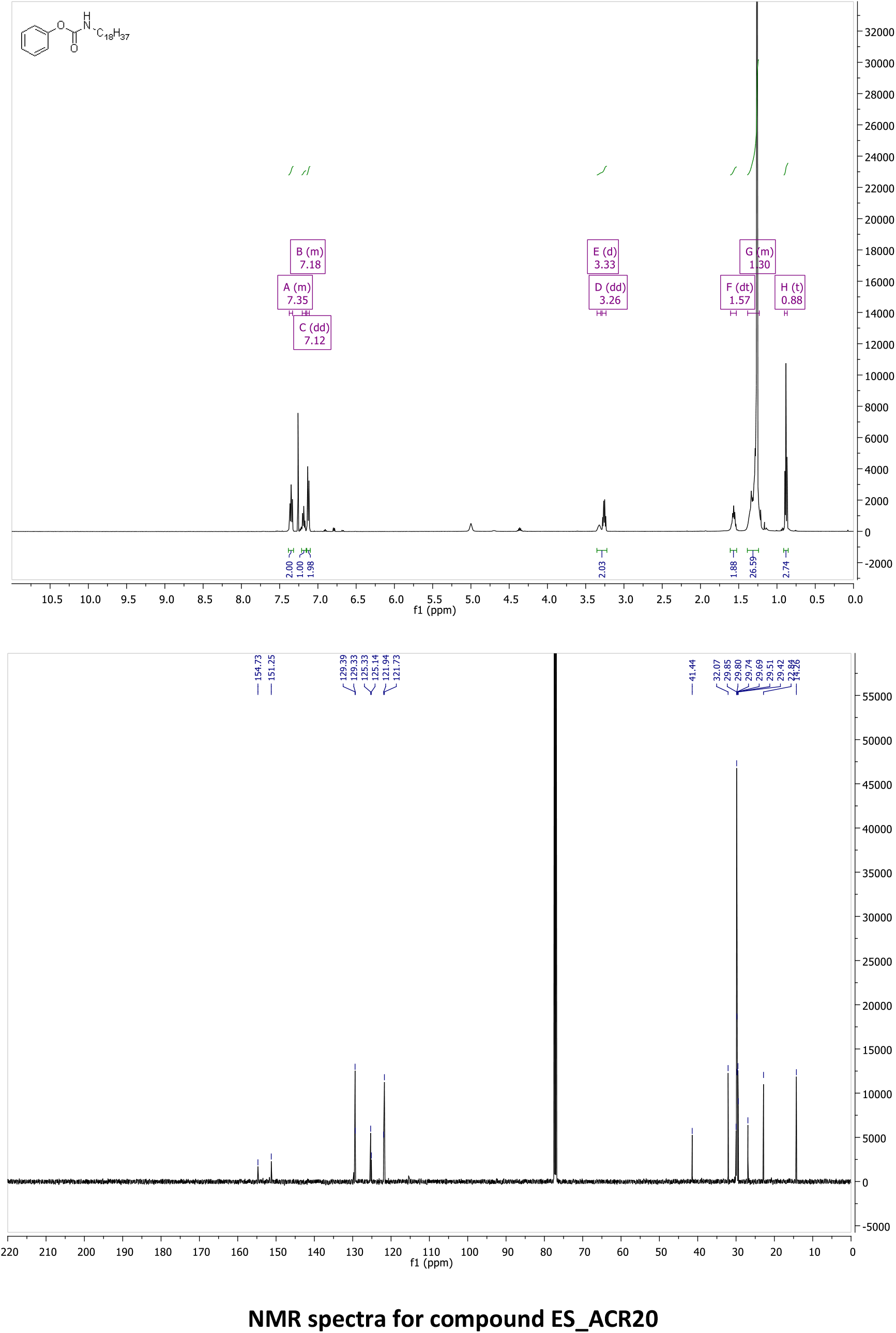
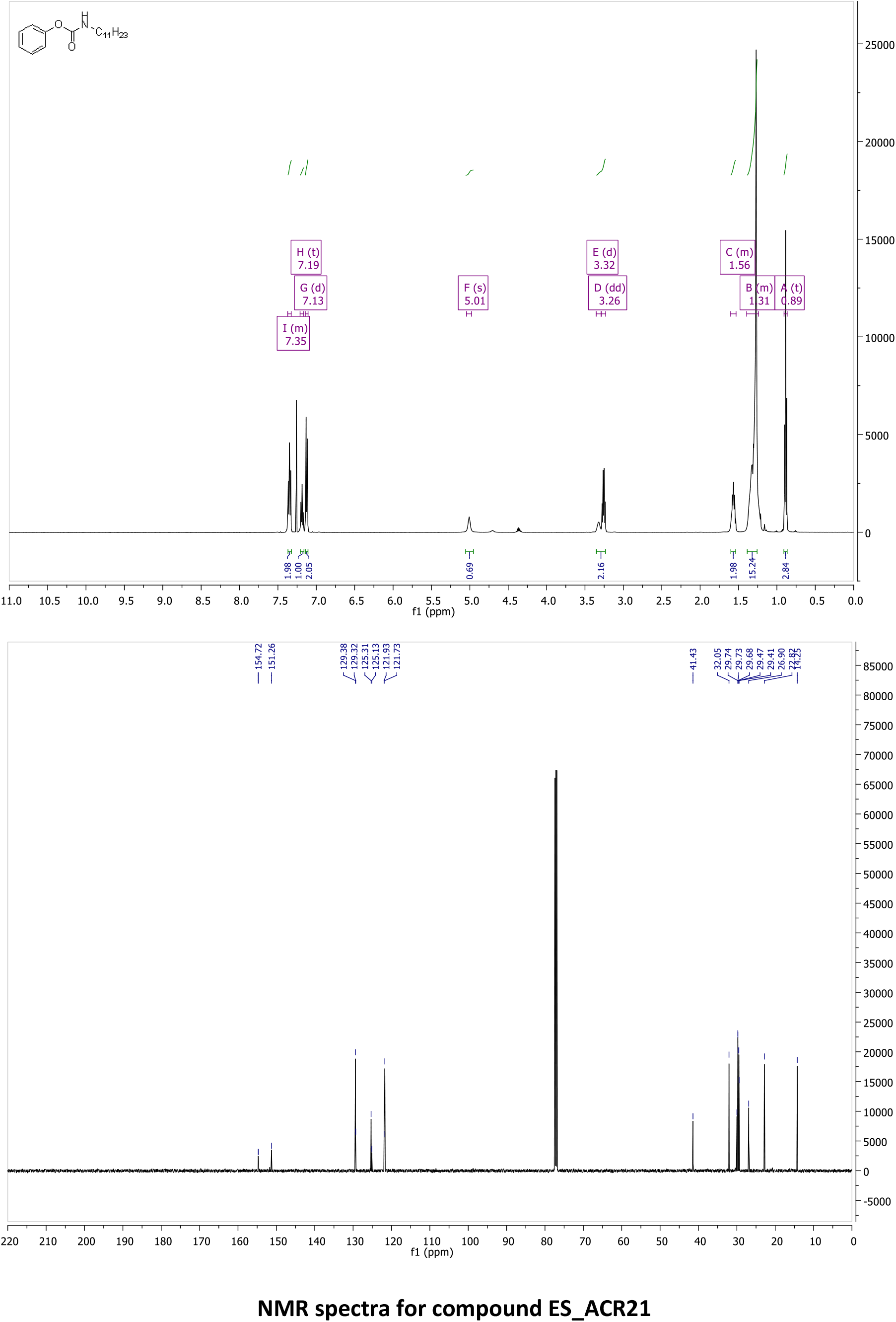
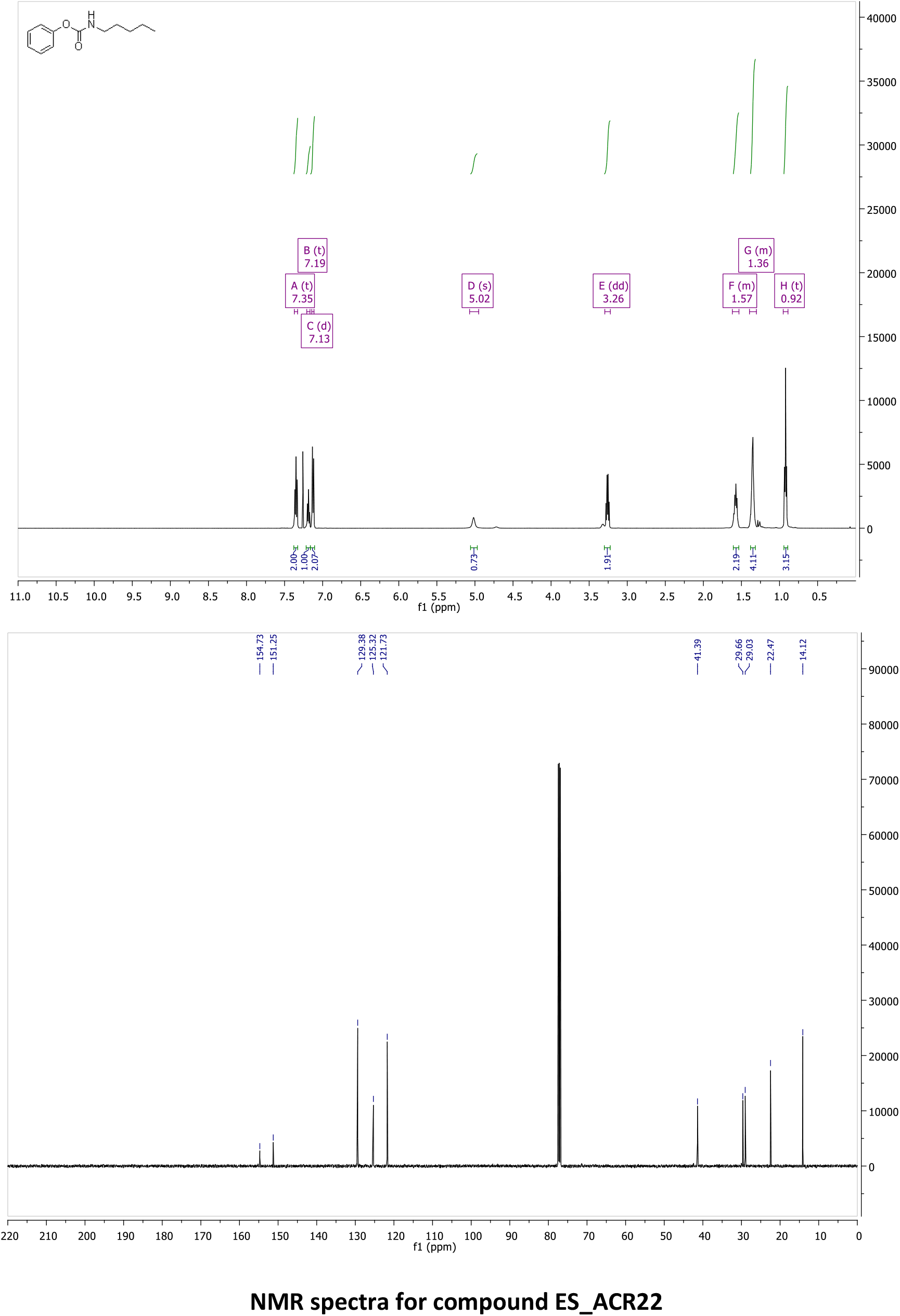
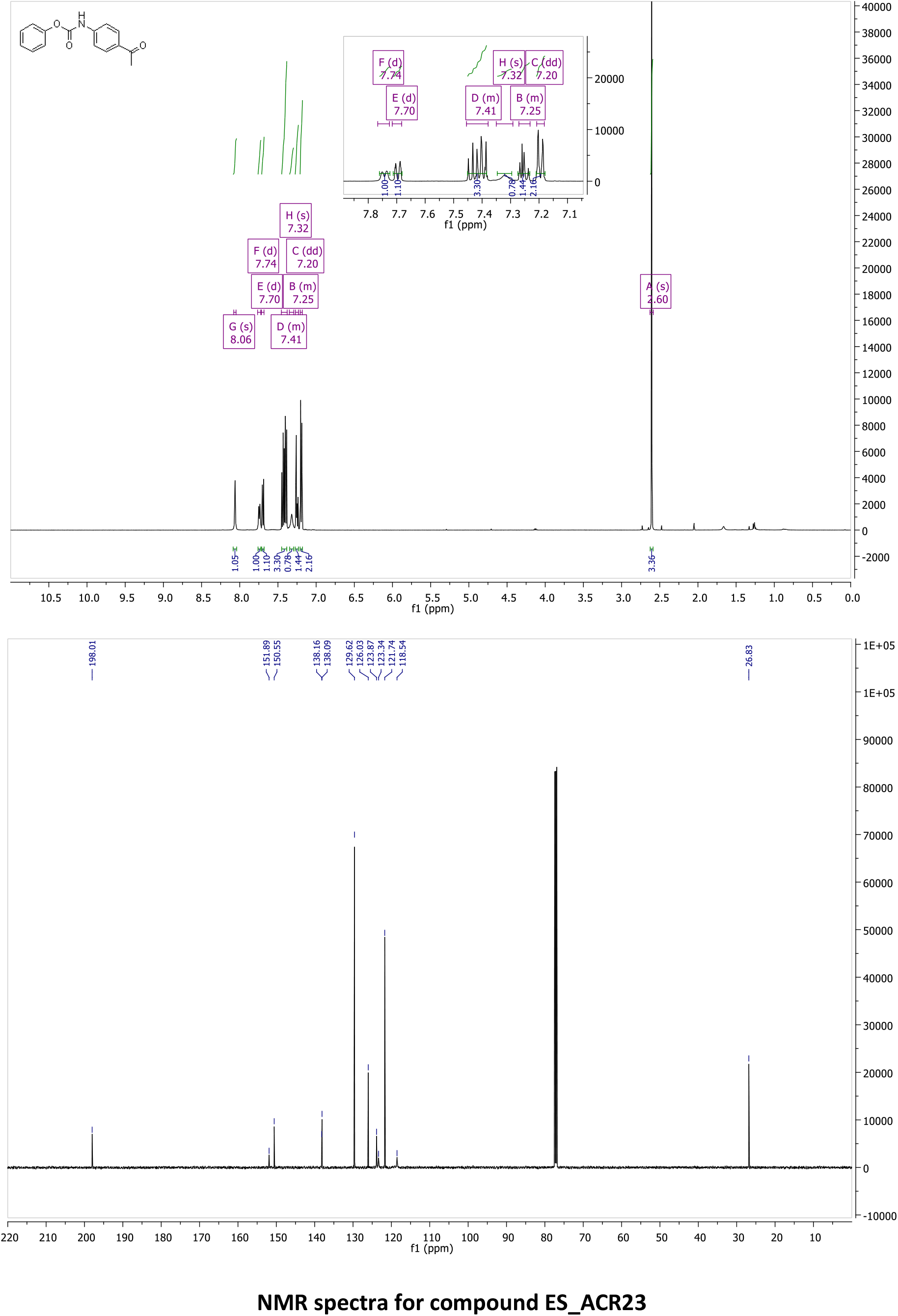
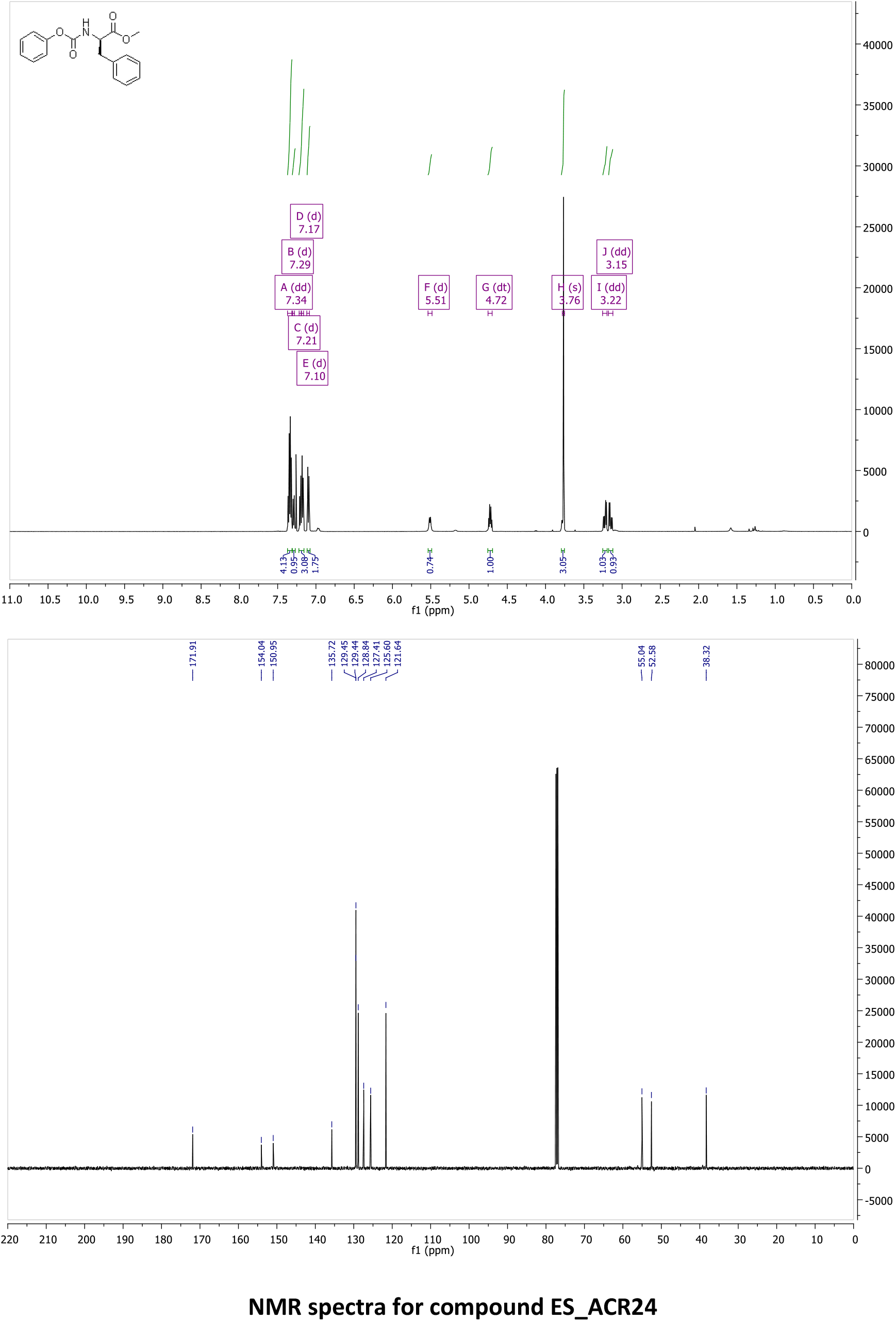
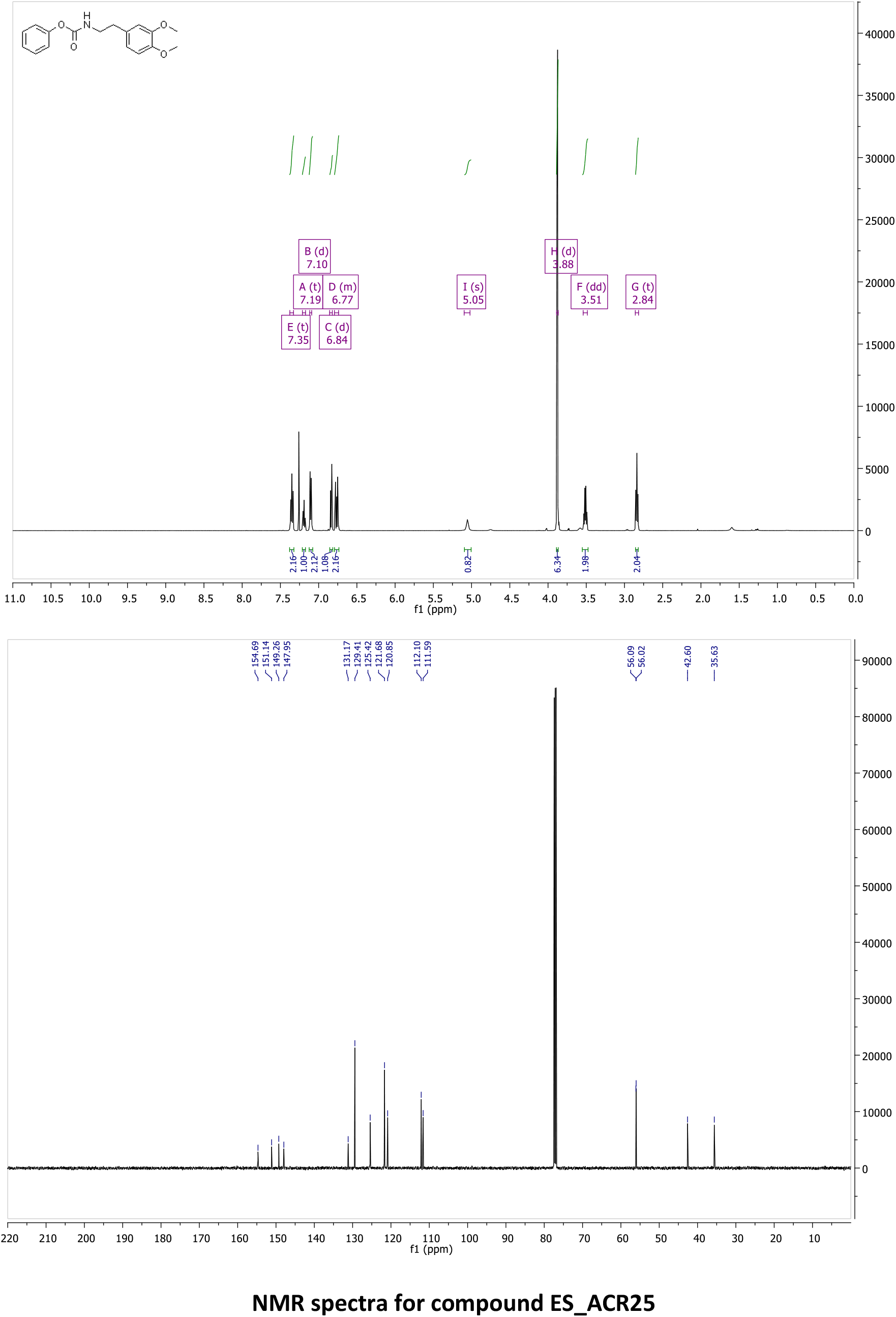
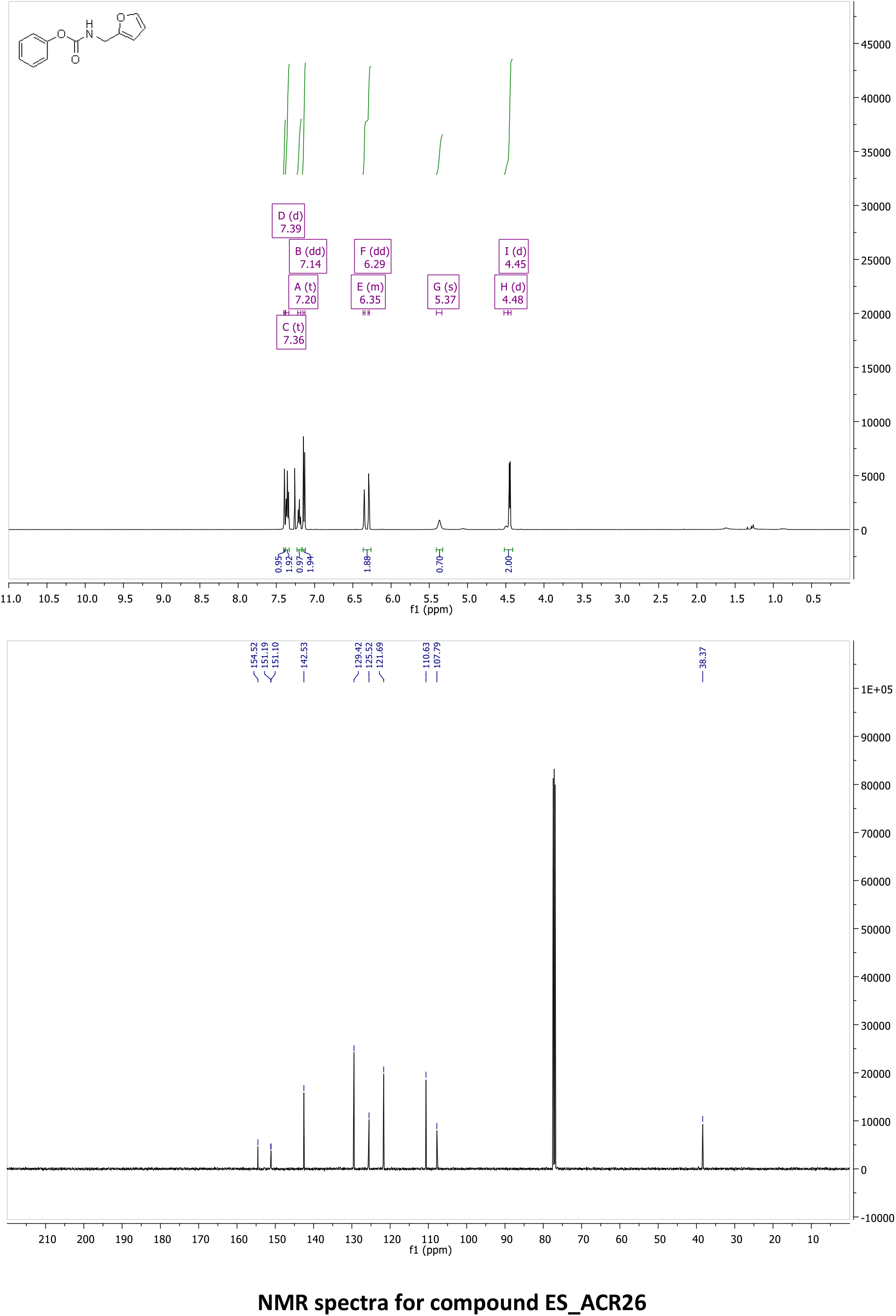
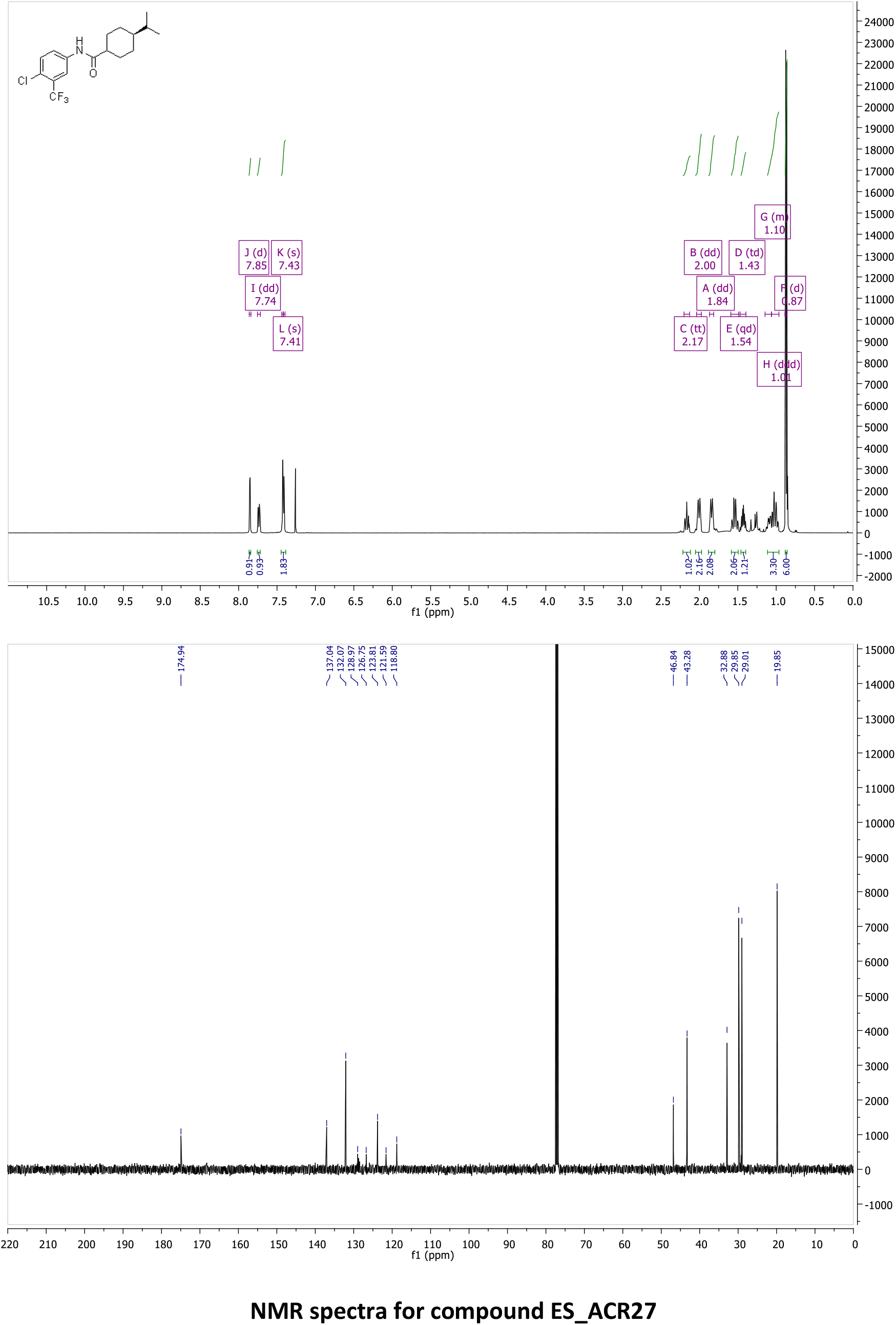

## References

1. Chaurasia, B. & Summers, S. A. Ceramides in Metabolism: Key Lipotoxic Players. Annu. Rev. Physiol. 83, 1–28 (2021).

2. Hannun, Y. A. & Obeid, L. M. Sphingolipids and their metabolism in physiology and disease. Nat. Rev. Mol. Cell Biol. 19, 175–191 (2017).

3. Futerman, A. H. Ceramide turnover (version 2019.4) in the IUPHAR/BPS Guide to Pharmacology Database. IUPHAR/BPS Guid. to Pharmacol. CITE 2019, (2019).

4. Summers, S. A., Chaurasia, B. & Holland, W. L. Metabolic Messengers: ceramides. Nat. Metab. 1, 1051–1058 (2019).

5. Lewis, A. C., Wallington-Beddoe, C. T., Powell, J. A. & Pitson, S. M. Targeting sphingolipid metabolism as an approach for combination therapies in haematological malignancies. Cell Death Discov. 4, 72 (2018).

6. Duarte, C. et al. Elusive Roles of the Different Ceramidases in Human Health, Pathophysiology, and Tissue Regeneration. Cells 9, 1–20 (2020).

7. Ogretmen, B. Sphingolipid metabolism in cancer signalling and therapy. Nat. Rev. Cancer 18, 33–50 (2018).

8. Summers, S. A. Could Ceramides Become the New Cholesterol? Cell Metab. 27, 276–280 (2018).

9. Morad, S. A. F. & Cabot, M. C. Ceramide-orchestrated signalling in cancer cells. Nat. Rev. Cancer 13, 51–65 (2013).

10. Vykoukal, J. et al. Caveolin-1-mediated sphingolipid oncometabolism underlies a metabolic vulnerability of prostate cancer. Nat. Commun. 11, (2020).

11. Hannun, Y. A. & Obeid, L. M. Principles of bioactive lipid signalling: Lessons from sphingolipids. Nat. Rev. Mol. Cell Biol. 9, 139–150 (2008).

12. Pant, D. C. et al. Loss of the sphingolipid desaturase DEGS1 causes hypomyelinating leukodystrophy. J. Clin. Invest. 129, 1240–1256 (2019).

13. Karsai, G. et al. DEGS1-associated aberrant sphingolipid metabolism impairs nervous system function in humans. J. Clin. Invest. 129, 1229–1239 (2019).

14. Vasiliauskaité-Brooks, I. et al. Structural insights into adiponectin receptors suggest ceramidase activity. Nature 544, 120–123 (2017).

15. Pei, J., Millay, D. P., Olson, E. N. & Grishin, N. V. CREST - a large and diverse superfamily of putative transmembrane hydrolases. Biol. Direct 6, 37 (2011).

16. Parveen, F. et al. Role of Ceramidases in Sphingolipid Metabolism and Human Diseases. Cells 8, 1573 (2019).

17. Dementiev, A. et al. Molecular Mechanism of Inhibition of Acid Ceramidase by Carmofur. J. Med. Chem. 62, 987–992 (2019).

18. Edvardson, S. et al. Deficiency of the alkaline ceramidase ACER3 manifests in early childhood by progressive leukodystrophy. J. Med. Genet. 53, 389–396 (2016).

19. Vasiliauskaité-Brooks, I. et al. Structure of a human intramembrane ceramidase explains enzymatic dysfunction found in leukodystrophy. Nat. Commun. 9, 5437 (2018).

20. Vasiliauskaité-Brooks, I., Healey, R. D. & Granier, S. 7TM proteins are not necessarily GPCRs. Mol. Cell. Endocrinol. 491, 110397 (2019).

21. Chen, C. et al. ACER3 supports development of acute myeloid leukemia. Biochem. Biophys. Res. Commun. 478, 33–38 (2016).

22. Yin, Y., Xu, M., Gao, J. & Li, M. Alkaline ceramidase 3 promotes growth of hepatocellular carcinoma cells via regulating S1P/S1PR2/PI3K/AKT signaling. Pathol. Res. Pract. 214, 1381–1387 (2018).

23. Wang, K. et al. Targeting alkaline ceramidase 3 alleviates the severity of nonalcoholic steatohepatitis by reducing oxidative stress. Cell Death Dis. 11, 28 (2020).

24. Bhabak, K. P. et al. Development of a novel FRET probe for the real-time determination of ceramidase activity. ChemBioChem 14, 1049–1052 (2013).

25. Saied, E. M., Banhart, S., Bürkle, S. E., Heuer, D. & Arenz, C. A series of ceramide analogs modified at the 1-position with potent activity against the intracellular growth of Chlamydia trachomatis. Future Med. Chem. 7, 1971–1980 (2015).

26. Mohamed, Z. H., Rhein, C., Saied, E. M., Kornhuber, J. & Arenz, C. FRET probes for measuring sphingolipid metabolizing enzyme activity. Chem. Phys. Lipids 216, 152–161 (2018).

27. Pinkert, T., Furkert, D., Korte, T., Herrmann, A. & Arenz, C. Amplification of a FRET Probe by Lipid-Water Partition for the Detection of Acid Sphingomyelinase in Live Cells. Angew. Chemie Int. Ed. 56, 2790–2794 (2017).

28. Hu, W. et al. Alkaline ceramidase 3 (ACER3) hydrolyzes unsaturated long-chain ceramides, and its down-regulation inhibits both cell proliferation and apoptosis. J. Biol. Chem. 285, 7964–7976 (2010).

29. Xu, R. Golgi alkaline ceramidase regulates cell proliferation and survival by controlling levels of sphingosine and S1P. FASEB J. 20, 1813–1825 (2006).

30. Remacle, A. G. et al. Novel MT1-MMP Small-Molecule Inhibitors Based on Insights into Hemopexin Domain Function in Tumor Growth. Cancer Res. 72, 2339–2349 (2012).

31. Pekkonen, P. et al. Lymphatic endothelium stimulates melanoma metastasis and invasion via MMP14-dependent Notch3 and β1-integrin activation. Elife 7, 1–28 (2018).

32. Clancy, J. W. et al. Regulated delivery of molecular cargo to invasive tumour-derived microvesicles. Nat. Commun. 6, 1–11 (2015).

33. Konermann, L., Pan, J. & Liu, Y.-H. Hydrogen exchange mass spectrometry for studying protein structure and dynamics. Chem. Soc. Rev. 40, 1224–1234 (2011).

34. Zheng, J., Strutzenberg, T., Pascal, B. D. & Griffin, P. R. Protein dynamics and conformational changes explored by hydrogen/deuterium exchange mass spectrometry. Curr. Opin. Struct. Biol. 58, 305–313 (2019).

35. Masson, G. R. et al. Recommendations for performing, interpreting and reporting hydrogen deuterium exchange mass spectrometry (HDX-MS) experiments. Nat. Methods 16, 595–602 (2019).

36. Wang, L., Friesner, R. A. & Berne, B. J. Replica Exchange with Solute Scaling: A More Efficient Version of Replica Exchange with Solute Tempering (REST2). J. Phys. Chem. B 115, 9431–9438 (2011).

37. Park, T. S., Rosebury, W., Kindt, E. K., Kowala, M. C. & Panek, R. L. Serine palmitoyltransferase inhibitor myriocin induces the regression of atherosclerotic plaques in hyperlipidemic ApoE-deficient mice. Pharmacol. Res. 58, 45–51 (2008).

38. Holland, W. L. et al. Inhibition of Ceramide Synthesis Ameliorates Glucocorticoid-, Saturated-Fat-, and Obesity-Induced Insulin Resistance. Cell Metab. 5, 167–179 (2007).

39. Ussher, J. R. et al. Inhibition of de novo ceramide synthesis reverses diet-induced insulin resistance and enhances whole-body oxygen consumption. Diabetes 59, 2453–2464 (2010).

40. Zhang, Q. J. et al. Ceramide mediates vascular dysfunction in diet-induced obesity by PP2A-mediated dephosphorylation of the eNOS-Akt complex. Diabetes 61, 1848–1859 (2012).

41. Ji, R. et al. Increased de novo ceramide synthesis and accumulation in failing myocardium. JCI insight 2, (2017).

42. Park, T. S. et al. Ceramide is a cardiotoxin in lipotoxic cardiomyopathy. J. Lipid Res. 49, 2101–2112 (2008).

43. Chaurasia, B. et al. Targeting a ceramide double bond improves insulin resistance and hepatic steatosis. Science (80-.). 365, 386–392 (2019).

44. Bielsa, N. et al. Discovery of deoxyceramide analogs as highly selective ACER3 inhibitors in live cells. Eur. J. Med. Chem. 216, 113296 (2021).

45. Lightwood, D. J. et al. A conformation-selective monoclonal antibody against a small molecule-stabilised signalling-deficient form of TNF. Nat. Commun. 12, (2021).

46. Doan, N. B. et al. Acid ceramidase and its inhibitors: A de novo drug target and a new class of drugs for killing glioblastoma cancer stem cells with high efficiency. Oncotarget 8, 112662–112674 (2017).

47. Lai, M. et al. Complete Acid Ceramidase ablation prevents cancer-initiating cell formation in melanoma cells. Sci. Rep. 7, 1–14 (2017).

48. Bai, A. et al. Dose dependent actions of LCL521 on acid ceramidase and key sphingolipid metabolites. Bioorg. Med. Chem. 26, 6067–6075 (2018).

49. Zhang, S. et al. TIMELESS regulates sphingolipid metabolism and tumor cell growth through Sp1/ACER2/S1P axis in ER-positive breast cancer. Cell Death Dis. 11, (2020).

50. Elegheert, J. et al. Lentiviral transduction of mammalian cells for fast, scalable and high-level production of soluble and membrane proteins. Nat. Protoc. 13, 2991–3017 (2018).

51. Lau, A. M., Claesen, J., Hansen, K. & Politis, A. Deuteros 2.0: peptide-level significance testing of data from hydrogen deuterium exchange mass spectrometry. Bioinformatics 37, 270–272 (2021).

52. Gordon, J. C. et al. H++: A server for estimating pKas and adding missing hydrogens to macromolecules. Nucleic Acids Res. 33, 368–371 (2005).

53. Wang, J., Cieplak, P. & Kollman, P. A. How well does a restrained electrostatic potential (RESP) model perform in calculating conformational energies of organic and biological molecules? J. Comput. Chem. 21, 1049–1074 (2000).

54. Frisch, M. J. et al. Gaussian 09. (2016).

55. Li, P. & Merz, K. M. MCPB.py: A Python Based Metal Center Parameter Builder. J. Chem. Inf. Model. 56, 599–604 (2016).

56. Van Der Spoel, D. et al. GROMACS: Fast, flexible, and free. J. Comput. Chem. 26, 1701–1718 (2005).

57. Tribello, G. A., Bonomi, M., Branduardi, D., Camilloni, C. & Bussi, G. PLUMED 2: New feathers for an old bird. Comput. Phys. Commun. 185, 604–613 (2014).

58. Salentin, S., Schreiber, S., Haupt, V. J., Adasme, M. F. & Schroeder, M. PLIP: Fully automated protein-ligand interaction profiler. Nucleic Acids Res. 43, W443–W447 (2015).

59. Airola, M. V. et al. Structural Basis for Ceramide Recognition and Hydrolysis by Human Neutral Ceramidase. Structure 23, 1482–1491 (2015).

60. Gebai, A., Gorelik, A., Li, Z., Illes, K. & Nagar, B. Structural basis for the activation of acid ceramidase. Nat. Commun. 9, 1621 (2018).

## References

1. Saied, E. M., Banhart, S., Bürkle, S. E., Heuer, D. & Arenz, C. A series of ceramide analogs modified at the 1-position with potent activity against the intracellular growth of *Chlamydia trachomatis*. Future Medicinal Chemistry 7, 1971–1980 (2015).

2. Samaha, D. et al. Liposomal FRET Assay Identifies Potent Drug-Like Inhibitors of the Ceramide Transport Protein (CERT). Chemistry 26, 16616–16621 (2020).

3. Bhabak, K. P. et al. Development of a Novel FRET Probe for the Real-Time Determination of Ceramidase Activity. ChemBioChem 14, 1049–1052 (2013).

4. Wessig, P. et al. Two-photon FRET pairs based on coumarin and DBD dyes. RSC Adv. 6, 33510–33513 (2016).

5. Wenskowsky, L. et al. Identification and Characterization of a Single High-Affinity Fatty Acid Binding Site in Human Serum Albumin. Angewandte Chemie International Edition 57, 1044–1048 (2018).

6. Collenburg, L. et al. The Activity of the Neutral Sphingomyelinase Is Important in T Cell Recruitment and Directional Migration. Front. Immunol. 8, (2017).

